# PalaeoChip Arctic1.0: An optimised eDNA targeted enrichment approach to reconstructing past environments

**DOI:** 10.1101/730440

**Authors:** Tyler J. Murchie, Melanie Kuch, Ana Duggan, Marissa L. Ledger, Kévin Roche, Jennifer Klunk, Emil Karpinski, Dirk Hackenberger, Tara Sadoway, Ross MacPhee, Duane Froese, Hendrik Poinar

**Author notes:** Author contributions: TJM: Research design, wet-lab processing, analysis, figure design, writing MK: Research design, wet-lab processing, manuscript editing AD: Bioinformatic assistance, analysis, manuscript editing MLL: Laboratory assistance, writing, manuscript editing KR: Laboratory assistance JK: Laboratory and research design assistance EK: Research design assistance, initial experiments with 4°C spin, manuscript editing DH: Initial experiments with 4°C spin TS: Metabarcoding and preliminary analyses of cores, core collection, manuscript editing RM: Research design, manuscript editing, collaborative principal investigator DF: Research design, analysis, manuscript editing, core collection, collaborative principal investigator HP: Research design, analysis, manuscript editing, McMaster Ancient DNA Centre principal investigator.

## Abstract

Ancient environmental DNA has been established as a viable biomolecular proxy for tracking taxonomic presence through time in a local environment, even in the total absence of primary tissues. It is thought that sedimentary ancient DNA (sedaDNA) survives through mineral binding. And while these organo-mineral complexes likely facilitate long-term preservation, they also challenge our ability to release and isolate target molecules. Two limitations in sedaDNA extraction impede many palaeoenvironmental reconstructions: the post-extraction carryover of enzymatic inhibitors, and sedaDNA loss when attempting to reduce inhibitor co-elution. Here, we present an optimised eDNA targeted enrichment approach for reconstructing past environments. Our new extraction protocol with targeted enrichment averages a 14.6-fold increase in on-target plant and animal DNA compared to a commercial soil extraction kit, and a 22.6-fold increase compared to a PCR metabarcoding approach. To illustrate the effectiveness of the PalaeoChip Arctic1.0 protocol, we present results of plant and animal presence from permafrost samples and discuss new potential evidence for the late survival (ca. 9685 BP) of mammoth (*Mammuthus sp*.) and horse (*Equus sp*.) in the Klondike Region of Yukon, Canada. This approach translates to a more diverse and sensitive dataset with increased sequencing efficiency of ecologically informative sedaDNA.

## Introduction

Means of recovering and analyzing ecologically informative sedimentary ancient DNA (sedaDNA) have improved substantially thanks to ongoing developments in high-throughput sequencing (HTS) technologies (Taberlet et al., 2018). SedaDNA molecules have been successfully recovered to evaluate the ‘local’ (Parducci et al., 2017, p. 930; Rawlence et al., 2014, p. 616) diachronic presence of animals (Giguet-Covex et al., 2014; Graham et al., 2016; Haile et al., 2009; Pedersen et al., 2016; Slon et al., 2017), plants (Alsos et al., 2015; Anderson-Carpenter et al., 2011; Epp et al., 2015; Niemeyer et al., 2017; Willerslev et al., 2014), fungi (Bellemain et al., 2013), microbiota (Ahmed et al., 2018; D’Costa et al., 2011), and eukaryotic parasites (Søe et al., 2018) from a diverse range of depositional settings. It is thought that much of sedaDNA survives in the absence of primary tissues through the formation of organo-mineral complexes (Arnold et al., 2011; Blum et al., 1997; Gardner and Gunsch, 2017; Greaves and Wilson, 1970; Lorenz and Wackernagel, 1987a, 1987b; Morrissey et al., 2015; Ogram et al., 1988) as extracellular genetic material binds to common constituents of sediments such as humics (Crecchio and Stotzky, 1998), calcite (Cleaves et al., 2011), clays (Cai et al., 2006; Goring and Bartholomew, 1952; Greaves and Wilson, 1969), and silica (Bezanilla et al., 1995; Lorenz and Wackernagel, 1987a). Soil minerals have been found to stabilize a fraction of environmental DNA, allowing those molecules to resist decomposition (Morrissey et al., 2015), but strong mineral binding can also result in marginal sedaDNA release (Alvarez et al., 1998; Saeki et al., 2010). Extracellular mineral-bound sedaDNA is recovered in bulk in the form of disseminated biomolecules from a diverse range of organisms. This fact typically prohibits genomic reconstructions of single individuals, but it can allow for identifying the presence (and to a lesser extent, absence and potentially even relative abundance) of taxa at ecologically informative taxonomic ranks. The method shows the most promise in reconstructing palaeoflora (Anderson-Carpenter et al., 2011; Niemeyer et al., 2017; Pedersen et al., 2016; Sjögren et al., 2016; Willerslev et al., 2014), or the differential presence of a particular taxon through time (Graham et al., 2016). A single library can be used to identify sedaDNA from multiple domains simultaneously with shotgun sequencing or can be targeted to amplify or enrich for specific taxa of interest.

Despite rapid advances in ancient DNA (aDNA) techniques, two extraction related challenges persist that can limit the ability to fully utilize sedimentary genetic archives: 1) the carryover of enzymatic inhibitors with techniques designed to maximize the recovery of aDNA characteristic molecules, and 2) the loss, due to overly vigorous inhibitor removal techniques, of ecologically informative sedaDNA that otherwise might be amenable to adapter ligation or amplification. To some degree PCR metabarcoding can mitigate inhibition through dilutions or additional purifications (McKee et al., 2015), with the addition of reagents such as bovine serum albumin (BSA) (Garland et al., 2010; Kreader, 1996), or with very high polymerase concentrations (Alsos et al., 2015). However, metabarcoding can be vulnerable to differential amplification rates due to variable molecular abundance per taxa, unequal damage, variability in metabarcode amplification efficiency, and PCR conditions (Bellemain et al., 2010; Kanagawa, 2003; Krehenwinkel et al., 2018; Nichols et al., 2018; Sze and Schloss, 2019). These factors compound downstream biases in taxonomic determinations, especially if there was substantial loss of low-abundance molecules during inhibitor removal from taxa with comparatively low biomass turnover (Yoccoz et al., 2012, p. 3651).

This study evaluates various inhibitor removal treatments for their ability to reduce the carryover of enzymatic inhibitors in sedaDNA extracts while maximally retaining endogenous palaeoenvironmental DNA that can successfully undergo library adapter ligation. Our aim is to minimize the need for excessive PCR amplification on purified eluates to mitigate the propagation of stochastic biases. Four previously studied (D’Costa et al., 2011; Mahony, 2015; Sadoway, 2014) open-air Yukon permafrost core exposures (Table 1, Figure 1) were chosen to experimentally optimize sedaDNA extraction (detailed in the supplementary materials). We based these modifications on our in-house lysis solution and high-volume binding buffer silica-spin column extraction method as per Dabney et al. (2013). Thereafter, an optimized protocol was selected to evaluate taxonomic assignments between shotgun sequenced and target enriched datasets with this extraction method, as compared with shotgun and enriched libraries extracted using the DNeasy PowerSoil DNA extraction kit (QIAGEN) following manufacturer specifications. For each of the four core sections, sediments were subsampled and homogenized, then split into three 250 mg replicates for both extraction methods. We also compared this sequence data with previously sequenced PCR metabarcoding data on the same core sections. Our sedaDNA modified Dabney et al. (2013) extraction protocol is described here (Figure 2, SET-E). Experimentation with various inhibition removal techniques are detailed in the supplementary materials as SET-A through SET-D.

**Table 1.**
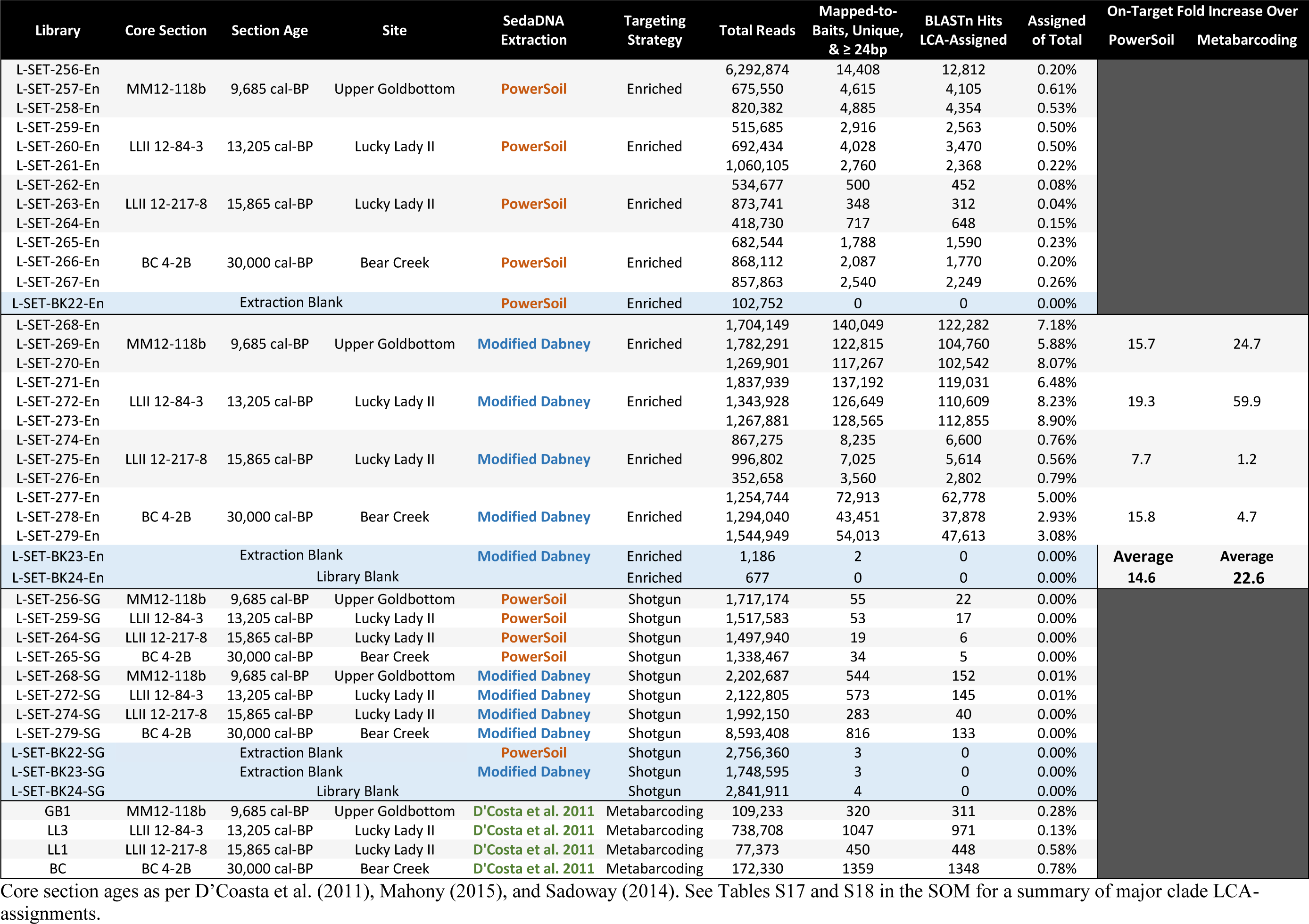
Sample IDs for experiment SET-E with read filtering summaries.

**Figure 1.**
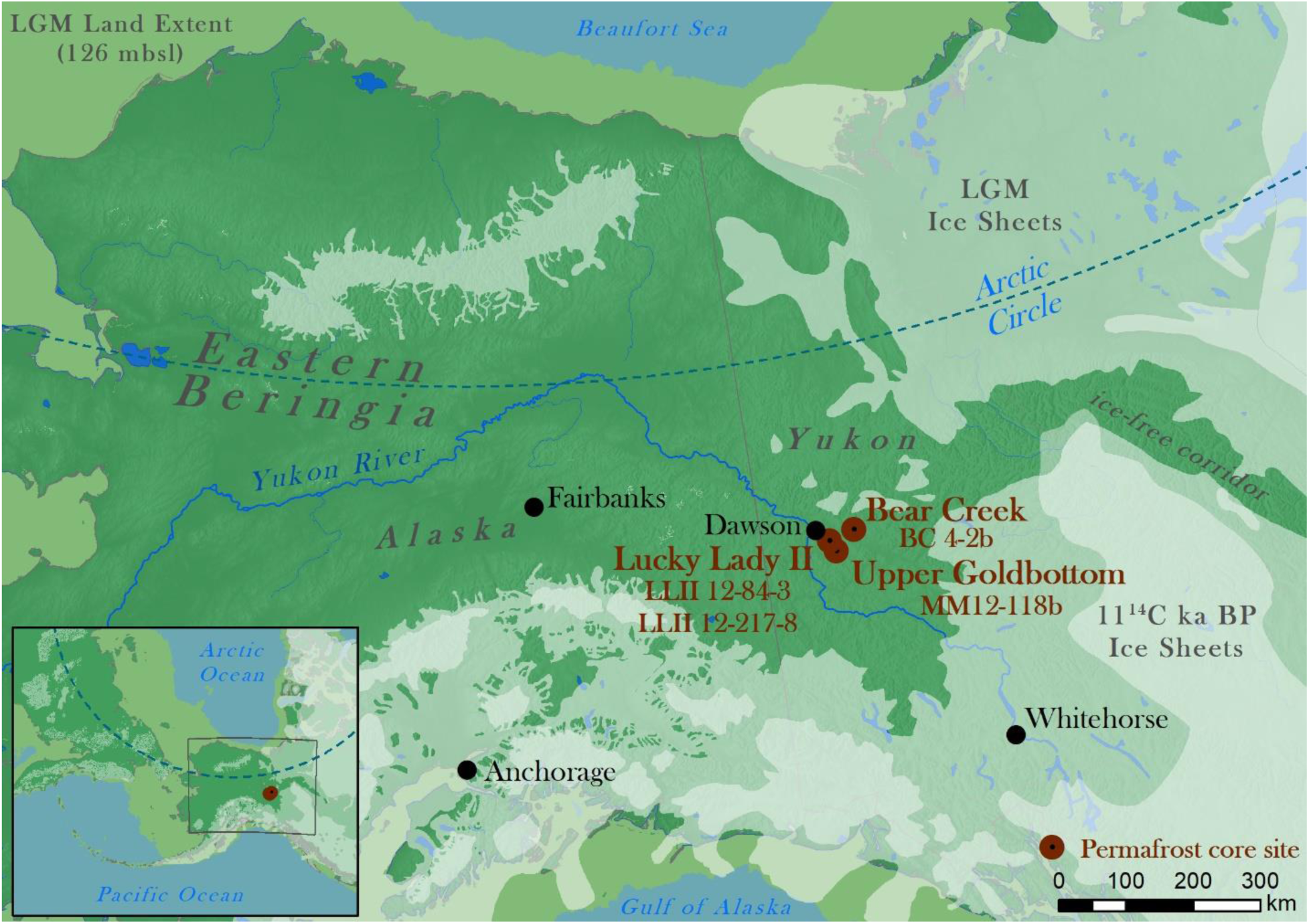
Permafrost coring sites in the Klondike of Yukon, Canada. Ice sheet data from Dyke (2004) and Ehlers et al. (2011). Sea level at Last Glacial Maximum (LGM, 26.5–19 ky BP) (Clark, 2009) set to 126 meters below sea level (msbl) based on midpoint between maximum and minimum eustatic sea level estimation models in Clark and Mix (2002).

**Figure 2.**
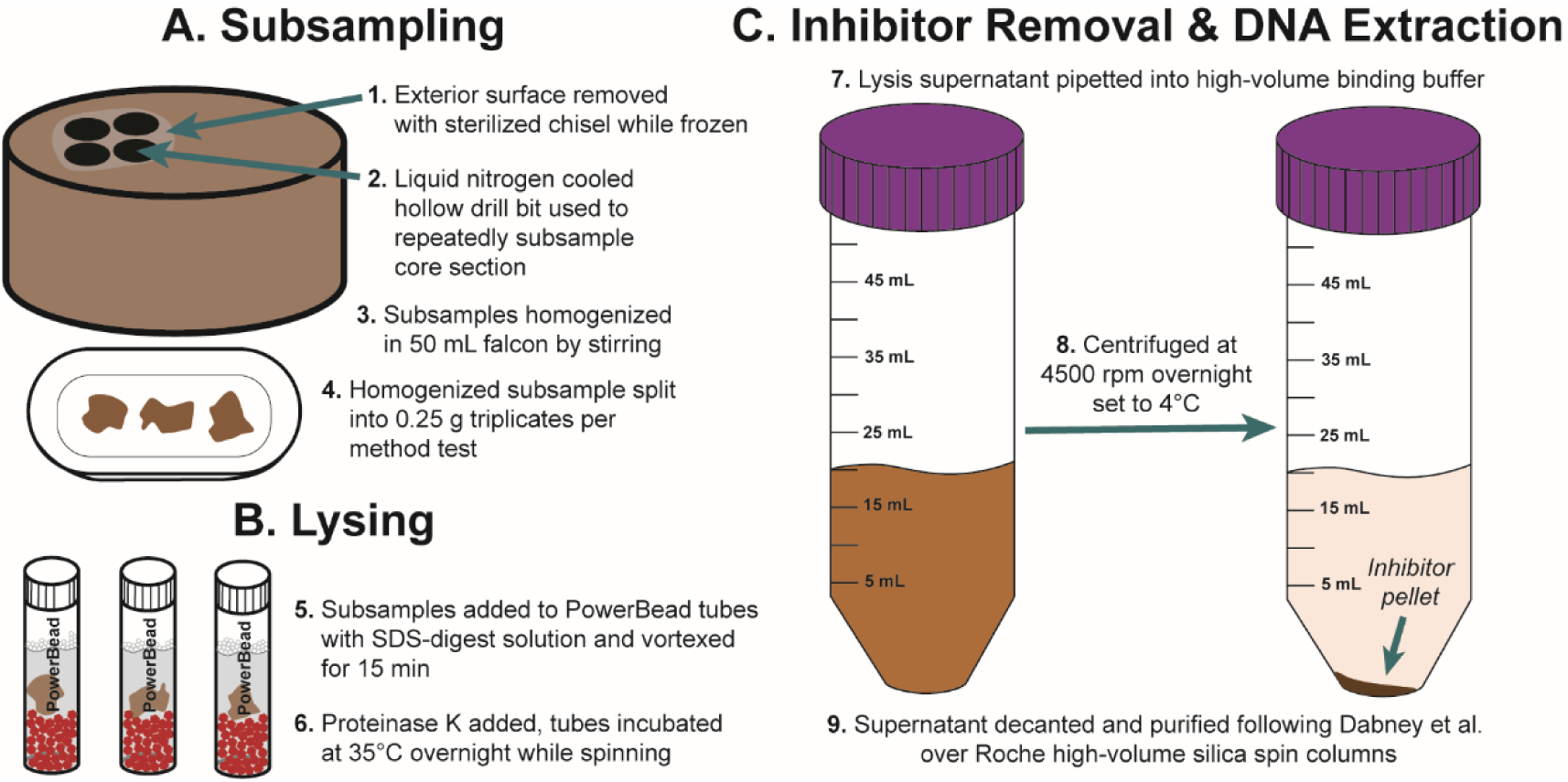
SedaDNA modified Dabney et al. (2013) extraction workflow. See the methods section for further details on extraction, double-stranded library preparation, capture enrichment, qPCR assays, and the bioinformatic workflow.

## Results

Quantitative PCRs (qPCR) on the adapted libraries show an up to 7.0-fold increase in total adapted DNA among the four core samples (average 3.6-fold increase) with our sedaDNA modified Dabney et al. (2013) extraction protocol, and an up to 5.6-fold increase in ‘endogenous’ *trnL* library adapted chloroplast DNA (average 3.0-fold increase) (see Figure 3). Inhibition indices for our sedaDNA modified Dabney extractions were lower than PowerSoil (average = 0.75 versus 0.95 for PowerSoil, see methods section 4 and Figure E14 for a description of the ‘inhibition index’), but this low-level constituent of latent polymerase inhibitors did not impede enzymes for adapter ligation as these samples quantify much higher than PowerSoil extracts post-library prep. Our ongoing experiments with a diverse range of other sediments suggest that extracts with inhibition indices over ∼0.3 are still amenable to library preparation, although potentially with reduced adapter ligation efficiency (see section SET-D in the supplementary online materials for a discussion of extract qPCR inhibition).

**Figure 3.**
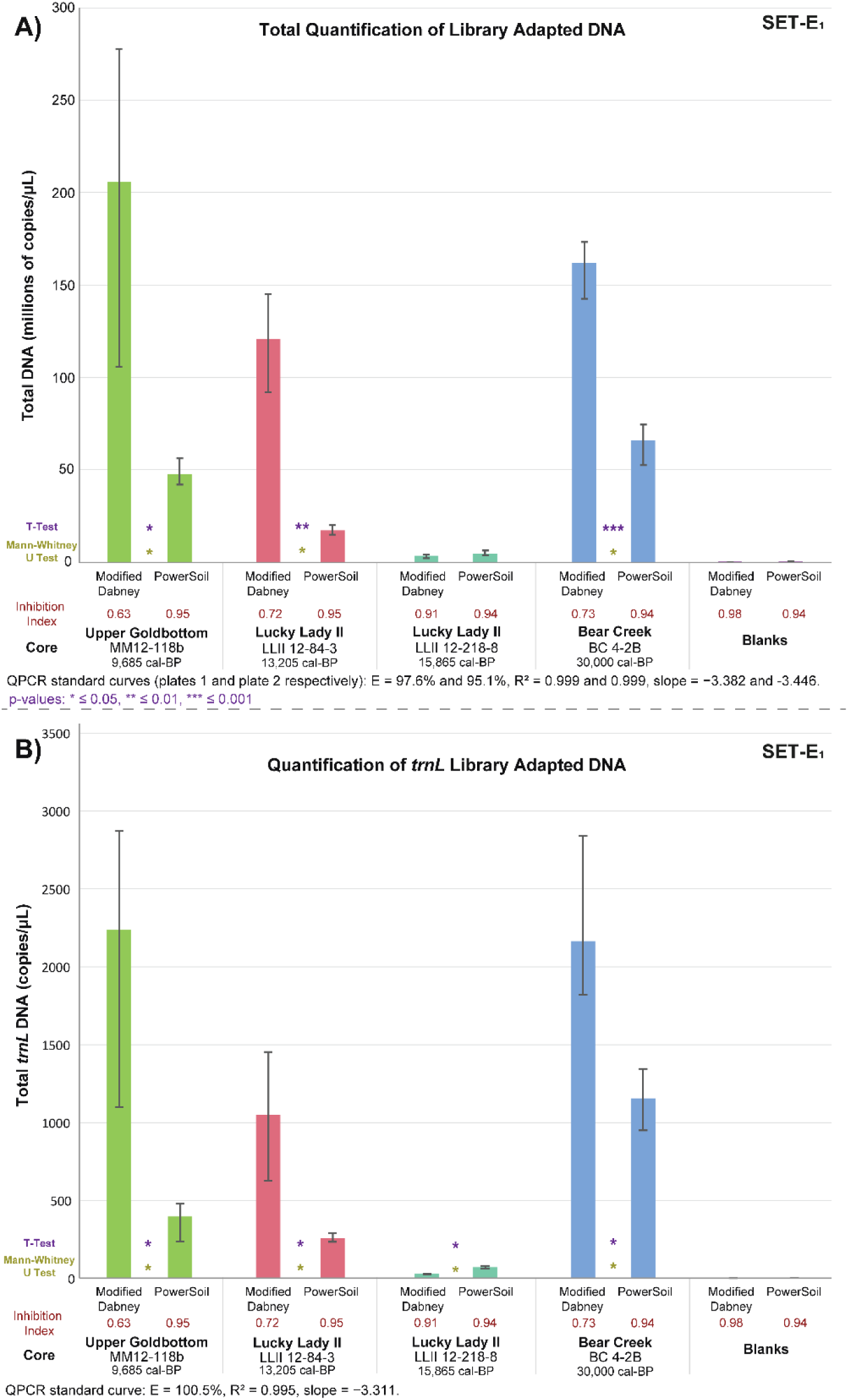
Comparative qPCRs of library adapted DNA. Each bar represents the average DNA copy number of the extraction-core triplicate, with the error bars indicating the maximum and minimum range. DNA concentration for each SET sample was averaged across PCR triplicates. Inhibition index refers to inhibition spike test (Table S7) on extracts prior to library prep. **A)** Total DNA quantification comparing both extraction methods by core; see Table S8 for PCR specifications. **B)** Total *trnL* adapted DNA; see Table S9 for PCR specifications. P-values calculated with both a two-sample t-test (parametric) and Mann-Whitney U test (non-parametric) (two-sided). The large range for modified Dabney extraction core MM12-118B is driven by a single lower copy number extraction replicate. Core LLII 12-217-8 consistently has low DNA recovery, but also a low co-elution of inhibition.

Mapped, *BLASTn* aligned, and LCA-assigned reads extracted with our modified Dabney protocol show an average 14.6-fold increase in bait on-target, map filtered reads over PowerSoil extractions, and a 22.6-fold increase in map-filtered reads over samples targeted with PCR metabarcoding (Table 1). This large fold increase in on-target molecules of total reads translates to a broader range of taxa identified, and a higher proportion of ecologically informative reads sequenced overall (Figures 4–7). Taxa with sufficiently high LCA-assigned read counts also show characteristic aDNA deamination patterns and fragment length distributions with *mapDamage* (Table E2; Figures E8–E12).

**Figure 4.**
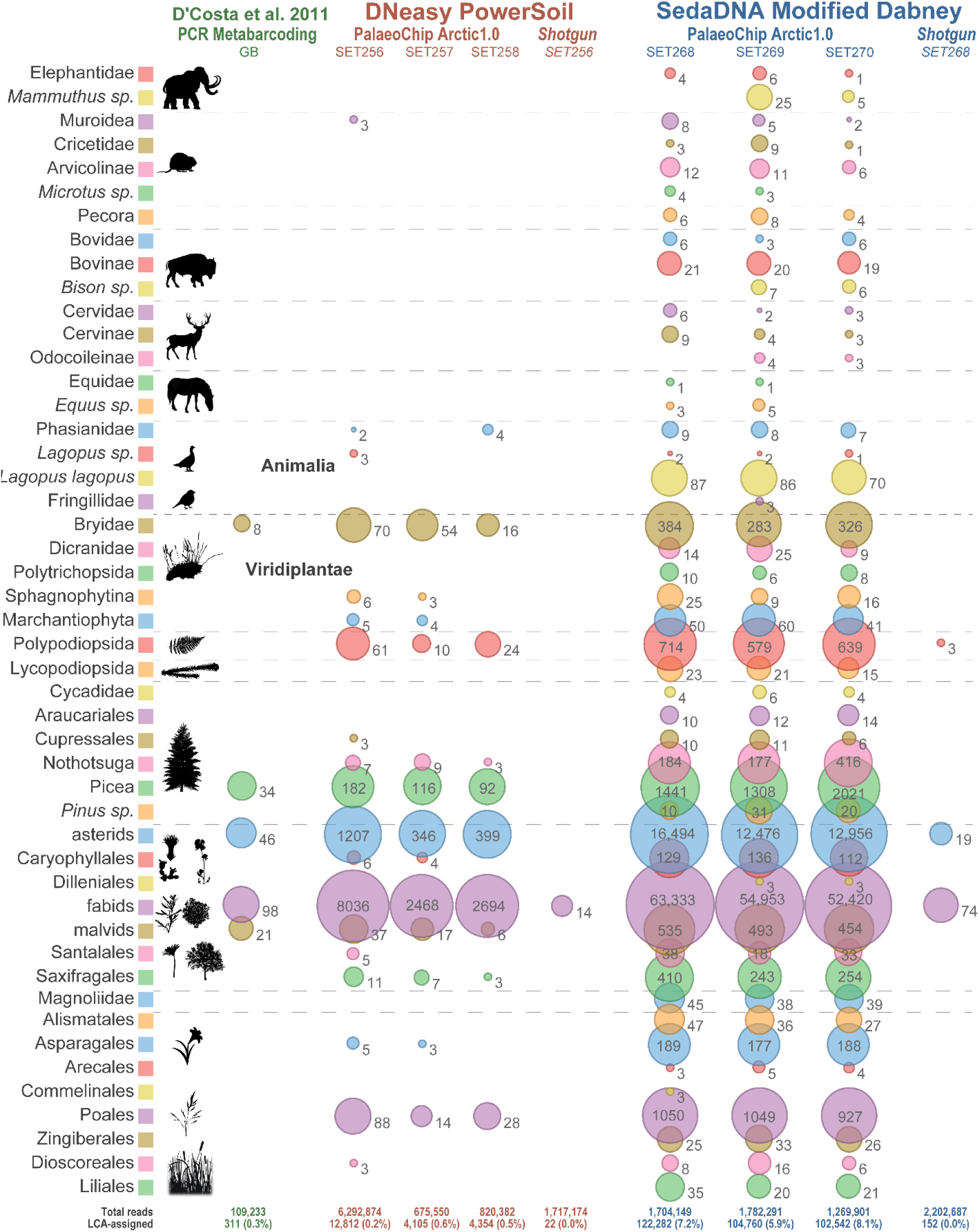
Metagenomic comparison of Upper Goldbottom permafrost core MM12-118b, reads mapped to baits, absolute counts (non-normalized). Core slice dated to 9,685 cal-BP (Mahony, 2015; Sadoway, 2014). Values indicate total reads assigned to that taxon node for Animalia, and a clade summation of reads for Viridiplantae. See Table 1 for read summaries.

**Figure 5.**
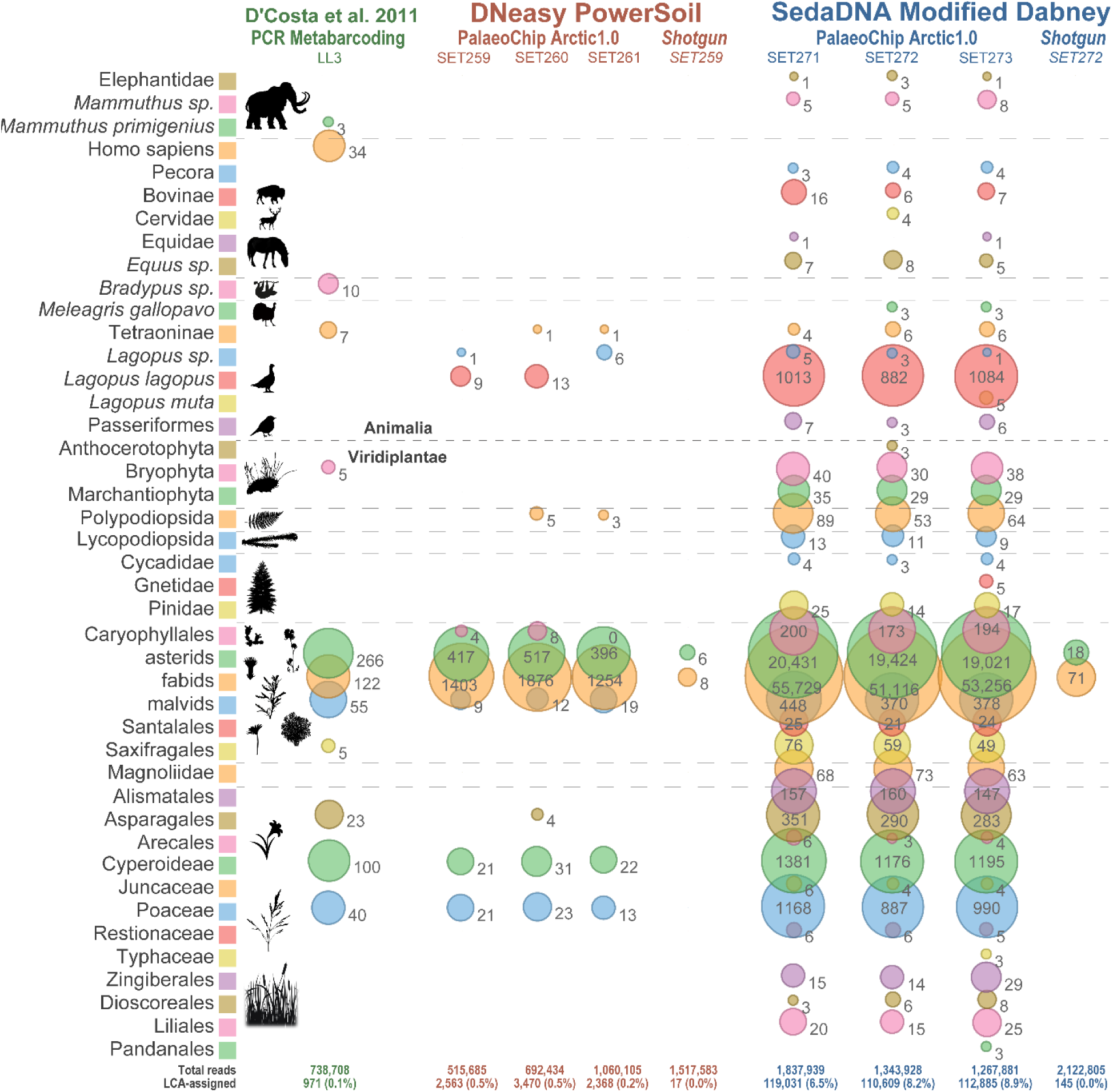
Metagenomic comparison of Lucky Lady II permafrost core LLII-12-84-3, reads mapped to baits, absolute counts (non-normalized). Core slice dated to 13,205 cal-BP (Sadoway, 2014). Values indicate total reads assigned to that taxon node for Animalia, and a clade summation of reads for Viridiplantae. See Table 1 for read summaries.

**Figure 6.**
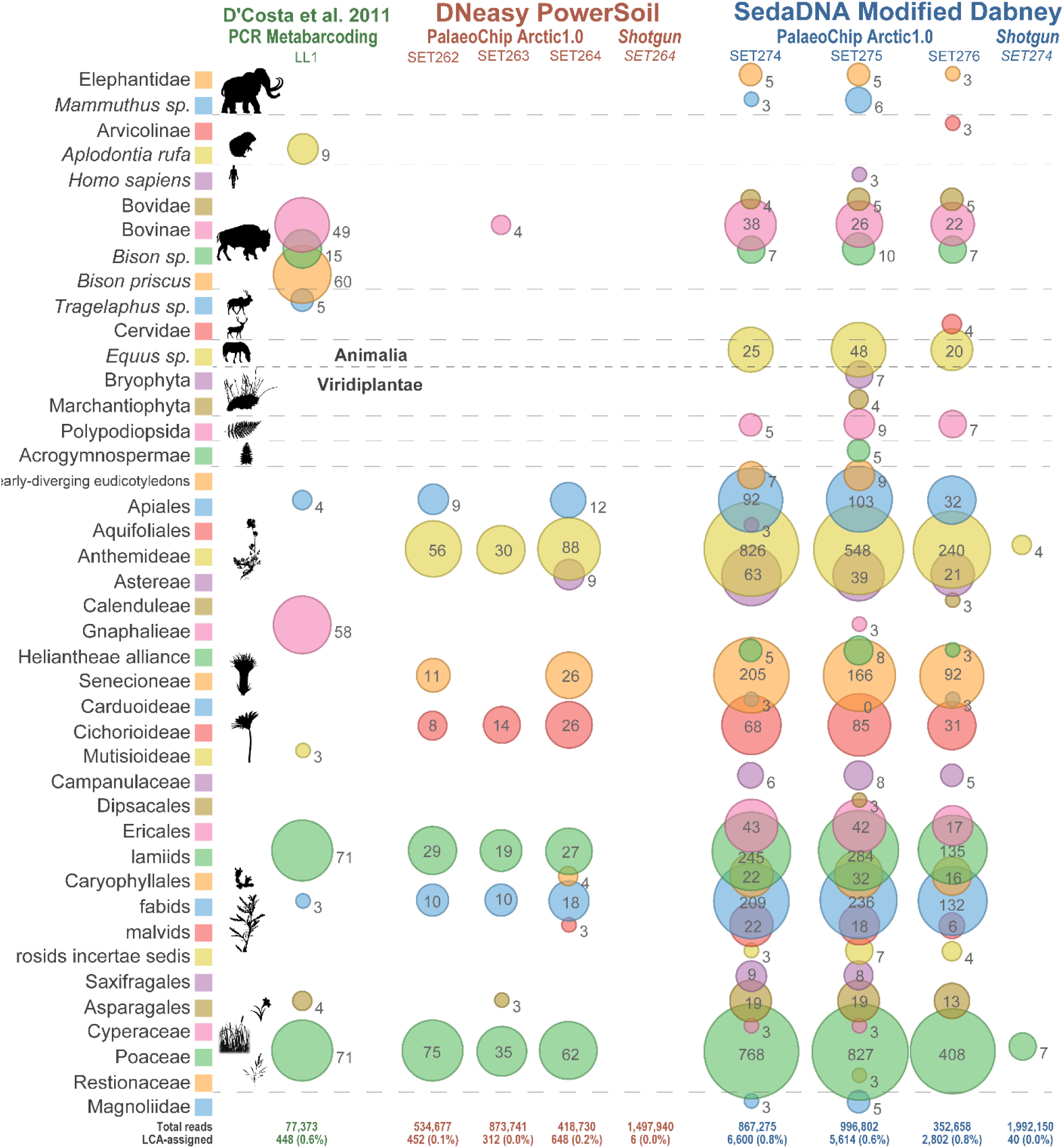
Metagenomic comparison of Lucky Lady II permafrost core LLII-12-217-8, reads mapped to baits, absolute counts (non-normalized). Core slice dated to 15,865 cal-BP (Sadoway, 2014). Values indicate total reads assigned to that taxon node for Animalia, and a clade summation of reads for Viridiplantae. See Table 1 for read summaries.

## Discussion

Our novel 4°C centrifuge inhibitor removal procedure paired with Dabney et al. (2013) aDNA purifications and targeted enrichment consistently outperformed a sedaDNA commercial extraction kit across all extraction replicates, as well as outperforming a PCR metabarcoding approach. These results demonstrate the viability of targeted enrichment for taxonomically diverse environmental samples from open-air sites without the necessity of PCR metabarcoding and the associated compounding biases therein. These data also demonstrate the significantly improved efficiency of ecologically informative sequencing with RNA capture enrichment compared with a shotgun approach. Deep shotgun sequencing to library exhaustion would be ideal as it is the least taxonomically biasing approach. However, until data storage, computational power, database completeness, and sequencing costs are improved, deep sequencing strategies are often unachievable for most users except for those with immense computational and sequencing resources.

### Overcoming enzymatic inhibitors

Of interest for further research is the interaction between SDS and the 4°C spin for inhibitor precipitation (see section “SET-D. SDS and sarkosyl” in Appendix A). We suspect that the efficiency of inhibitor precipitation with this method could be further optimized as experiments suggest that the presence of SDS in the lysis buffer, which we hypothesize leads to the formation of micelles through constant agitation during the spin, significantly contributes to the precipitation of humics and other inhibitors at low temperatures. This technique is unlikely to be optimal for all forms of sedaDNA inhibition however, as it has been observed that identifying the specific inhibitory substances involved is critical to mitigating the compound specific mechanisms that affect enzymatic reactions (Opel et al., 2010). Further research, potentially with mass spectrometry, is needed to identify the inhibitor constituents of sedaDNA target samples in order to improve the inhibitor precipitation we observed while maximizing sedaDNA retention.

### SedaDNA authenticity

Damage profiles for taxa with sufficiently high read counts (≳200 reads at minimum map quality 30) consistently show characteristic aDNA deamination patterns and short fragment length distributions. We observed when mapping to the mitogenome that taxa with ≲200 reads typically have insufficient mapping coverage to confidently identify damage patterns, making it difficult to authenticate rare taxa with low read counts in this dataset. The inflated number of reads mapping to specific taxa compared with the read counts that were *bwa* mapped to our curated baits, *BLASTn* aligned, and *MEGAN* LCA-assigned are suggestive that our quality filtering steps are sufficiently conservative to dramatically reduce the noise characteristic of metagenomic datasets (Eisenhofer et al., 2019; Lu and Salzberg, 2018), but may also strip out some potentially informative (but less confidently assigned) reads. Our pre-*BLASTn* map-filtering approach allows for a much more streamlined analysis with confidently LCA-assigned taxa in the mapped dataset and less confident LCA-assigned taxa in the unmapped reads.

Extraction and library preparation blanks do not contain map-filtered reads (Table 1). The unmapped LCA-assigned reads for these blanks are predominantly adapter contaminated sequences (Figure E13). None of the ecologically informative taxa identified in the metagenomic comparisons appear in the blanks, suggesting patterns observed in our sediment samples are authentic and not the result of contamination.

### Palaeoecology

This study is intended as a proof of concept to demonstrate the viability of targeted enrichment for complex environmental datasets. Additional ongoing research is intended to utilize these methods and complementary palaeoecological techniques on Yukon lake sediment and permafrost cores from the Klondike Plateau (in addition to the metagenomic data from this paper) to temporally track ecological change during the Pleistocene/Holocene transition in Eastern Beringia. However, it is worth briefly contextualizing these broad taxonomic trends here for authenticity purposes. The Bear Creek (BC 4-2B, 30,000 cal-BP [D’Costa et al., 2011]) and older Lucky Lady II core sections (LLII 12-217-8, 15,865 cal-BP [Mahony, 2015]) both date to a period in which Eastern Beringia is thought to have been largely a herb tundra biome, dominated by exposed mineral surfaces, prostrate willows, grasses, forbs (nongraminoid herbs), and occupied by diverse and abundant megafauna (Dyke, 2005; Mahony, 2015). Our data reflects this environmental setting, particularly in the case of the Bear Creek core which has been demonstrated previously to exhibit remarkable preservation (D’Costa et al., 2011). We identified a similar series of mammal species compared to previous work on the same core by D’Costa et al. (2011), but with additional taxa (e.g. caribou, *Rangifer tarandus*) and more specific taxonomic assignments (e.g. potentially yak, *Bos mutus*/*grunniens*). Results from the younger Lucky Lady II core section (LLII 12-84-3, 13,205 cal-BP [Sadoway, 2014]) indicate an expansion of Western Beringian birch shrub tundra (Dyke, 2005), reflected by a decrease in grasses and a proportional increase in *Betula* and *Salix*. Analysis of the most recent core sample (MM12-118b, 9,685 cal-BP [Mahony, 2015]), suggests that there had been a shift in the forest/tundra ecotone in the Yukon before the development of the boreal (taiga) forest, which first established in southern Yukon by ∼9,000 cal-BP (Dyke, 2005). Our data shows a proportional increase in conifers, particularly spruce (*Picea*).

The mammalian constituents also display a marked change, dwindling moving forward in time into the Holocene (Figures 4–6), but perhaps less sharply than commonly thought. For example, we recovered genetic evidence of both woolly mammoth (*Mammuthus primigenius*) and horse (*Equus sp*.) in the Upper Goldbottom core dated to ∼9,685 cal-BP (Mahony, 2015). Previous radiocarbon analyses appeared to indicate that horses disappeared from high-latitude northwestern North America relatively early, ca. 12,500 ^14^C BP (“last appearance date” 13,125 cal-BP, based on AMNH 134BX36 from Upper Cleary Creek [Guthrie, 2003]). This ∼3,500 year difference implies the existence of a substantial *ghost range* (Haile et al., 2009) (i.e., spatiotemporal range extending beyond its last appearance age as indicated by directly dated materials) that cannot (yet) be corroborated by the macrofossil record for *Equus*, but consistent with previous ancient eDNA results from central Alaska (Haile et al., 2009). However, in the absence of additional information, it is difficult to assess whether this signal may be considered chronostratigraphically reliable, or may have been affected by factors such as leaching, cryo- or bioturbation, or reworking (redeposition) (Arnold et al., 2011), altering the relative positions of sequenced sedaDNA organomineral complexes. In the case of the mammoth reads, after merging the sequenced data from the three Upper Goldbottom core (MM12-118b) extraction replicates, coverage was insufficient (low read count) to reliably assess characteristic aDNA damage patterns (Figure E12), although there is arguably some indication of damage with the merged *mapDamage* and FLD profile. We hope that with subsequent sedaDNA analyses from this region (with geographic targets selected to minimize the probability of containing stratigraphically allochthonous sedaDNA) we will be more able to closely evaluate the temporal authenticity of the megafaunal *ghost ranges* hinted at here with this dataset.

### Limitations of comparison

There are several caveats to keep in mind when assessing our comparison of protocols and the application PalaeoChip Arctic1.0. First, the lysing stage of our PowerSoil and Modified Dabney protocols were not equivalent in duration or reagents. We followed manufacturer specifications for PowerSoil, but the lysis stage of extraction with equivalent kits can be increased in duration and augmented with additional reagents to theoretically increase DNA yield (Niemeyer et al., 2017). Further, a recently released update to the PowerSoil kit, the DNeasy PowerSoil Pro, claims to have an 8-fold increase in DNA yield compared with comparative commercial kits (it is unclear what the fold increase over standard PowerSoil is with this updated kit). Our experiments with the PowerSoil inhibitor removal solution C3 found consistently low DNA retention compared with our longer duration 4°C spin as an inhibitor removal technique (see section SET-B in Appendix A). The PowerSoil inhibitor removal solution is effective at rapidly precipitating enzymatic inhibitors, but this study suggests that it is overly aggressive and consistently precipitated viable sedaDNA in the process (see Figure S7 in Appendix A). We suspect that a longer lysis stage with PowerSoil would increase overall yields, but would not mitigate the substantial losses associated with overly aggressive humic precipitation when utilizing solution C3. We found that the 4°C spin is sufficiently effective at removing enzymatic inhibition to allow for successful adapter ligation, even if the extracts were not as inhibitor free as PowerSoil (Figure 3).

Second, metabarcoding is not directly equivalent to enrichment when comparing taxonomic coverage and LCA-assigned read counts. Metabarcoding is not limited by pre-synthesized baits, but rather by primer design (binding site conservation, database completeness, etc.). Mapping our data back to the baits does strip out potentially taxonomically informative hits *a priori*. To mitigate this, we have included a non-mapped comparative variant in Appendix A (see section SET-E). We observe that mapping to the curated baits (which have low complexity and non-diagnostic regions masked or removed) substantially reduces the number of low confidence (potential false-positive) spurious hits. Although, it is still worth looking through non-mapped data for positive, bait off-target assignments. This component highlights a limitation of enrichment overall, that one’s taxonomic recovery is limited by the quality of the relevant bait-set, whereas PCR metabarcoded data is limited by biased PCR amplifications (low abundance taxa being swamped out by the over-amplification of high abundance or comparatively undamaged molecules) as well as primer sequence conservation (or lack thereof), and database completeness for design and alignment.

However, taxonomic alignments using amplicon sequence data may improve in the future as reference databases are updated. For example, *Sus scrofa* is not included in our targeted baits as members of this family (which includes peccaries or New World pigs) are not considered to have been present in the subarctic during the Pleistocene/Holocene transition (see Appendix B, note that flat-headed and collared peccary [*Pecari tajacu* and *Platygonus peccary* respectively] are included in the bait-set). However, Sadoway’s (2014) metabarcoding approach was able to amplify suid reads in core BC 4-2B (Figure 7), which pass mapped-LCA filtering. Although some species of peccaries were able to live close to the ice margin in mid-continental North America during the late Pleistocene (Kurten and Anderson, 1980), there is no palaeontological evidence for their presence at that time in the Yukon (Stuart, 2015). In this case, one suspects that those metabarcoding reads are amplified contaminants. Regardless of specific instances like this, it is reasonable to employ *a priori* limitations on a capture-based approach for filtering out taxa (either intentionally or unintentionally) prior to sequencing. The false-negative error potential of enrichment with sedaDNA can be mitigated with improved iterations of the bait-set (e.g. PalaeoChip Arctic2.0, Mediterranean1.5) as genetic reference databases are updated.

**Figure 7.**
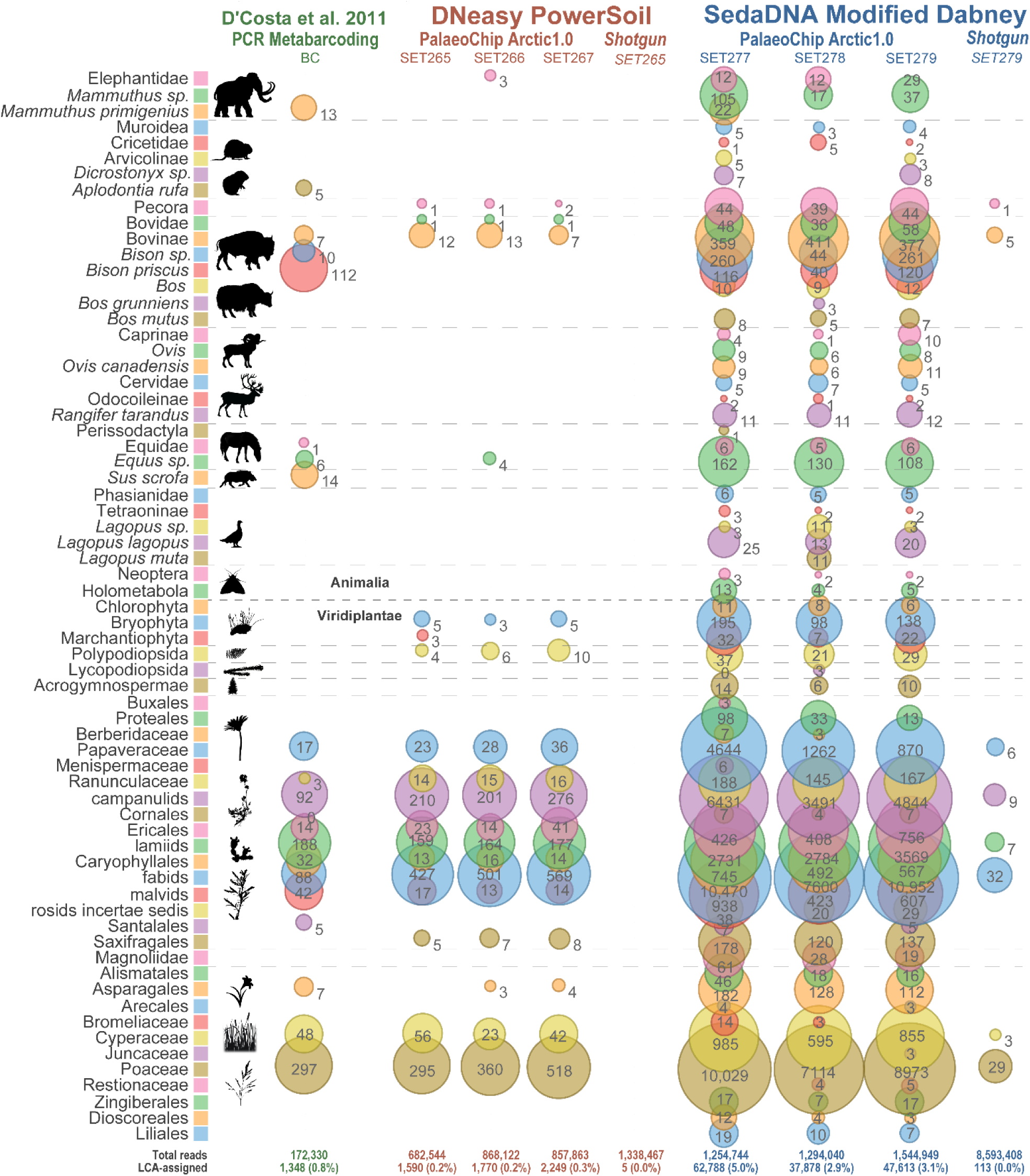
Metagenomic comparison of Bear Creek permafrost core BC 4-2B, reads mapped to baits, absolute counts (non-normalized). Core slice dated to ∼30,000 cal-BP (D’Costa et al., 2011; Mahony, 2015; Sadoway, 2014). Values indicate total reads assigned to that taxon node for Animalia, and a clade summation of reads for Viridiplantae. See Table 1 for read summaries.

It should be noted that the PCR metabarcoding data utilized in this analysis was not extracted with our sedaDNA optimized strategy. It is likely that a metabarcoding approach may also be further improved by utilizing a 4°C inhibitor removal procedure to maximize sedaDNA retention. We acknowledge that this is an unfair limitation of our comparison, as the extraction methodology between our metabarcoding and enriched data are not equivalent. This topic is worthy of further research but lies outside the scope of our analysis here, which is intended to establish the viability of enrichment for sedaDNA contexts, and to report on a new inhibitor removal technique that may be further optimized. We intend to expand on PalaeoChip Arctic1.0 with further target sequences for regionally specific vegetation, mammals, insects, fungi, and microbiota. PalaeoChip is also intended to be optimized for other non-arctic regions.

## Conclusion

We believe the experiments outlined in this report through the development of a Holarctic PalaeoChip, in this case Arctic1.0, clearly demonstrate the utility of our novel inhibitor removal technique paired with high volume Dabney et al. (2013) purifications for overcoming enzymatic inhibitors, which likewise enables the viability of targeted enrichment for sedaDNA analyses. This approach can simultaneously capture complex environmental DNA without excessive PCR amplifications (and the associated costs of using high concentrations of polymerase to overcome co-eluted enzymatic inhibitors). By increasingly iterating on the taxonomic breadth of our complex environmental baits, and by further optimizing enrichment and sedaDNA extraction conditions, this technique can continue to improve the sequenced fraction of on-target molecules without deep shotgun sequencing, or potentially biased PCR amplifications. We believe our extraction method and enrichment strategy are of relevance not only for ancient DNA research, but also potentially for modern environmental DNA monitoring applications where the DNA copy number is high and targeted capture is of even more relevance for improving the analytic and cost efficiency of high-throughput metagenomic sequencing.

## Methods

All supplementary procedures, tables, and figures reported here with an ‘S’ prefix (e.g. Table S1, Figure S1) are included in the supplementary online materials in Appendix A.

### Lab setting

Laboratory work was conducted in clean rooms at the McMaster Ancient DNA Centre, which are subdivided into dedicated facilities for sample preparation, stock solution setup, and DNA extraction through library preparation. Post-indexing and enrichment clean rooms are in a physically isolated facility, while high-copy PCR workspaces are in separate building with a one-way workflow progressing from low-copy to high-copy facilities. Each dedicated workspace is physically separated with air pressure gradients between rooms to reduce exogenous airborne contamination. Prior to all phases of laboratory work, dead air hoods and workspaces were cleaned using a 6% solution of sodium hypochlorite (commercial bleach) followed by a wash with Nanopure purified water (Barnstead) and 30 minutes of UV irradiation at >100 mJ/cm^2^.

#### 1. Subsampling

Metal sampling tools were cleaned with commercial bleach, rinsed with Nanopure water immediately thereafter (to reduce rusting), and heated overnight in an oven at ∼130°C. Once the tools had cooled, work surfaces were cleaned with bleach and Nanopure water and covered with sterile lab-grade tin foil. Sediment cores previously split into disks (D’Costa et al., 2011, p. SI. 4–5; Sadoway, 2014, chap. 1) and stored at −20°C had the upper ∼1 mm of external sediment chiselled off to create a fresh sampling area free of exogenous contaminants for a hollow cylindrical drill bit. The drill bit (diameter 0.5 cm) was immersed in liquid nitrogen prior to sampling and a drill press was used to repeatedly subsample the disk sections(D’Costa et al., 2011, fig. S3). Sediment was pushed out of the drill bit using a sterile nail and the bottom 1–2 mm of sediment from the bit was removed before dislodging the remaining sample. This exterior core portion was carefully removed as it has a higher chance of containing sedaDNA from other stratigraphic contexts due to coring and core splitting. A bulk set of subsampled sediment from the same core disk was homogenized by stirring in a 50 mL falcon tube and stored at −20°C for subsequent extractions. This process was repeated individually for each core.

#### 2. Physical disruption, chemical lysis, and extraction

DNeasy PowerSoil DNA Extraction kit samples were extracted following manufacturer specifications; purified DNA was eluted twice with 25 µL EBT buffer (10 mM Tris-Cl, 0.05% Tween-20). Samples extracted with the sedaDNA modified Dabney et al. (2013) procedure were processed as follows (see Figure 2):

##### Lysis

1. 500 μL of a digestion solution (see Table S1) initially without proteinase K was added to PowerBead tubes (already containing garnet beads and 750 µl 181 mM NaPO4 and 121 mM guanidinium isothiocyanate).
2. 250 mg of homogenized sediment was added to each PowerBead tube.
3. PowerBead tubes were vortexed at high speed for 15 minutes, then centrifuged briefly to remove liquid from the lids.
4. 15.63 µL of proteinase K (stock 20 mg/mL) was added to each tube to reach a proteinase K concentration of 0.25 mg/mL in the digestion and PowerBead solution (1.25 mL).
5. Tubes were finger vortexed to disrupt sediment and beads that had pelleted in step 3.
6. PowerBead tubes were securely fixed in a hybridization oven set to 35°C and rotated overnight for ∼19 hours, ensuring that the digestion solution, sediment, and PowerBeads were moving with each oscillation.
7. PowerBead tubes were removed from the oven and centrifuged at 10,000 x g for 5 minutes (the maximum speed recommended for PowerBead tubes). Supernatant was pipetted into a MAXYMum Recovery 2 mL tube and stored at −20°C.

##### Purification

8. Digestion supernatant was thawed, briefly centrifuged, pipetted into 16.25 mL (13 volumes) of high-volume Dabney binding buffer (see Table S2) in a 50 mL falcon tube and mixed.
9. Falcon tubes were spun at 4500 rpm in a refrigerated centrifuge set to 4°C for 20 hours overnight.
10. After centrifugation, falcon tubes were carefully removed and the supernatant was decanted, taking care to not disturb the darkly coloured pellet that had formed during the cold spin at the base of the tube.
11. The binding buffer was passed through a high-volume silica-column (High Pure Extender Assembly, Roche Diagnostics) and extraction proceeded as per Dabney et al. (2013).
12. Purified DNA was eluted off the silica columns twice with 25 µL EBT.

Prior to all subsequent experiments, both the modified Dabney and PowerSoil extracts were centrifuged at 16,000 x g for 5 minutes to pellet remaining co-eluted inhibitors.

#### 3. Library preparation

Doubled stranded libraries were prepared as described in Meyer and Kircher (2010) with modifications from Kircher et al. (2012) and a modified end-repair reaction to account for the lack of uracil excision (Table S3). Samples were purified after blunt-end repair with a QIAquick PCR Purification Kit (QIAGEN) (to maximally retain small fragments) and after adapter ligation (Table S4) with a MinElute PCR Purification Kit (QIAGEN), both using manufacturers protocols. Library preparation master mix concentrations can be found in Tables S3-S6.

#### 4. QPCR: Inhibition spike tests, total quantification, and indexing

A positive control spike qPCR assay (Enk et al., 2016; King et al., 2009) was used to assess the relative impact of DNA independent inhibitors (co-eluted substances such as humics that inhibit enzyme function) on the enzymatic amplification efficiency of a spiked amplicon in the presence of template sedaDNA derived from variable lysing and extraction methods (Table S7). We suspected that enzymes in library preparation would be inhibited similarly to AmpliTaq Gold polymerase in qPCR. Shifts in the qPCR amplification slope of our spiked oligo with AmpliTaq Gold (due to co-eluted inhibitors in sedaDNA extracts) could then be quantified and used to infer the likelihood of failed adapter ligation due to enzymatic inhibitors (rather than a lack of sedaDNA). Admittedly, AmpliTaq Gold is not a 1:1 stand-in for inhibition sensitivity during blunt-end repair and adapter ligation, as AmpliTaq is among the most sensitive polymerases to inhibition induced reductions in amplification efficiency (Al-Soud and Radstrom, 1998), and due to qPCR specific inhibition such as the reduction in florescence despite successful amplification (Sidstedt et al., 2015). Our experiments do suggest that these enzymes have a very roughly commensurate inhibition sensitivity, insofar as eluates completely inhibited during this spike test are unlikely to successfully undergo library adapter ligation.

To quantify the co-eluted inhibition affecting each spiked amplification, we compared the qPCR slope of an oligo-spiked sedaDNA extract (1 µL of sample eluate spiked with 1 µL of a 49-bp oligo [1000 copies {E^3^}], see Table S7) with the qPCR slope of 1 µL E^3^ oligo standard in 1 µL of EBT. Average Cq and max relative fluorescence units (RFU) for each PCR replicate were calculated, as was the hill slope of the amplification curve by fitting a variable-slope sigmoidal dose-response curve to the raw fluorescence data using GraphPad Prism v. 7.04 (based on King et al. [2009]). The E^3^ oligo-spiked averages (Cq, RFU, and sigmoidal hillslope) were divided by the corresponding E^3^ oligo standard amplification value, then averaged together to generate an ‘inhibition index’ per PCR replicate, which were averaged again across PCR replicates to determine an extract’s inhibition index, ranging from 0–1. In this case, 0 indicates a completely inhibited reaction (no measurable increase in RFU), and 1 indicates a completely uninhibited reaction relative to the spiked E^3^ oligo-standard (see Figure E14). Anything above 0.9 (the bottom range for blanks and standards of differing starting quantities) is considered essentially uninhibited insofar as *Taq* polymerase inhibition is concerned.

Most total DNA quantifications in SET-A to SET-D2 (detailed in the supplementary materials) used the short amplification primer sites on the library adapters and were compared against the same library prepared 49-bp oligo standard used in the spike tests (see Table S8). This assay was also modified in some instances to quantify the ‘endogenous’ chloroplast constituent of adapted molecules by pairing the *trn*L P6-loop forward primer-g (Taberlet et al., 2007) with the reverse P7R library adapter primer (IS8, see Table S9). Enk et al. (2013) demonstrated that a single-locus qPCR assay can be used to predict on-target ancient DNA high-throughput sequencing read counts. Previous analyses (D’Costa et al., 2011; Sadoway, 2014) indicated that ancient vegetation was the most consistently abundant fraction of the biomolecules in these cores, and as such could serve as a rough proxy for assessing aDNA retention for successfully library adapted molecules between various inhibitor removal strategies. For all qPCR results reported here, standard curve metrics are included in the associated captions. Ideal standard curve values are: R^2^ = 1, slope = −3.3 (or between −3.1 and −3.5), efficiency = 90–105%.

The extraction triplicate with the highest DNA concentrations for each of the four cores and two extraction methods (as based on the short amplification qPCR) was indexed for shotgun sequencing (8 samples + 3 blanks). All extraction triplicates (24 samples + 3 blanks) were indexed separately thereafter for targeted enrichment.

#### 5. Enrichment: PalaeoChip, Arctic1.***0***

The PalaeoChip, Arctic1.0 RNA hybridization bait-sets were designed to target whole mtDNA of extinct and extant Quaternary animals (focused primarily on megafauna; number of taxa ≈ 180), and high latitude plant cpDNA based on curated reference databases developed by Sønstebø et al. (2010), Soininen et al. (2015), and Willerslev et al. (2014), initially targeting *trn*L (n ≈ 1650 taxa) (see Appendix B for taxonomic list). This list was queried with the *NCBI Mass Sequence Downloader* software (Pina-Martins and Paulo, 2015) to recover additional nucleotide data from GenBank (Benson et al., 2018) for *trn*L, as well as adding targets for *mat*K and *rbc*L. These three regions were selected as they are among the most sequenced and taxonomically informative portions of the chloroplast genome (Hollingsworth et al., 2011). Baits were designed in collaboration with Arbor Biosciences to 80 bp with ∼3x flexible tiling density, clustered with >96% identity and >83% overlap, and baits were removed with >25% soft-masking (to reduce low complexity baits with a high chance of being off-target in complex environmental samples). Bait sequences were queried with *BLASTn* against the NCBI database on a local computer cluster using a July 2018 database, then inspected in *MEGAN* (Huson et al., 2016, 2007). Baits with a mismatched taxonomic target and *BLASTn* alignment were queried again using a web-blast script (Camacho et al., 2009; NCBI Resource Coordinators, 2018) to determine if these mismatches were due to local database incongruities. Mismatches were again extracted with *MEGAN*, individually inspected, then removed from the bait-set if determined to be insufficiently specific.

Hybridization and bait mixes were prepared to the concentrations in Table S11. For each library, 7 µL of template was combined with 2.95µL of Bloligos (blocking oligios which prevent the hybridization between library adapter sequences). The hybridization and bait mixes were combined and pre-warmed to 60°C, before being combined with the library-Bloligo mixture. The final reaction was incubated for 24 hours at 55°C for bait-library hybridization.

The next day, beads were dispensed (540 µL total between two tubes), washed three times with 200 µL of binding buffer for each tube, then suspension in 270 µL of binding buffer per tube and aliquoted into PCR strips. Baits were captured using 20 µL of the bead suspension per library, incubated at 55°C for 2.5 minutes, finger vortexed and spun down, and incubated for another 2.5 minutes. Beads were pelleted and the supernatant (the non-captured library fraction) was removed and stored at −20°C. The beads were resuspended in 180 µL of 55°C Wash Buffer X and washed four times following the MYbaits V4 protocol. Beads were eluted in 15 µL EBT, PCR reamplified (Table S6), then purified over MinElute following manufacturer’s protocols in 15 µL EBT.

#### 6. Post-indexing total quantification, pooling, size-selection, and sequencing

Enriched and shotgun samples were quantified using the long-amplification total library qPCR assay (Table S10). Enriched and shotgun libraries were equal-molar pooled separately. The two pools were size-selected with gel excision following electrophoresis for molecules ranging from 150 bp to 600 bp. Gel plugs were purified using the QIAquick Gel Extraction Kit (QIAGEN), according to manufacturer’s protocol, then sequenced on an Illumina HiSeq 1500 with a 2 × 90 bp paired-end protocol at the Farncombe Metagenomics Facility (McMaster University, ON).

#### 7. PCR Metabarcoding

Sadoway (2014) previously worked with these and many other open-air Yukon permafrost core samples using a metabarcoding approach. These libraries had been extracted in duplicate with guanidinium protocols (Boom et al., 1990; D’Costa et al., 2011) from 250 mg of the same core sections, purified with silica (Höss and Pääbo, 1993), and eluted twice (Handt et al., 1996). Extensive inhibition at the time, as detected using similar qPCR spike tests developed by King et al. (2009), necessitated a tenfold extract dilution, which were then amplified in duplicate for each primer set, targeting: *rbc*L (CBOL Plant Working Group, 2009; Hollingsworth, 2011; Willerslev et al., 2003), *trnL* (Taberlet et al., 2007), *16S rRNA* (Höss et al., 1996), *and 12S rRNA* (Kuch et al., 2002), each following cited PCR conditions. The locus *cytochrome b* (*cyt-b*) was also targeted using a set of degenerate primers designed with FastPCR (Kalendar et al., 2011; Sadoway, 2014). *Cyt-b* amplifications were found to be most efficient in 20 µL reactions using AmpliTaq Gold (0.05U/µL), 1X PCR Buffer II, 2.5 mM MgCl2, 0.25 mM dNTPs, 0.5X Evagreen, 250 nM (forward/reverse primers) when cycled with a 3 minute denaturation at 95°C, 45 cycles of 95°C for 30 seconds, and 60°C for 30 seconds (Sadoway, 2014). QPCR products were purified with 10K AcroPrep Pall plates (Pall Canada Direct Ltd., Mississauga, ON, Canada) using a vacuum manifold. QPCR assays were used to pool each amplicon set in equimolar concentrations, which were library prepared and dual-index following the same Illumina protocols as described above (Kircher et al., 2012; Meyer and Kircher, 2010). Each sample extract had its own unique combination of forward and reverse indices, which were sequenced on a HiSeq 1500 Rapid Run (2 × 100bp, Illumina Cambridge Ltd, Essex, UK) at the Farncombe Metagenomics Facility (McMaster University, ON) to approximately 100,000 reads each. These PCR metabarcoded libraries were processed for this paper with an identical bioinformatic pipeline as described below.

#### 8. Bioinformatics

Reads from all library sets (enriched, shotgunned, and PCR metabarcoded) were demultiplexed with *bcl2fastq* (v 1.8.4), converted to bam files with *fastq2bam* (https://github.com/grenaud/BCL2BAM2FASTQ), then trimmed and merged with using *leeHom* (Renaud et al., 2014) using ancient DNA specific parameters (--ancientdna). Reads were then aligned to a concatenated reference of the animal and plant bait-set with *network-aware-BWA* (Li and Durbin, 2009) (https://github.com/mpieva/network-aware-bwa) with a maximum edit distance of 0.01 (-n 0.01), allowing for a maximum two gap openings (-o 2), and with seeding effectively disabled (-l 16500). Mapped reads that were either merged or unmerged but properly paired were extracted with *libbam* (https://github.com/grenaud/libbam), collapsed based on unique 5’ and 3’ positions with *biohazard* (https://bitbucket.org/ustenzel/biohazard), and restricted to a minimum length of 24bp. Both mapped and non-map filtered reads were string deduplicated using the *NGSXRemoveDuplicates* module of *NGSeXplore* (https://github.com/ktmeaton/NGSeXplore), then queried with *BLASTn* to return the top 100 alignments (-num_alignments 100 -max_hsps 1) against a July 2018 version of the NCBI Nucleotide database on a local computer cluster. Non-map filtered libraries were treated identically, although only returned the top 10 alignments to mitigate unwieldy (>10 gb) file sizes. Sequencing summary counts are in Table 1.

Blast and fasta files for each sample (unmapped and mapped variants) were passed to *MEGAN* (Huson et al., 2016, 2007) using the following LCA parameters: min-score = 50 (default), max expected (e-value) = 1.0E-5, minimum percent identity = 95% (allows 1 base mismatch at 24 bp, 2 at 50 bp, and 3 at 60 bp to account for cytosine deamination and other aDNA characteristic damage or sequencing errors), top percent consideration of hits based on bit-score = 15% (allows for slightly more conservative taxonomic assignments than the 10% default based on trial and error), minimum read support = 3 (number of unique reads aligning to an NCBI sequence for that taxon to be considered for LCA), minimum complexity = 0.3 (default minimum complexity filter), and utilizing the LCA weighted algorithm at 80% (two rounds of analysis that purportedly increases specificity but doubles run time over the native algorithm). Metagenomic profiles were compared in *MEGAN* using absolute and normalized read counts (normalized in *MEGAN* to the lowest read count, keeping at least 1 read per taxon). Libraries were not subsampled to an equal depth prior to processing; McMurdie and Holmes (2014) have demonstrated that this rarefying approach is the most ineffective means of accounting for unequally sequenced metagenomic data. However, in an attempt to fairly account for biases inherent in comparing capture enriched and amplicon data of unequal sequencing depths, we have included four versions of the bubble chart taxonomic comparisons per core: 1) absolute counts mapped filtered to our baits (in the main text) with total reads and proportional LCA-assigned totals at the bottom of each column, 2) normalized counts mapped to baits, 3) absolute counts for all reads (not map filtered), and 4) normalized counts for all reads (not map filtered). The latter three comparisons are included in Appendix A.

To visualize the taxonomic variability between these replicates, comparative trees in *MEGAN* were uncollapsed to the ‘order’ rank (meaning that all lower taxonomic assignments within that ‘order’ are summed); animalia was then fully uncollapsed (as the read counts were more manageable compared with plant assignments). Viridiplantae clades were collapsed to higher ranks (higher than ‘order’) in some cases for summarized visualizations (otherwise there were too many leaves to display at once in a single figure, even if when only showing summaries by ‘order’). Thereafter, all leaves were selected and visualized with logarithmically scaled bubble charts; additional higher LCA-assigned animalia ranks were also selected where taxonomically informative (for example, reads that could only be conservatively LCA-assigned to Elephantidae or *Mammuthus sp*., but which in this context likely represent hits to *Mammuthus primigenius* [woolly mammoth]). Low abundance (<3 reads), non-informative and non-target clades (e.g. bacteria, fungi, or LCA-assignments to high ranks) were excluded for visualization purposes. Reads within viridiplantae frequently hit low taxonomic ranks (family, genus, and occasionally species) but this resolution is not shown here to facilitate our goal of an inter-method comparison.

Taxa with high blast and LCA-assigned read-counts were also selected to evaluate damage patterns and fragment length distributions (FLD) (Table E2). Enriched libraries were mapped to reference genomes of either the LCA-assigned organism itself (e.g. *Mammuthus primigenius*) or a phylogenetically closely related organism (e.g. *Equus caballus*) if there was no species call or if a reliable reference genome does not yet exist. Mapping followed the aforementioned parameters and software, with an additional map-quality filter to ≥30 with *samtools* (https://github.com/samtools/samtools) and passed to *mapDamage* (Jónsson et al., 2013) (v 2.0.3, https://ginolhac.github.io/mapDamage/). Plant chloroplast DNA references were reduced to the target barcoding loci (*trn*L, *rbc*L, and *mat*K), each separated by 100 Ns. Mitochondrial reference genomes were used for animal taxa of interest.

## Supporting information

Optimization experiments and supplemental materials to manuscript

List of taxa in PalaeoChip Arctic1.0 bait-set

## Acknowledgements

We wish to thank the Arctic Institute of North America, the Canadian Institutes of Health Research, the Garfield Weston Foundation, the Natural Sciences and Engineering Research Council of Canada, McMaster University, Polar Knowledge Canada (POLAR), and the Social Sciences and Humanities Research Council of Canada for each funding various components of this research.

## Main Text Extended Tables

**Table E2.**
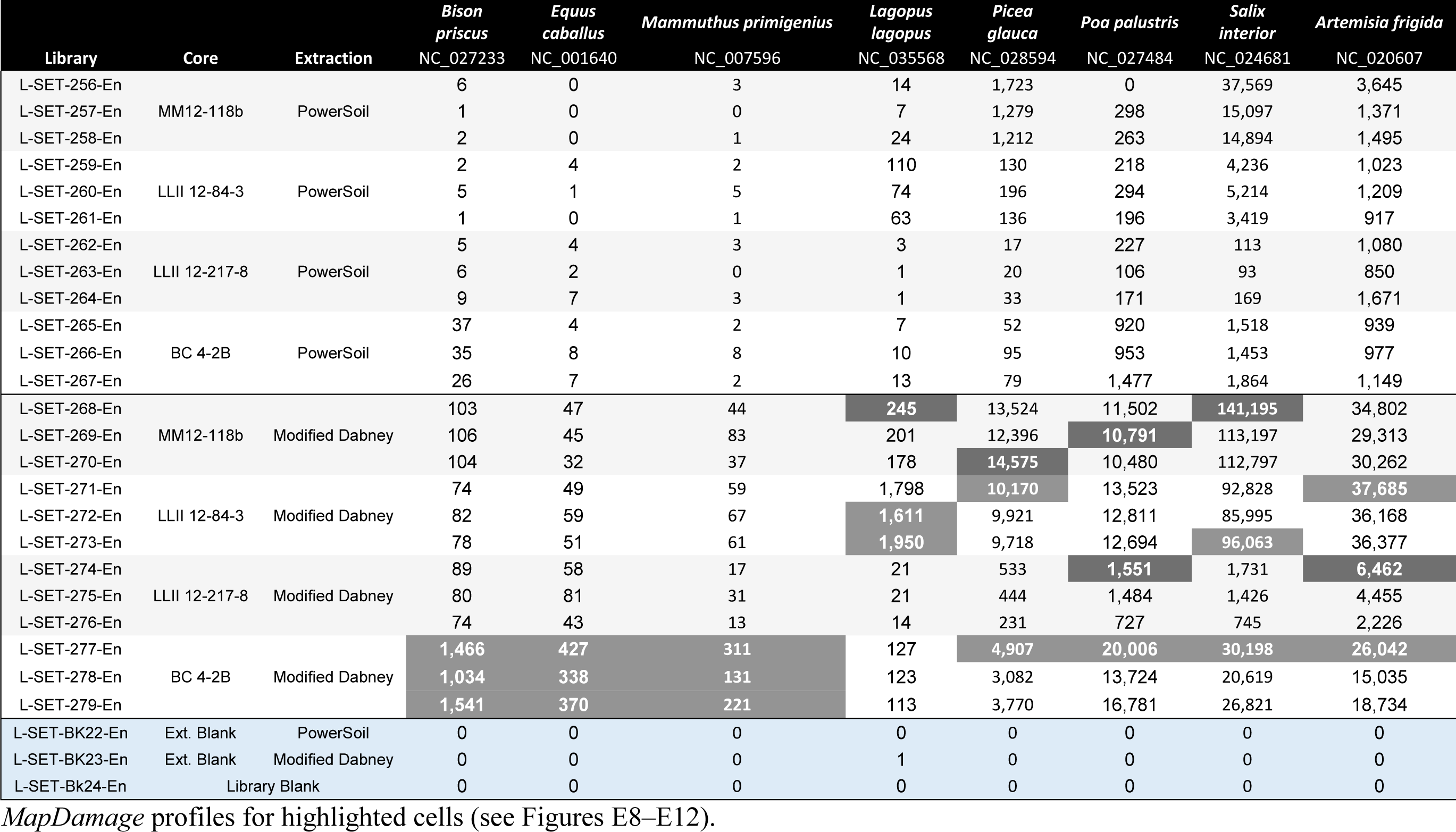
Taxon specific mapping summary at a minimum length of 24 bp and mapping quality of 30.

## Main Text Extended Figures

**Figure E8.**
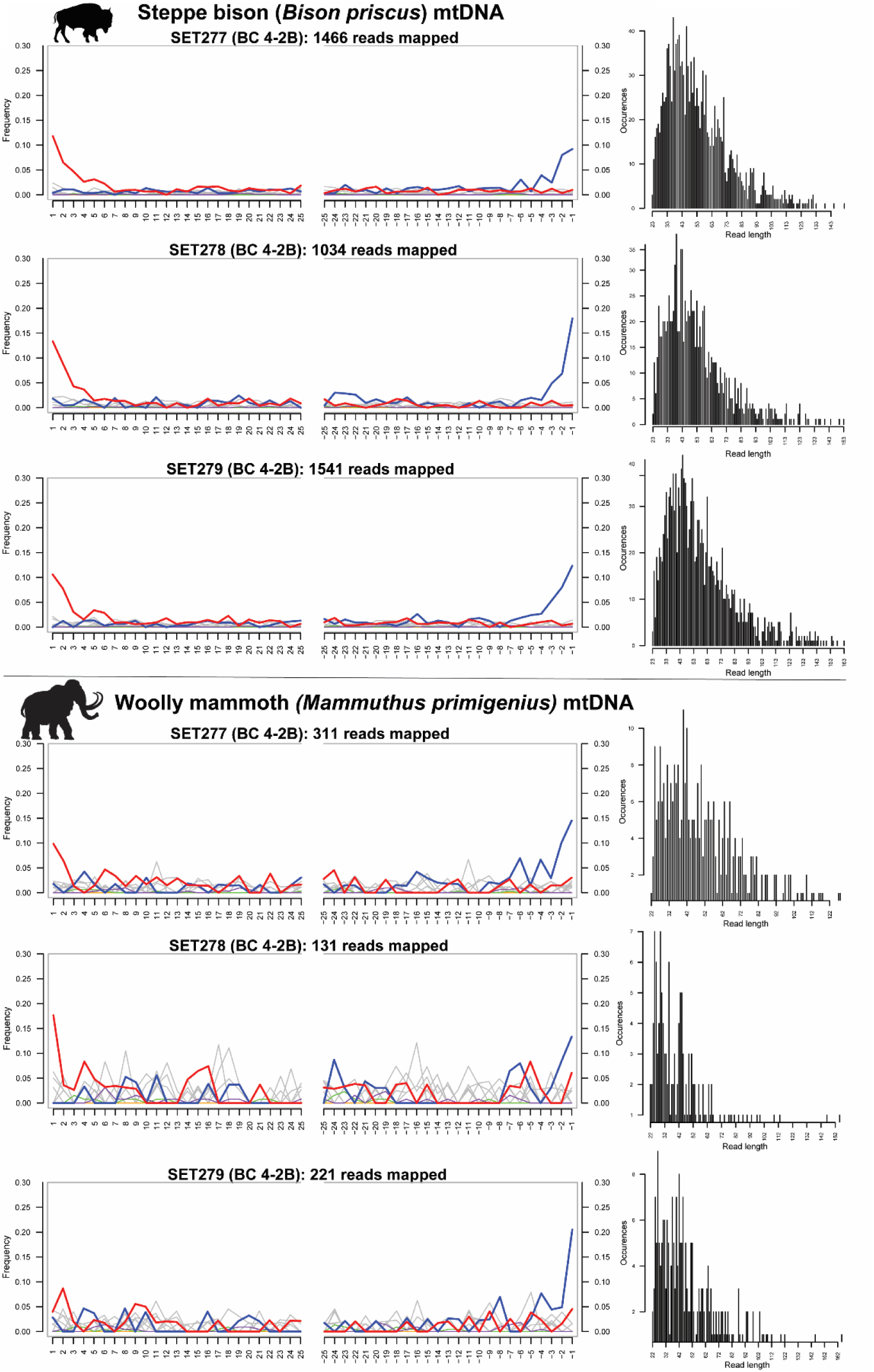
*MapDamage* plots for *Bison priscus* and *Mammuthus primigenius*. Minimum length = 24 bp, minimum mapping quality = 30.

**Figure E9.**
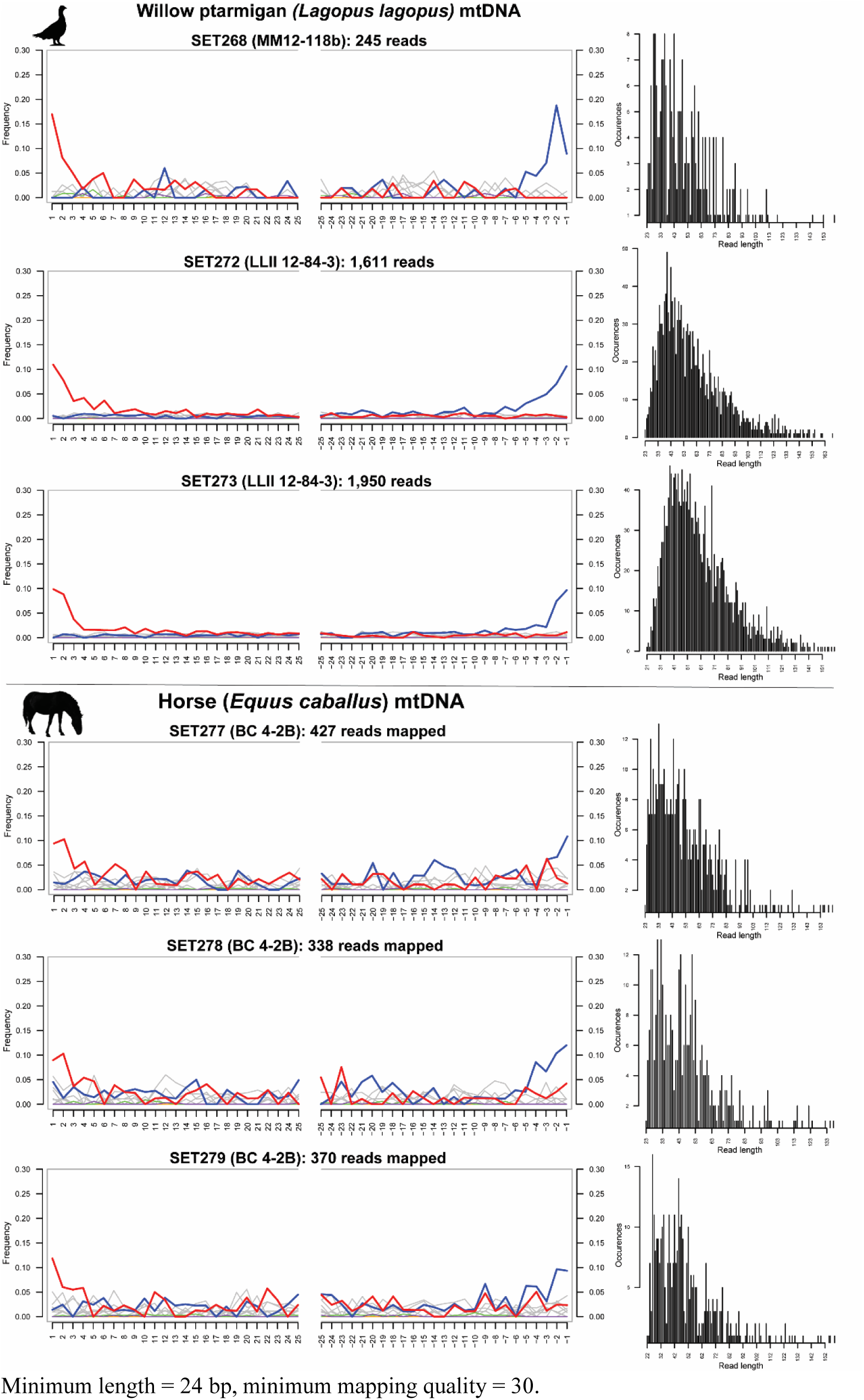
*MapDamage* plots for *Lagopus lagopus* and *Equus caballus*. Minimum length = 24 bp, minimum mapping quality = 30.

**Figure E10.**
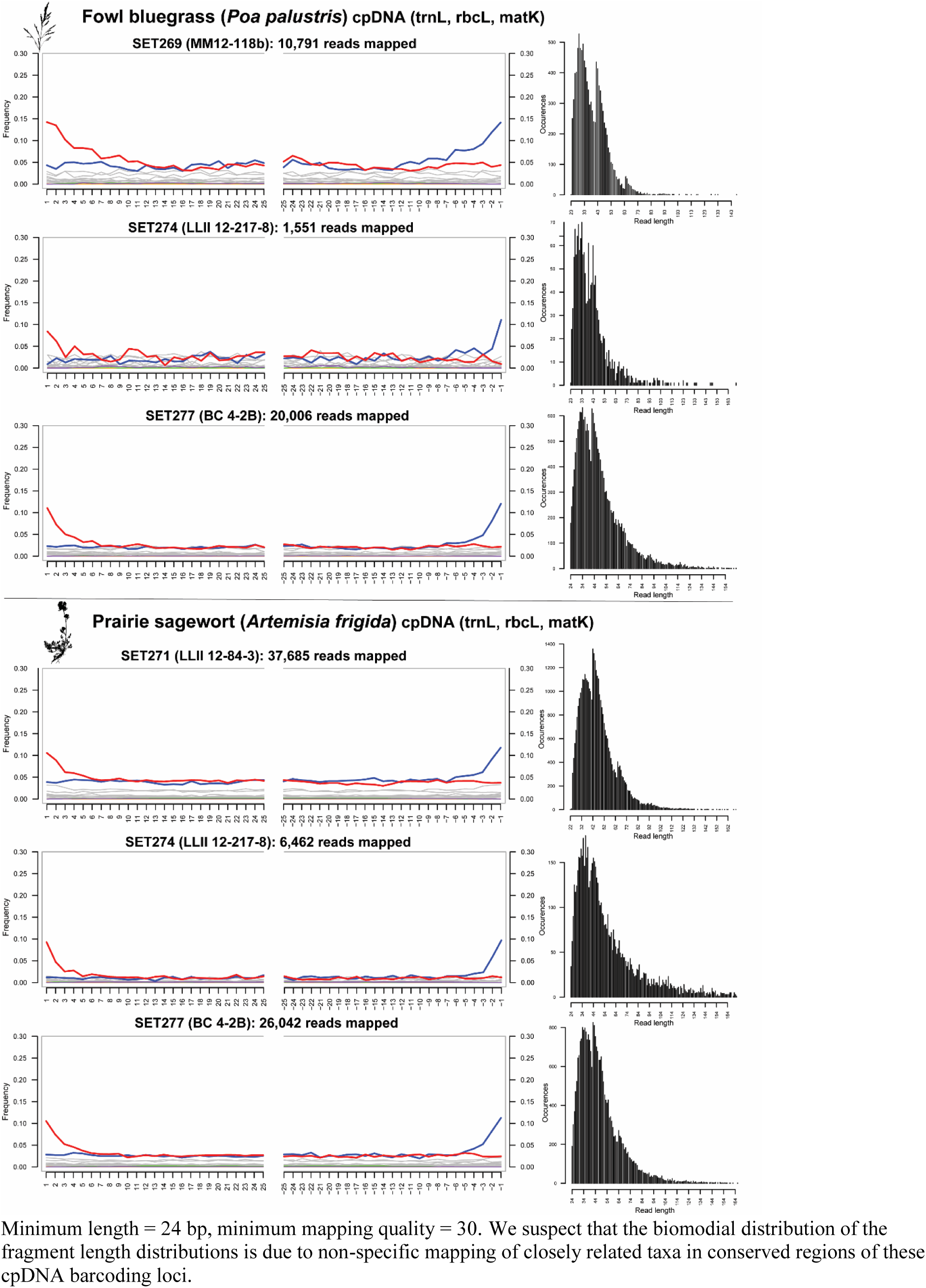
*MapDamage* plots for *Poa palustris and Artemisia figida*. Minimum length = 24 bp, minimum mapping quality = 30. We suspect that the biomodial distribution of the fragment length distributions is due to non-specific mapping of closely related taxa in conserved regions of these cpDNA barcoding loci.

**Figure E11.**
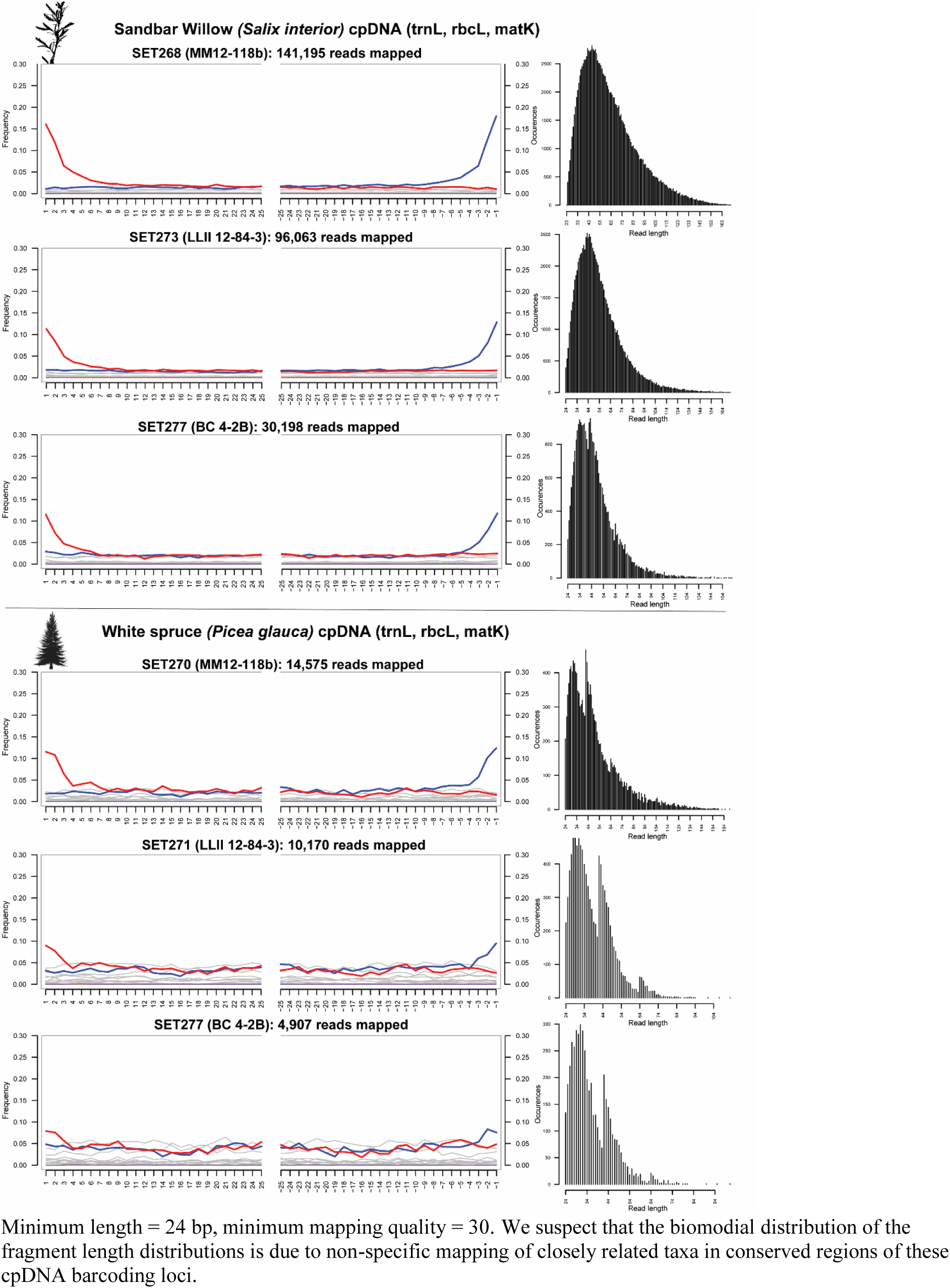
*MapDamage* plots for *Salix interior* and *Picea glauca*. Minimum length = 24 bp, minimum mapping quality = 30. We suspect that the biomodial distribution of the fragment length distributions is due to non-specific mapping of closely related taxa in conserved regions of these cpDNA barcoding loci.

**Figure E12.**
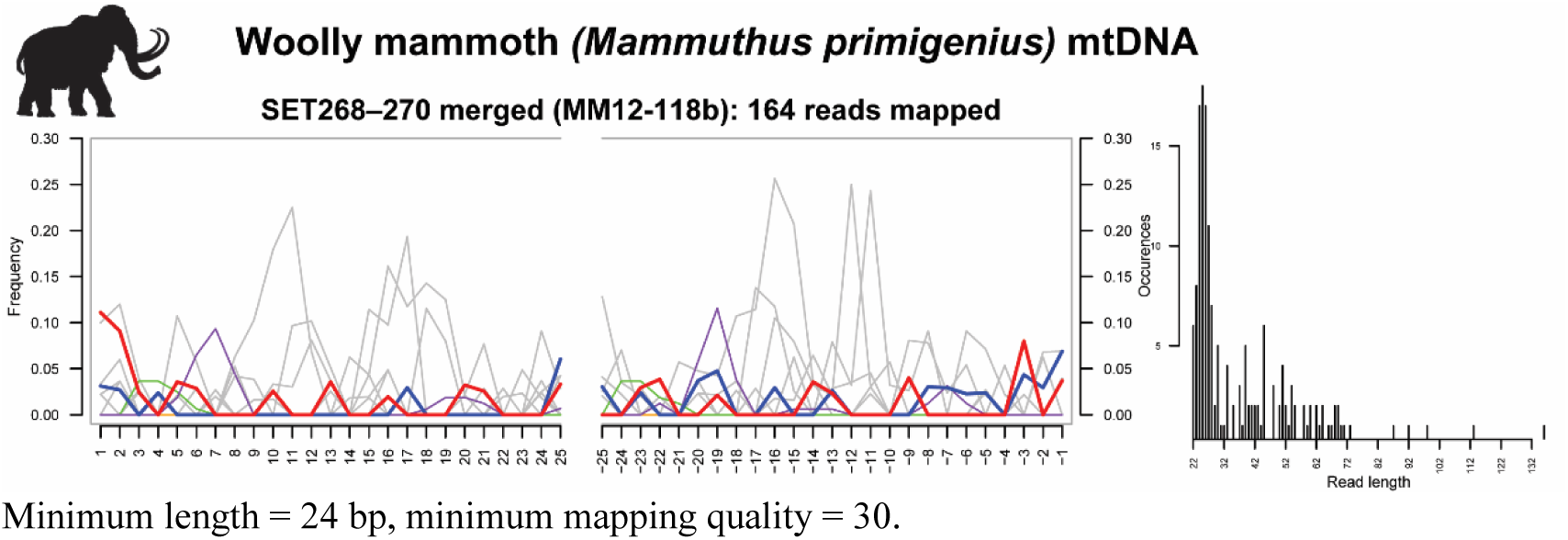
*MapDamage* MM12-118b merged replicates plot for *Mammuthus primigenius*. Minimum length = 24 bp, minimum mapping quality = 30.

**Figure E13.**
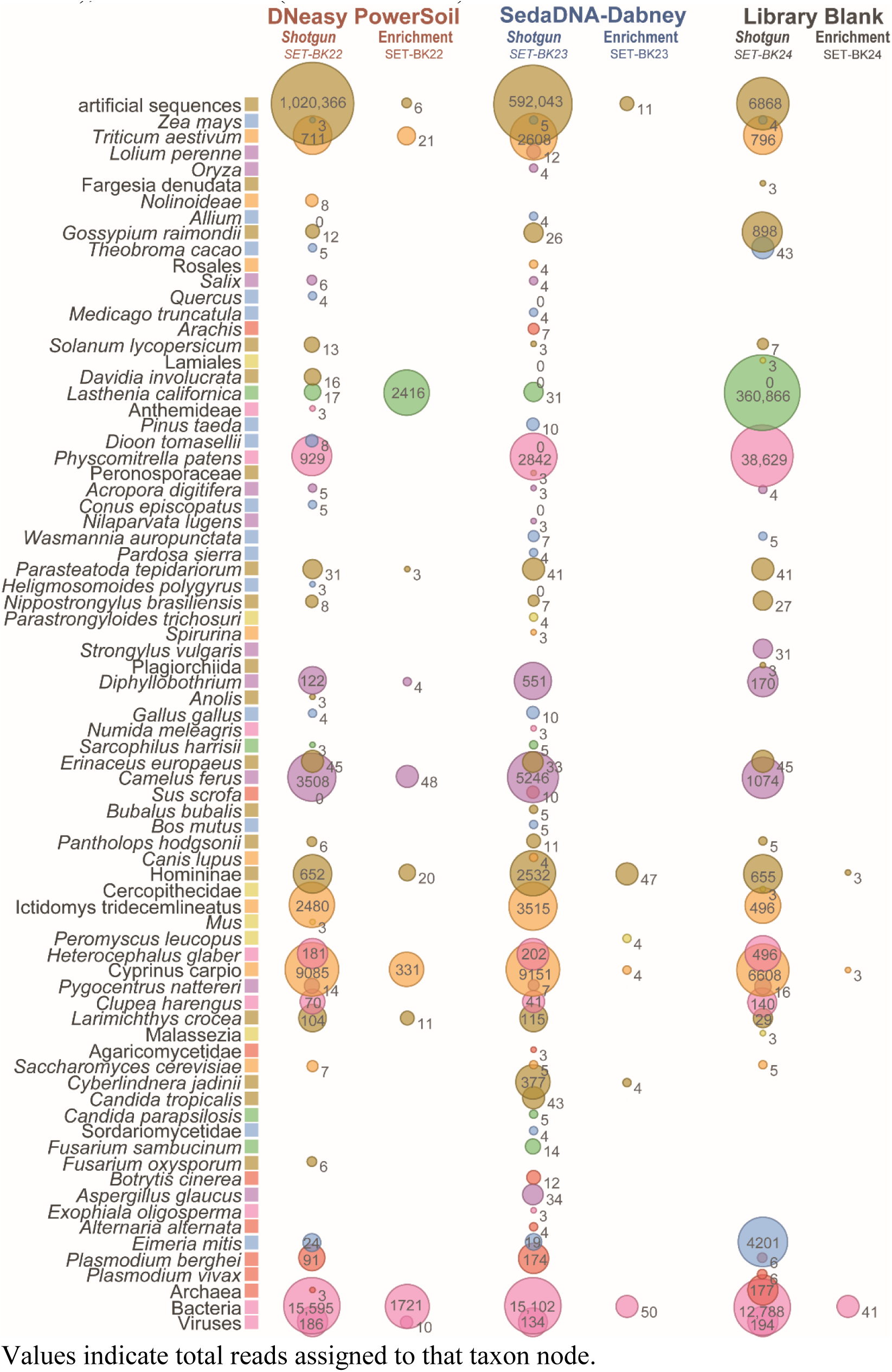
Metagenomic comparison of extraction and library blanks, all reads (not map-filtered), absolute counts (non-normalized). Values indicate total reads assigned to that taxon node.

**Figure E14.**
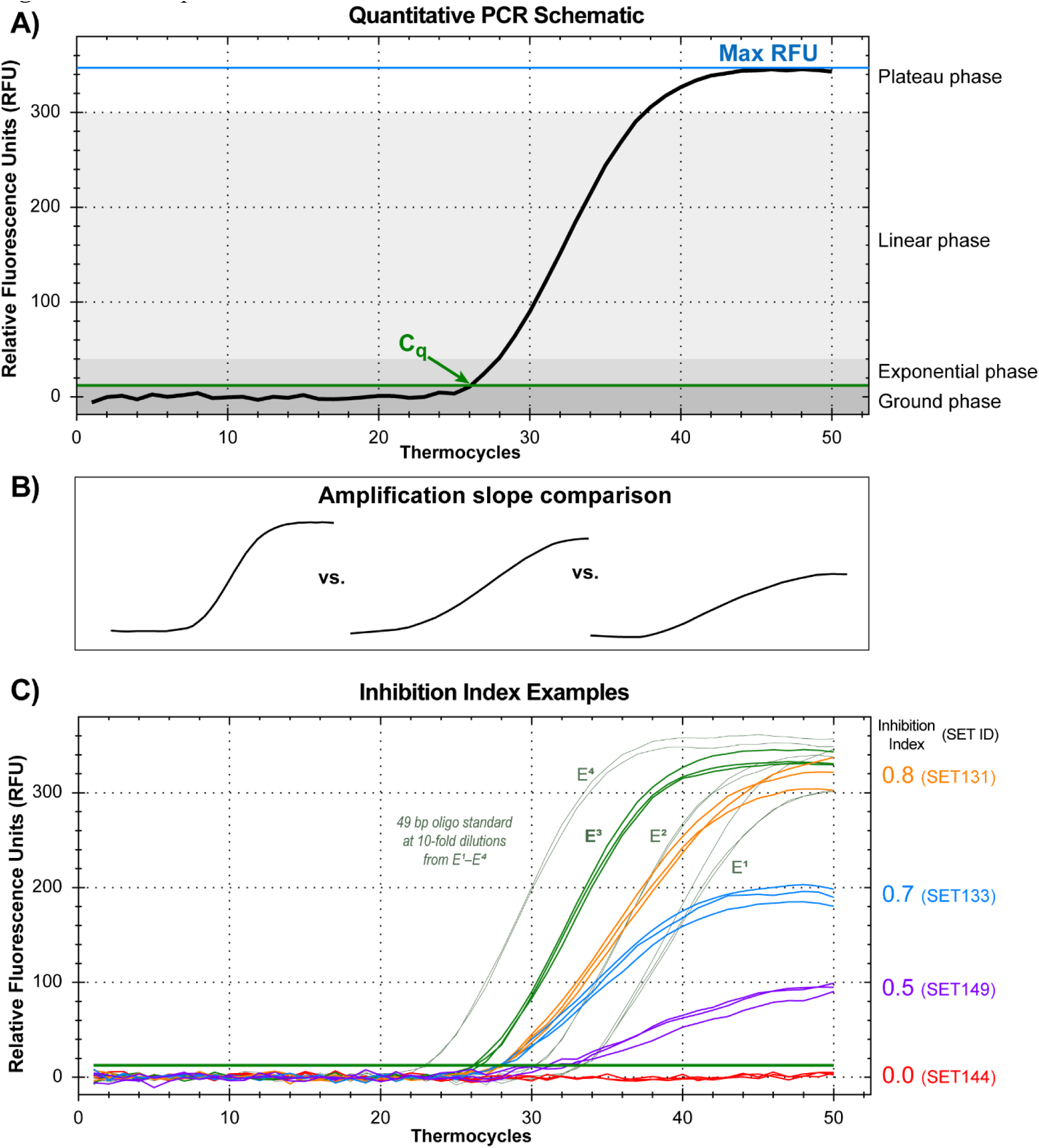
Components of the inhibition index. **A)** A standard qPCR reaction showing C_q_ and max RFU. **B)** A comparison of various amplification slopes from a typical reaction (left), towards increasingly inhibited reactions (right). **C)** Example inhibition indices derived from averaging the C_q_, max RFU, and by fitting a variable-slope sigmoidal dose-response curve to the raw fluorescence data (using GraphPad Prism v. 7.04) based on King et al.(2009) for each PCR replicate by sample against the spiked E^3^ standard. Inhibition index values <0.5 tend to occur when individual PCR replicates fail in a triplicate series; blanks and standard serial dilutions E^2^ and E^1^ tend to have inhibition indices >0.9 despite their 10- and 100-fold reduction in starting DNA causing a 3 or 6 cycle C_q_ shift. QPCR standard curve: E = 94.2%, R^2^ = 0.997, slope = −3.469. See Table S7 for PCR assay specifications.

