## Supplementary material for "PalaeoChip Arctic1.0: An optimised eDNA targeted enrichment approach to reconstructing past environments": Optimization experiments and supplemental materials to manuscript

Each of the following sets of experiments followed the same protocols for subsampling, library preparation, indexing, qPCR inhibition spike tests, and qPCR total quantifications as described in the main paper. Master mix concentrations for each of the aforementioned reactions can be found in Tables S1–S9. Variations in extraction protocols for testing inhibition cleanup techniques are detailed with the description of that experimental SET below, from SET-A to SET-$D_2$. The following table of contents outlines the sections of this appendix.

### SET-A: Initial explorations

Our first set of experiments was intended to determine the best sedaDNA extraction strategy to compare shotgun and targeted enrichment sequence strategies with previously sequenced PCR metabarcoding data collected by Sadoway (2014) on four Yukon sediment cores. However, SET-A informed us that inhibition was a substantial problem with these sediments using our typical in-house demineralization-digestion and Dabney et al. (2013) extraction protocol (hereafter referred to as Dabney), whereas the DNeasy PowerSoil DNA Extraction Kit (hereafter referred to as PowerSoil) only successfully recovered sedaDNA in one of the four cores (with a successful positive control amplification). We felt that this necessitated further experimentation to see if we could overcome enzymatic inhibition and library adapt our target environmental DNA (eDNA) molecules. Permafrost core disks from two strata at Lucky Lady II as well as Bear Creek and Upper Goldbottom Creek (Figure 1, main text) were tested to compare a kit-based sediment DNA extraction strategy (PowerSoil) with our in-house Dabney extraction method.

*SET-A. Extraction*

Samples processed with PowerSoil were extracted following manufacturer specifications. Samples processed using Dabney were first subjected to a two-stage lysis buffer: 1) samples were demineralized in 1 mL of 0.5M EDTA, then rotated continuously for 18 hours at 25°C, 2) samples were then spun down, supernatants were removed, and a proteinase K buffer (Table S1) was added to the sediments to digest overnight at 25°C. The demineralization-digestion supernatants were extracted using a high-volume binding buffer and silica columns following Dabney et al.(2013).

*SET-A. Bioanalyer, inhibitor clean-up, and indexing*

Straight and 1/10 diluted extracts were run on an Agilent 2100 Bioanalyzer as a high sensitivity DNA assay (see Figure S1 and Figure S2). Straight extracts from Dabney samples were darkly coloured and failed to produce a detectable DNA signal on the Bioanalyzer (the baseline was unable to be determined), which we suspected was indicative of abundant co-eluted substances that would adversely affect library preparation. We tested whether a 1/10 dilution or additional purification step prior to blunt end repair would sufficiently remove inhibitors for library preparation in the Dabney extracts, or whether the straight uninhibited PowerSoil extracts would perform better (Table S12). In a qPCR indexing reaction (Table S6) following double-stranded library preparation (DsLp) (Kircher et al., 2012; Meyer and Kircher, 2010), only the positive controls and a single PowerSoil extract from Bear Creek clearly amplified (see Figure S3) despite all samples previously producing positive amplifications and sequence data with a PCR metabarcoding approach (Sadoway, 2014). We suspected that the Dabney extracts were highly inhibited rather than lacking in endogenous DNA. We also suspected that while the PowerSoil kit was effective at removing sedimentary inhibitors, it was ineffective at retaining the kinds of low abundance and highly degraded molecules characteristic of ancient DNA (aDNA).

*SET-A. Inhibition spike tests, qPCR, and DsLp*

In SET-A, a subset of straight extracts, 1/10 and 1/100 diluted extracts, as well as library adapted samples were spiked with the $E^3$ 49 bp standard (see the section "Methods – QPCR:…" in the main text for a description of this spike test). Straight and 1/10 extracts prepared with Dabney were completely inhibited (see Figure S4). Even library adapted samples that had two purification steps and 1/100 diluted extracts were partially inhibited during the inhibition spike test. To determine whether these inhibitors were causing library preparation to fail, we spiked our positive control into each of the Dabney extracts and brought the samples through DsLp. The only samples with positive indexing qPCR amplifications were the positive control and the partially inhibited reaction from LLII 12-127-8 (see Figure S5). All other spiked reactions flatlined, which indicates that inhibition is a significant problem when attempting to bring these extracts into libraries using our in-house lysis and extraction techniques. Compared with the kit however, DNeasy PowerSoil was only sporadically successful at retaining sedaDNA despite all of these core samples previously having been found to contain ancient environmental DNA with PCR metabarcoding (D'Costa et al., 2011; Sadoway, 2014). Our follow-up set of experiments was designed to test various inhibitor removal treatments from the PowerSoil kit and other associated methods to minimize inhibition while maximizing the retention of sedaDNA with our in-house extraction protocols.

**SET-B: Comparing inhibition removal strategies**

The second set of experiments tested the following inhibitor removal augmentations to a digestion and Dabney extraction protocol:

1) Physical disruption with PowerSoil beads (vortexing for 10 minutes) and a proteinase K digestion buffer (see Table S1) (proteinase K was added to each sample individually after vortexing so as to not damage the enzyme, but prior to overnight incubation with continuous oscillation) to release bound DNA.
2) The addition of solution C3 (120 mM aluminum ammonium sulfate dodecahydrate) from the PowerSoil kit prior to Dabney purifications to precipitate inhibitors (maintaining the 1/3 volumes of solution C3 to digest supernatant).
3) A 1 hour 4°C centrifuge at 3900 x *g* with the high-volume Dabney binding buffer in 50 mL tubes to precipitate inhibitors prior to DNA isolation.
4) Sonication with and without a post-sonication purification to disrupt bonds between inhibitors and 'endogenous' sedaDNA Table S14.

Each combination of these inhibitor removal treatments was used on three cores, subsampled from homogenized triplicates. See Table S13 for the SET-B sample list. The extracts were assayed with a qPCR inhibition spike test (Table S7) then brought through DsLp and quantified using P5/P7 adapter primers (Table S8). A subset of these were also indexed to confirm that the qPCR indexing reaction correlates with our short amplification observations (Figure S8).

*SET-B. Results and interpretations*

The 4°C centrifuge samples (and sonicated derivatives) outperformed other inhibitor removal treatments in terms of DNA retention (Figure S6). All other treatment variations had low DNA retention, but frequently outperformed the 1 hour 4°C centrifuge variant in the inhibition spike test. Sonication and post-sonication purifications (to re-concentrate DNA) seemed to help with reducing polymerase inhibition in the qPCR spike test, but also resulted in DNA loss compared to their non-sonicated counterparts. The use of PowerSoil beads for physical disruption resulted in visually clearer extracts and less inhibition than an EDTA based demineralization. Overall, the more treatments utilized, the less inhibition observed, but also the less DNA retained. Solution C3 from the PowerSoil kit was effective at reducing inhibition (both visually in terms of eluate colour retention and in the subsequent inhibition assay), but also resulted in substantial DNA loss. This may explain the results of SET1 where the PowerSoil kit was observed to be effective at removing DNA inhibition but had low DNA retention. The kit is effective with modern sediments and soils, but might precipitate tightly bound organo-mineral complexes (Arnold et al., 2011, p. 418; Haile, 2008, p. 18) in which sedaDNA is preserved. There seems to be an important balance between releasing enough DNA, but not releasing too many inhibitors, as well as removing enough inhibition for enzymatic reactions, while not removing the majority of the ‘endogenous’ sedaDNA (Figure S7). We also wanted to verify that our short amplification assay roughly correlates with amplification during the qPCR indexing reaction (Figure S8). In this assay we can see that sonication results in fewer adaptable molecules during indexing, which is a trend also apparent in the short amplification assay in Figure S6.

**SET-C: Fine-tuning the cold spin**

The only viable treatment from SET-B appears to be the 4°C spin. Our follow-up goal was to determine whether we could maximize the inhibitor removal of the spin at various timings, in this case testing 1, 6, and 19 hours (Table S15). We found that increasing the duration of the 4°C spin reduced the polymerase inhibition observed during the qPCR spike test, which correlated with higher quantifications following library adapter ligation (Figure S9). We also attempted to quantify ‘endogenous’ sedaDNA in extracts prior to adapter ligation using two chloroplast barcoding primer sets, *rbc*L-H1a/H1b (Poinar et al., 1998) and *trn*L P6-loop g/h (Taberlet et al., 2007). However, we found that even in extracts with an inhibition index of ~0.6–0.8 that these assays were uninformative for quantifying pre-DsLp DNA concentrations because the amplification curves were non-standard (the exponential and linear phases had shallower slopes [see Figure E14 in the main text as an example]). This was likely due to both DNA-dependent and -independent inhibition. We found that varying the polymerase concentration dramatically changed the extract quantifications on chloroplast amplicons, which will be elaborated on further in section SET-D.

The final experiment with SET-C samples was to assess whether varying the extract input or enzymatic concentration during blunt-end repair (the first phase of DsLp) would affect the total number of adapter ligated molecules between a highly inhibited sample (BC 4-2B) and an uninhibited sample (LLII 12-217-8) (Figure S10). While there was a reduction in the extraction range of total quantified DNA between replicates of the inhibited core (BC 4-2B) using double the blunt-end repair enzymatic concentrations (T4 polynucleotide kinase and T4 DNA polymerase, see Table S3 for standard concentrations), the effect was marginal. There was also no improvement in halving the extract input in terms of reducing inhibition load on the blunt-end repair enzymes beyond reducing the total adapted DNA by half. Increasing blunt-end repair enzyme concentrations is only recommended for critically important samples where the additional cost is not a concern, as our results suggest that the improvement is marginal.

**SET-D: Optimizing our modified Dabney extraction protocol**

This final experimental set was intended to fine-tune components of our modified Dabney protocol (Table S16). We wanted to determine whether a physical disruption with PowerBeads is necessary, or if a straight digestion or combined demineralization and digestion buffer is best for releasing sedaDNA (SET-B led us to suspect that EDTA might be releasing too many inhibitors). We also wanted to determine whether the duration of the lysis stage was significant, so 6- and 19-hour lysis variants were tested. Finally, we were interested whether increasing the 4°C spin to two days would plateau our inhibition removal procedure, or if we could further improve the amount of library adapted molecules in highly inhibited cores without losing ‘endogenous’ sedaDNA. Part-way though this experiment, we also discovered an interesting (initially, admittedly, very frustrating) variable effect on inhibitor retention linked to the lysis detergent. For the first set of samples (SET-$D_1$) sarkosyl was used as the detergent during lysis instead of SDS. Once this unintended reagent change was discovered (after a series of experiments where the long cold spin was unexpectedly no longer removing inhibition in our most inhibited core sample, BC 4-2B) a second set of homogenized subsamples was taken (SET-$D_2$) for our highly inhibited core where we switched back to SDS as the lysis detergent. We also conducted tests to quantify 'endogenous' chloroplast sedaDNA on extracts (prior to DsLp) from SET-C and -D samples (as alluded to in section SET-C). However, we are unsatisfied with these assays on purified extracts as they still contain co-eluted inhibition (likely both DNA independent and dependent inhibitors), and as such remain unconvinced by their quantifications. Further work developing these specific extract assays (by improving florescence detection, improving inhibition removal, or by size selecting out 'viable' DNA fragments) is recommended.

*SET-D. Results and interpretations*

SET-D experiments yielded five results of interest as observed in Figure S11. First, in column I, there is some indication that increasing the duration of lysis from 9 hours to 19 hours increases inhibitor release (likely as well as DNA release). Inhibitors were not effectively removed during the long cold spin in this instance as sarkosyl appears to be an ineffective detergent to pair with this inhibitor removal technique. This is the only glimpse we observed into the variation of lysis time spans. The optimal interval for our workflow is 19 hours (leaving the samples to oscillate overnight at 35°C). The long cold spin is effective at removing additional inhibition when paired with SDS as observed with columns IV and V in Figure S11. We did not investigate variation in lysis period further as 19 hours is most effective with our workflow and has higher DNA release (as observed in SET-E in the main paper).

Second, the proteinase K buffer with sarkosyl—without either EDTA for demineralization or PowerBeads for physical disruption—has the lowest inhibitor retention (inhibition indices $\gtrsim$0.9) as observed in column II. However, subsequent experiments (detailed in the last SET-D subsection) found that this method also has the worst DNA release.

Third, the detergent used during lysis makes a significant difference in inhibitor retention. All extraction replicates of the inhibited core (BC 4-2B) lysed with the detergent sarkosyl in the proteinase K buffer (with a PowerBead disruption) remain highly inhibited (all failed the inhibition spike test) despite the 4°C inhibition precipitation spin as observed with the inhibition indices in columns III. This is starkly contrasted with column IV where the only experimental change was the use of SDS in the lysing buffer rather than sarkosyl. This was also visually observed in the sarkosyl samples with the lack of a 'dark inhibitor pellet' following the cold spin and much more darkly stained silica-columns and brown-to-black eluates. Our hypothesis for this detergent interaction is detailed in the subsequent SET-D subsection.

Fourth, columns IV and V have equivalent inhibition indices. Subsequent library preparations and short amp quantifications found that increasing the cold spin to 48 hours does slightly increase adapter ligated DNA for both methods (Figure S12). Further, EDTA and PowerBeads as a means of disrupting organo-mineral sedaDNA complexes have roughly equivalent DNA yields, but PowerBeads do release more DNA on average.

Fifth, there appears to be a saturation point for inhibitor removal (at least qPCR polymerase sensitive inhibitors) after approximately 24 hours with the cold spin with our current reagent concentrations. The cold spin at 48 hours show no difference in inhibition indices. This could potentially be modified with higher concentrations of SDS (discussed further in the following subsection on SDS), but this variant was not tested in our optimizations.

*SET-D. SDS and sarkosyl*

The most unexpected result of this project was the effect of unintentionally switching detergents on inhibitor precipitation during the 4°C spin. While there is some degree of precipitation during cold centrifugation with a sarkosyl lysing buffer, or even when spinning low-volume purified extracts at room temperature prior to adapter ligation, the marked increase in precipitation with SDS is visually distinct with thick inhibitor pellets forming during the 4°C spin. SDS (sodium dodecyl sulfate) is an anionic surfactant. Surfactants form self-aggregates (micelles) as their concentration increases (Tanford, 1980). These micelles arrange to have exterior hydrophilic heads, and interior hydrophobic tails. Typically, surfactants are present in submicellar concentrations, but these self-aggregate structures can form at sufficiently high concentrations, particularly with constant mixing. Micelles can also co-aggregate with other amphiphilic compounds (those with hydrophobic and hydrophilic domains) such as humic substances (Koopal et al., 2004; Otto et al., 2003), which we suspect is one of the main DNA independent inhibitors in these sediments due to dark colouration (Alaeddini, 2012). SDS precipitates at 4°C and calcium has been found to increase the precipitation of SDS micelles. We hypothesize that our cold spin with high guanidinium concentrations and SDS based digestion buffer (also containing $CaCl_2$ intended for improving proteinase K efficiency) might have created some form of optimal conditions for micelle formation and the subsequent precipitation of humics and other amphiphilic compounds that bound to SDS micelles. It is also possible that pH is involved in humic acid solubility and is affecting this precipitate reaction (Shaban and Mikulaj, 1998), or that proteins also play some role (Schlager et al., 2012), potentially as related to disentangling sedaDNA from its protective organo-mineral complex (Bezanilla et al., 1995; Blum et al., 1997; Cleaves et al., 2011; Crecchio and Stotzky, 1998; Greaves and Wilson, 1970, 1969; Khanna et al., 2005; Lorenz and Wackernagel, 1987; Ogram et al., 1988; Taylor and Parkinson, 1988). The interaction we have observed here would be of benefit for investigation to further improve the purification of palaeoenvironmental DNA. It is likely that components of the binding or lysing buffers, mixing strategy, or temperature could be tweaked further to improve inhibitor precipitation, thus increasing sedaDNA yield from highly inhibited materials.

*SET-D. Chloroplast assay of 'purified' extracts with variable Taq*

As discussed in section SET-C, we had intended to test 'endogenous' cpDNA retention and release through our procedure before and after library preparation. However, we found that this assay, either with *rbc*L or *trn*L primers, was not a reliable means of assessing 'endogenous' sedaDNA retention through our inhibition removal technique or library adapter ligation efficiency. We found that even our uninhibited core (LLII 12-217-8) had inconsistent DNA quantifications (Figure S13), largely due to non-standard (shallow) amplification slopes affecting the starting quantity metric in our qPCR assay. It is possible that co-eluted humics not removed during the cold-spin may impact florescence detection with these qPCR assays on extracts (Sidstedt et al., 2015), in addition to directly affecting the polymerase. We found substantial quantification increases when doubling *Taq* concentrations (Figure S14). Despite the overall unreliability of these extract assays however, there are two important pieces of information that can be gleaned. First, there is almost no DNA release during lysis when utilizing just a proteinase K digestion buffer with sarkosyl—meaning, without EDTA or physical disruption from PowerSoil beads. This complete lack of amplifiable DNA rules out column II (Figure S11) as a viable extraction strategy. We found the least inhibition with this method-core combination (both in terms of its inhibition indices [Figure S13, Figure S14] and visually non-coloured elutes). Second, despite the qPCR inhibition assay detecting minimal inhibition in this core overall with any method, there is still some sort of inhibition in the extracts affecting the efficiency of AmpliTaq Gold polymerase. This might indicate that while this sample has low DNA independent inhibition (humics and other enzymatically inhibitory substances) that do not impact the amplification of spiked, undamaged synthetic amplicons, this sample likely has high DNA dependent inhibition. Meaning, there might be substantial aDNA damage (such as blocking lesions) or an abundance of extremely short molecules that the *Taq* polymerase is either getting stuck on or stuck amplifying repeatedly, which ultimately leads to poor florescence. This core also had the lowest DNA recovery (see Table 1, main text), which when paired with this assay, suggests that the sedaDNA in this core is more fragmentary and damaged than the other permafrost samples.

Potentially spiking lysing buffers with a known quantity of aDNA size and damage characteristic non-target DNA, then assaying those synthetic molecules after library preparation with a dual adapter/synthetic target-specific primer set (such as described in Table S9), would be a more reliable means of assessing DNA loss from the cold spin and DNA independent inhibitor effects on library preparation efficiency. However, this assay would not assess DNA dependent inhibition specific to the sedaDNA constituents of the sample. The polymerases used in qPCR amplifications are not directly equivalent to those used in library preparation. Extracts or libraries with florescence inhibition (Sidstedt et al., 2015) might yet be amenable to adapter and indexing ligations, as well as potentially sequencing, but be largely undetectable with *Taq* based qPCR assays unless the reaction is maximally saturated with polymerase to mitigate various forms of inhibition. While potentially feasible with small sample-sets, this strategy would be costly, and likely difficult to standardize across highly variable molecular constituents even within the ‘same’ homogenized sedimentary sample (however ‘homogenized’ a sediment sample could be on a molecular scale).

**SET-E: Additional metagenomic comparisons for main-text experiment**

In our attempt to fairly compare the three sedaDNA approaches described in the main paper (while acknowledging that any ‘normalization’ inadvertently introduces new biases), we have generated metagenomic bubble charts with normalized read counts (that had been mapped back to our curated baits), as well as total read metagenomic profiles that have not been map-filtered (with both absolute and normalized count variants). These were generated identically to the metagenomic bubble charts in the main text, except only the top 10 alignments (rather than top 100) were kept for the non-map filtered blasts to reduce unwieldy file sizes. With all normalized charts here, libraries were normalized to the lowest sequenced library in *MEGAN* while maintaining at least one taxon read for reduced libraries. Normalized charts do not contain exact value read counts as these values have been manipulated during normalization, a scale chart is included for relative comparisons. For non map-filtered bubble charts, major prokaryotes are shown. Within the eukaryotes, only chordates and viridiplantae shown (to be able to visualize these comparisons). Additional potentially ‘authentic’ sedaDNA taxa are identified in the non-map filtered (all reads) bubble charts. However, most of these taxa are in the curated baits (such as moose, *Alces alces*). So, either 1) those sequences had been clustered or masked in the curated baits, which is why they did not map during initial filtering, 2) are from non-specific regions that were removed from the baits, or 3) the low top alignments (10 versus 100) resulted in less conservative LCA-assignments when not map-filtering the reads. It is also important to note that some common contaminants and adapter-contaminated genomes on NCBI (e.g. camel, carp, wheat) remain in the non-map filtered metagenomic profiles despite attempts to filter out adaptamers (whereas these problematic hits are filtered out early in processing by mapping to a curated bait-set). This inflates the read counts in the comparisons to some degree, particularly with taxa collapsed at such high ranks to allow for the entire metagenomic profile of each core to be easily visualized. The following list describes the bubble chart metagenomic comparisons included in this appendix.

1. Map-filtered, normalized bubble charts: Figures S15–S18.
2. Non map-filtered, absolute count bubble charts: Figures S19–S23.
3. Non map-filtered, absolute count bubble charts, extraction and library blanks: Figure S23.
4. Non map-filtered, normalized bubble charts: Figures S24–S27

A summation of major clade *MEGAN* LCA-assignments from the main text are also included in Tables S17 and S18 (page 24).

### Tables

**Table S1** Final concentrations of components in the proteinase K digestion solution.

| Proteinase K Digestion Solution | |
|---|---|
| Component | Final Concentration |
| Tris-Cl (pH 9.0) | 0.02 M |
| SDS | 0.5 % |
| Proteinase K | 0.25 mg/ml |
| CaCl2 | 0.01 M |
| DTT | 100 mM |
| PVP | 2.5 % |
| PTB | 5 mM |

Samples were digested overnight at 35°C with rotation. Nanopure Barnstead water was used to bring up the volume to the desired concentration. Concentrations based on Karpinski et al. (2016). For samples where sarkosyl was used instead of SDS, the final detergent concentration was unchanged.

**Table S2** Final concentrations of components in the Dabney binding buffer.

| Dabney Binding Buffer | |
|---|---|
| Component | Final Concentration |
| Guanidine Hydrochloride | 5 M |
| Isopropanol (100%) | 40 % |
| Tween-20 | 0.05 % |
| 3 M Sodium Acetate (pH 5.2) | 0.09 M |

Nanopure Barnstead water was used to bring up the volume to the desired concentration. Concentrations based on Dabney et al. (2013).

**Table S3** Final concentrations of components in the blunt-end repair mixture.

| Blunt-End Repair Mixture | |
|---|---|
| Component | Final Concentration |
| NE Buffer 2.1 | 1X |
| DTT | 1 mM |
| dNTP mix | 100 µM |
| ATP | 1 mM |
| T4 polynucleotide kinase | 0.5 U/µL |
| T4 DNA polymerase | 0.1 U/µL |

A final volume of 40 µL was used for the mixture and template DNA. Nanopure Barnstead water was used to bring up the volume to the desired concentration.

**Table S4** Final concentrations of all components in the adapter ligation mixture.

| 3. Adapter Ligation Mixture | |
|---|---|
| Component | Final Concentration |
| T4 DNA Ligase Buffer | 1X |
| PEG-4000 | 5% |
| Adapter Mix | 0.5 µM |
| T4 DNA Ligase | 0.125 U/µl |
| 2. Adapter Mix | |
| IS1_adapter_P5.F | 200 µM |
| IS2_adapter_P7.F | 200 µM |
| IS3_adapter_P5+P7.R | 200 µM |
| Oligo Hybridization Buffer | 1X |
| 1. Oligo Hybridization Buffer | |
| NaCl | 500 mM |
| Tris-Cl, pH 8.0 | 10 mM |
| EDTA, pH 8.0 | 1 mM |

Oligo Hybridization Buffer was prepared prior to the Adapter Mix, which was prepared separately for IS1_adapter_P5.F and IS2_adapter_P7.F. These two mixes were then combined after an incubation at 95°C for 10 seconds, and a ramp from 95°C to 12°C at a rate of 0.1°C/sec. A final volume of 40 µl was used for the mixture and template DNA. Nanopure Barnstead water (not listed) was used to bring the volume up to the desired concentration.

**Table S5** Final concentrations of components in the adapter fill-in mixture.

| Adapter Fill-In Mixture | |
|---|---|
| Component | Final Concentration |
| ThermoPol Reaction Buffer | 1X |
| dNTP Mix | 250 µM |
| BST Polymerase (large fragment) | 0.4 U/µl |

A final volume of 40 µl was used for the mixture and template DNA with the addition of Nanopure Barnstead water to bring the mix up to the desired concentration and volume.

**Table S6** Primer sequences, PCR master mix, and cycling protocol for indexing amplification.

<table>
<tr><th colspan="4">Indexing PCR Master Mix</th></tr>
<tr><th colspan="2">Component</th><th colspan="2">Final Concentration</th></tr>
<tr><td colspan="2">KAPA SYBR®FAST qPCR Master Mix (2X)</td><td colspan="2">1X</td></tr>
<tr><td colspan="2">Forward primer</td><td colspan="2">750 nM</td></tr>
<tr><td colspan="2">Reverse primer</td><td colspan="2">750 nM</td></tr>
<tr><th colspan="4">Primer Sequences</th></tr>
<tr><td>Forward Primer</td><td colspan="3">AATGATACGGCGACCACCGAGATCTACACNNNNNNNACACTCTTTCCCTACACGACGCTCTT</td></tr>
<tr><td>Reverse Primer</td><td colspan="3">CAAGCAGAAGACGGCATACGAGATTATNNNNNNNACTGGAGTTCAGACGTGT</td></tr>
<tr><th colspan="4">Indexing PCR Protocol</th></tr>
<tr><th>Phase</th><th>Temperature (°C)</th><th>Time</th><th>Cycles</th></tr>
<tr><td>Initial Denaturation</td><td>98</td><td>3 min</td><td></td></tr>
<tr><td>Denaturation</td><td>98</td><td>20 sec</td><td rowspan="3">Repeated for 8-12 cycles</td></tr>
<tr><td>Annealing</td><td>*60*</td><td>*20 sec*</td></tr>
<tr><td>Extension</td><td>72</td><td>25 sec</td></tr>
<tr><td>Final Extension</td><td>72</td><td>3 min</td><td></td></tr>
</table>

The N in each primer sequence represents the 7 bp index specific to each primer. A final reaction volume of 40 µl was used for the assay, with 12.5 µl of the adapter ligated DNA libraries. Nanopure Barnstead water (not listed) was used to bring the volume up to the desired concentration.
Fluorescence readings were recorded post-annealing as indicated above with asterisks.

**Table S7** Inhibition spike test qPCR assay.

| PCR Master Mix | |
|---|---|
| Component | Final Concentration |
| 10X PCR Buffer II | 1X |
| MgCl2 | 2.5 mM |
| dNTP mix | 250 µM |
| BSA | 1 mg/ml |
| Forward primer (971) | 0.25 µM |
| Reverse primer (1040) | 0.25 µM |
| EvaGreen | 0.5X |
| AmpliTaq Gold | 0.05 U/µL |
| Oligo | Sequence (5'–3') |
| Forward primer (971_Mamm_Fwd) | CCCTAAACTTTGATAGCTACC |
| Reverse primer (1040_Mamm_Rev) | GTAGTTCTCTGGCGGATAGC |
| Double stranded 49 bp amplicon based on the mammoth *12S* mitochondrial gene | ACACTCTTTCCCTACACGACGC<br>TCTTCCGATCTCCCTAAACTTT<br>GATAGCTACCTTTACAAAGCTA<br>TCCGCCAGAGAACTACAGATC<br>GGAAGAGCACACGTCTGAACT<br>CCAGTCAC |
| Input* | Volume |
| PCR master mix | 8 µL |
| sedaDNA extract template | 1 µL |
| 49 bp amplicon spike | 1 µL |

| PCR Protocol | | | |
|---|---|---|---|
| Phase | Temperature (°C) | Time | Cycles |
| Initial Denaturation | 95 | 5 min | |
| Denaturation | 95 | 30 sec | Repeated for 50 cycles |
| Annealing | 54 | 30 sec | |
| Extension | **72** | 50 sec | |
| Final Extension | 72 | 1 min | |
| Melt Curve | **55–95** | **5 sec per degree** | |

*Sample wells = 1 µL template + 1 µL spike. QPCR standard wells = 1 µL spike, 1 µL 0.1X TE. Non-template controls = 2 µL 0.1X TE.
**Fluorescence readings were recorded post-annealing and during the melt curve as indicated above with asterisks.
Assay from Enk et al. (2016).
Nanopure Barnstead water was used to bring the master mix up to the desired concentration and volume.

**Table S8** Library adapted short amp total quantification PCR.

| PCR Master Mix | |
|---|---|
| **Component** | **Final Concentration** |
| KAPA SYBR®FAST qPCR Master Mix (2X) | 1X |
| Forward primer | 0.2 µM |
| Reverse primer | 0.2 µM |
| **Oligos** | **Sequence (5'–3')** |
| Forward primer (ILPr_shortampP5F_MeyerIS7) | ACACTCTTTCCCTACACGAC |
| Reverse primer (ILPr_shortampP7R_MeyerIS8) | GTGACTGGAGTTCAGACGTGT |
| Library adapted oligo based on the mammoth *12S* mitochondrial gene | GTGACACTCTTTCCCTACACGACTGGGCAATC<br>CTGAGCCAAATGATATGATTTGAGATATTGA<br>TAGAATTGAATGCATAGTGATAAAAGGATGA<br>TATATTAGGATAGGTGCAGAGACTCAATGGA<br>ACACGTCTGAACTCCAGTCACGTA |
| **Input** | **Volume** |
| PCR master mix | 6 µL |
| Library adapted template | 4 µL |

| PCR Protocol | | | |
|---|---|---|---|
| **Phase** | **Temperature (ºC)** | **Time** | **Cycles** |
| Initial Denaturation | 95 | 5 min | |
| Denaturation | 95 | 30 sec | Repeated for 30 cycles |
| Annealing + Extension | 60 | 45 sec | |
| Melt Curve | **65–95** | **5 sec per degree** | |

Nanopure Barnstead water was used to bring the mix up to the desired concentration and volume. Oligo based on Enk et al. (2016); primers based Meyer and Kircher (2010).

**Table S9** Library adapted *trnL* short amp total quantification PCR.

| PCR Master Mix | |
|---|---|
| **Component** | **Final Concentration** |
| 10X PCR Buffer II | 1X |
| MgCl2 | 2.5 mM |
| dNTP mix | 250 µM |
| BSA | 1 mg/ml |
| Forward primer | 0.25 µM |
| Reverse primer | 0.75 µM |
| EvaGreen | 0.5X |
| AmpliTaq Gold | 0.05 U/µL |
| **Oligos** | **Sequence (5'–3')** |
| Forward primer (trnL_P6-g_F) | GGGCAATCCTGAGCCAA |
| Reverse primer (ILPr_shortampP7R_MeyerIS8) | GTGACTGGAGTTCAGACGTGT |
| Oligo with binding sites for library adapter primers and *trnL* primers from Taberlet et al. (2007). Oligo insert shows no significant similarity with blastn and a top blast hit to Staphylococcus aureus with an E-value of 0.056 using megablast at the time of publication. | GGGCAATCCTGAGCCAAATGATATGA<br>TTTGAGATATTGATAGAATTGAATGC<br>ATAGTGATAAAAGGATGATATATTAG<br>GATAGGTGCAGAGACTCAATGGAACA<br>CGTCTGAACTCCAGTCAC |
| **Input** | **Volume** |
| PCR master mix | 8 µL |
| Library adapted template | 1 µL |

| PCR Protocol | | | |
|---|---|---|---|
| **Phase** | **Temperature (°C)** | **Time** | **Cycles** |
| Initial Denaturation | 95 | 5 min | |
| Denaturation | 95 | 30 sec | Repeated for 50 cycles |
| Annealing | 51 | 30 sec | |
| Extension | *72* | 50 sec | |
| Final Extension | 72 | 1 min | |
| Melt Curve | *55–95* | *1 sec per degree* | |

*Fluorescence readings were recorded post-annealing and during the melt curve as indicated above with asterisks.

Nanopure Barnstead water was used to bring the master mix up to the desired concentration and volume.

Library adapter primer based on Meyer and Kircher (2010); primer *trnL*-g targets the P6 loop of the *trnL* cpDNA intron, and is based on Taberlet et al. (2007).

**Table S10** Library adapted and indexed long amp total quantification PCR.

| PCR Master Mix | |
|---|---|
| **Component** | **Final Concentration** |
| KAPA SYBR®FAST qPCR Master Mix (2X) | 1X |
| Forward primer | 0.2 µM |
| Reverse primer | 0.2 µM |
| **Oligos** | **Sequence (5'–3')** |
| Forward primer (ILPr_shortampP5F_MeyerIS5) | AATGATACGGCGACCACCGA |
| Reverse primer (ILPr_shortampP7R_MeyerIS6) | CAAGCAGAAGACGGCATACGA |
| PhiX library adapted control standard from 100 pM to 62.6 fM | AATGATACGGCGACCACCGA<br>*ADAPTER INSERT*<br>TCGTATGCCGTCTTCTGCTTG |
| **Input** | **Volume** |
| PCR master mix | 6 µL |
| Library adapted and indexed template | 4 µL |

| PCR Protocol | | | |
|---|---|---|---|
| **Phase** | **Temperature (ºC)** | **Time** | **Cycles** |
| Initial Denaturation | 95 | 5 min | 1 |
| Denaturation | 95 | 30 sec | Repeated for 35 cycles |
| Annealing + Extension | 60 | 45 sec | |
| Cooldown | 8 | 30 sec | 1 |

Nanopure Barnstead water was used to bring the mix up to the desired concentration and volume. Primers from Meyer and Kircher (2010).

**Table S11** Enrichment mastermixes.

| Hybridization MasterMix | |
|---|---|
| Component | Final Concentration |
| Hyb N (19.46X SSPE, 13.5 mM EDTA) | 9X, 6.25mM |
| Hyb D (50X Denhardt's Solution) | 8.75X |
| Hyb S (10% SDS) | 0.25% |
| Hyb R RNAsecure | 1.56X |
| Bait Mixture (200 ng baits per reaction) | 11.11 ng/µL |
| Bait Mixture | |
| Component | Final Concentration |
| Plant: 18,672 baits | 83.33 ng/rxn |
| Animal: 57,588 baits | 138.89 ng/rxn |
| Library MasterMix | |
| Component | Final Concentration |
| Block A (Illumina bloligos xGens) | 0.04 ng/µL |
| Block C (Human COt-1 DNA) | 0.19 ng/µL |
| Block O (Salmon Sperm DNA) | 0.19 ng/µL |
| Library template input | 7 µL |
| Wash Buffer X (0.2X WB) | |
| Component | Final Concentration |
| HYB S (10% SDS) | 0.08 % |
| Wash Buffer<br>(0.1X SSC; 0.1% SDS; 1mM EDTA) | 0.2X |

Nanopure Barnstead water was used to bring mixes up to the desired concentration and volume.

**Table S12** SET-A sample list.

| SET ID | Extraction method | Extract clean-up prior to DsLp | Core/sample | Previous ID | Site | Sample Type |
|---|---|---|---|---|---|---|
| SET2 | PowerSoil | | MM12-118b | GB1 | Upper Gold Bottom Creek | Permafrost |
| SET4 | DD-Dabney | | MM12-118b | GB1 | Upper Gold Bottom Creek | Permafrost |
| SET5 | DD-Dabney | 1/10 dilution | MM12-118b | GB1 | Upper Gold Bottom Creek | Permafrost |
| SET6 | DD-Dabney | QiaQuick Purification | MM12-118b | GB1 | Upper Gold Bottom Creek | Permafrost |
| SET9 | PowerSoil | | LLII 12-84-3 | LL3 | Lucky Lady II | Permafrost |
| SET10 | PowerSoil | 1/10 dilution | LLII 12-84-3 | LL3 | Lucky Lady II | Permafrost |
| SET13 | DD-Dabney | | LLII 12-84-3 | LL3 | Lucky Lady II | Permafrost |
| SET14 | DD-Dabney | 1/10 dilution | LLII 12-84-3 | LL3 | Lucky Lady II | Permafrost |
| SET15 | DD-Dabney | QiaQuick purification | LLII 12-84-3 | LL3 | Lucky Lady II | Permafrost |
| SET17 | PowerSoil | | LLII 12-217-8 | LL1 | Lucky Lady II | Permafrost |
| SET19 | DD-Dabney | | LLII 12-217-8 | LL1 | Lucky Lady II | Permafrost |
| SET20 | DD-Dabney | 1/10 dilution | LLII 12-217-8 | LL1 | Lucky Lady II | Permafrost |
| SET21 | DD-Dabney | QiaQuick purification | LLII 12-217-8 | LL1 | Lucky Lady II | Permafrost |
| SET23 | PowerSoil | | BC 4-2B | BC | Bear Creek | Permafrost |
| SET25 | DD-Dabney | | BC 4-2B | BC | Bear Creek | Permafrost |
| SET26 | DD-Dabney | 1/10 dilution | BC 4-2B | BC | Bear Creek | Permafrost |
| SET27 | DD-Dabney | QiaQuick purification | BC 4-2B | BC | Bear Creek | Permafrost |
| SETPC1 | PowerSoil | | *N. shastensis* | 089 | Gypsum Cave, Nevada | Palaeofeces |
| SETPC2 | DD-Dabney | | *N. shastensis* | 089 | Gypsum Cave, Nevada | Palaeofeces |
| SETBK1 | PowerSoil | **Extraction Blanks** | | | | |
| SETBK2 | DD-Dabney | | | | | |

Core/previous ID as per Sadoway (2014). All sediment cores from the Yukon.
PowerSoil: DNeasy PowerSoil extraction kit.
DD-Dabney: a two-stage demineralization (0.5 M EDTA) and digestion (proteinase K buffer, see Table S1) (each overnight) followed by purification with a high-volume binding buffer and Roche Diagnostics silica-spin column following Dabney et al.(2013).
DsLp: Double-stranded library preparation (Kircher et al., 2012; Meyer and Kircher, 2010).
PC1/2: Positive control 089, *Nothrotheriops shastensis (*Shasta ground sloth) palaeofeces (Poinar et al., 1998).
Pre-DsLp clean-up with a QiaQuick PCR Purification Kit, or the extract was diluted to 1/10 prior to DsLp.
Observations: Dabney extracts without an additional clean-up were very darkly coloured compared to the clear PowerSoil extracts.

**Table S13** SET-B sample list.

| Sample Information | | | | | Treatment | | |
|---|---|---|---|---|---|---|---|
| SET ID | Extract ID | Core | Previous ID | Sample Type | PowerBeads | Solution C3 | 4°C Spin |
| **SET28** | D7a | LLII 12-84-3 | LL3 | Pf | Y | - | - |
| **SET29** | D8a | LLII 12-84-3 | LL3 | Pf | Y | - | Y |
| **SET30** | D9a | LLII 12-84-3 | LL3 | Pf | Y | Y | - |
| **SET31** | D10a | LLII 12-84-3 | LL3 | Pf | Y | Y | Y |
| **SET32** | D11a | LLII 12-84-3 | LL3 | Pf | -DD | Y | Y |
| **SET33** | D7b | LLII 12-84-3 | LL3 | Pf | Y | - | - |
| **SET34** | D8b | LLII 12-84-3 | LL3 | Pf | Y | - | Y |
| **SET35** | D9b | LLII 12-84-3 | LL3 | Pf | Y | Y | - |
| **SET36** | D10b | LLII 12-84-3 | LL3 | Pf | Y | Y | Y |
| **SET37** | D11b | LLII 12-84-3 | LL3 | Pf | -Dig | Y | Y |
| **SET38** | D7c | LLII 12-84-3 | LL3 | Pf | Y | - | - |
| **SET39** | D8c | LLII 12-84-3 | LL3 | Pf | Y | - | Y |
| **SET40** | D9c | LLII 12-84-3 | LL3 | Pf | Y | Y | - |
| **SET41** | D10c | LLII 12-84-3 | LL3 | Pf | Y | Y | Y |
| **SET43** | D12a | LLII 12-217-8 | LL1 | Pf | Y | - | - |
| **SET44** | D13a | LLII 12-217-8 | LL1 | Pf | Y | - | Y |
| **SET45** | D14a | LLII 12-217-8 | LL1 | Pf | Y | Y | - |
| **SET46** | D15a | LLII 12-217-8 | LL1 | Pf | Y | Y | Y |
| **SET47** | D16a | LLII 12-217-8 | LL1 | Pf | -DD | Y | Y |
| **SET48** | D12b | LLII 12-217-8 | LL1 | Pf | Y | - | - |
| **SET49** | D13b | LLII 12-217-8 | LL1 | Pf | Y | - | Y |
| **SET50** | D14b | LLII 12-217-8 | LL1 | Pf | Y | Y | - |
| **SET51** | D15b | LLII 12-217-8 | LL1 | Pf | Y | Y | Y |
| **SET52** | D16b | LLII 12-217-8 | LL1 | Pf | -Dig | Y | Y |
| **SET53** | D12c | LLII 12-217-8 | LL1 | Pf | Y | - | - |
| **SET54** | D13c | LLII 12-217-8 | LL1 | Pf | Y | - | Y |
| **SET55** | D14c | LLII 12-217-8 | LL1 | Pf | Y | Y | - |
| **SET56** | D15c | LLII 12-217-8 | LL1 | Pf | Y | Y | Y |
| **SET58** | D17a | BC 4-2B | BC | Pf | Y | - | - |
| **SET59** | D18a | BC 4-2B | BC | Pf | Y | - | Y |
| **SET60** | D19a | BC 4-2B | BC | Pf | Y | Y | - |
| **SET61** | D20a | BC 4-2B | BC | Pf | Y | Y | Y |
| **SET62** | D21a | BC 4-2B | BC | Pf | -DD | Y | Y |
| **SET63** | D17b | BC 4-2B | BC | Pf | Y | - | - |
| **SET64** | D18b | BC 4-2B | BC | Pf | Y | - | Y |
| **SET65** | D19b | BC 4-2B | BC | Pf | Y | Y | - |
| **SET66** | D20b | BC 4-2B | BC | Pf | Y | Y | Y |
| **SET67** | D21b | BC 4-2B | BC | Pf | -Dig | Y | Y |
| **SET68** | D17c | BC 4-2B | BC | Pf | Y | - | - |
| **SET69** | D18c | BC 4-2B | BC | Pf | Y | - | Y |
| **SET70** | D19c | BC 4-2B | BC | Pf | Y | Y | - |
| **SET71** | D20c | BC 4-2B | BC | Pf | Y | Y | Y |
| **SETBK3** | D23 | Extraction blank | | | -DD | - | - |
| **SETBK4** | D24 | Extraction blank | | | Y | Y | Y |
| **SETBK5** | D25 | Extraction blank | | | Y | Y | Y |

| Sample Information | | | | | Treatment | | | | |
|---|---|---|---|---|---|---|---|---|---|
| SET ID | Extract ID | Core | Previous ID | Sample Type | PowerBeads | Solution C3 | 4°C Spin | Sonication | Post-sonication purification |
| **SET73** | D7a | LLII 12-84-3 | LL3 | Pf | Y | - | - | Y | Y |
| **SET74** | D8a | LLII 12-84-3 | LL3 | Pf | Y | - | Y | Y | Y |
| **SET75** | D9a | LLII 12-84-3 | LL3 | Pf | Y | Y | - | Y | Y |
| **SET76** | D10a | LLII 12-84-3 | LL3 | Pf | Y | Y | Y | Y | Y |
| **SET77** | D11a | LLII 12-84-3 | LL3 | Pf | -DD | Y | Y | Y | Y |
| **SET78** | D7b | LLII 12-84-3 | LL3 | Pf | Y | - | - | Y | - |
| **SET79** | D8b | LLII 12-84-3 | LL3 | Pf | Y | - | Y | Y | - |
| **SET80** | D9b | LLII 12-84-3 | LL3 | Pf | Y | Y | - | Y | - |
| **SET81** | D10b | LLII 12-84-3 | LL3 | Pf | Y | Y | Y | Y | - |
| **SET82** | D11b | LLII 12-84-3 | LL3 | Pf | -Dig | Y | Y | Y | - |
| **SET83** | D7c | LLII 12-84-3 | LL3 | Pf | Y | - | - | Y | Y |
| **SET84** | D8c | LLII 12-84-3 | LL3 | Pf | Y | - | Y | Y | - |
| **SET85** | D9c | LLII 12-84-3 | LL3 | Pf | Y | Y | - | Y | - |
| **SET86** | D10c | LLII 12-84-3 | LL3 | Pf | Y | Y | Y | Y | Y |
| **SET88** | D12a | LLII 12-217-8 | LL1 | Pf | Y | - | - | Y | Y |
| **SET89** | D13a | LLII 12-217-8 | LL1 | Pf | Y | - | Y | Y | Y |
| **SET90** | D14a | LLII 12-217-8 | LL1 | Pf | Y | Y | - | Y | Y |
| **SET91** | D15a | LLII 12-217-8 | LL1 | Pf | Y | Y | Y | Y | Y |
| **SET92** | D16a | LLII 12-217-8 | LL1 | Pf | -DD | Y | Y | Y | Y |
| **SET93** | D12b | LLII 12-217-8 | LL1 | Pf | Y | - | - | Y | - |
| **SET94** | D13b | LLII 12-217-8 | LL1 | Pf | Y | - | Y | Y | - |
| **SET95** | D14b | LLII 12-217-8 | LL1 | Pf | Y | Y | - | Y | - |
| **SET96** | D15b | LLII 12-217-8 | LL1 | Pf | Y | Y | Y | Y | - |
| **SET97** | D16b | LLII 12-217-8 | LL1 | Pf | -Dig | Y | Y | Y | - |
| **SET98** | D12c | LLII 12-217-8 | LL1 | Pf | Y | - | - | Y | Y |
| **SET99** | D13c | LLII 12-217-8 | LL1 | Pf | Y | - | Y | Y | - |
| **SET100** | D14c | LLII 12-217-8 | LL1 | Pf | Y | Y | - | Y | - |
| **SET101** | D15c | LLII 12-217-8 | LL1 | Pf | Y | Y | Y | Y | Y |
| **SET103** | D17a | BC 4-2B | BC | Pf | Y | - | - | Y | Y |
| **SET104** | D18a | BC 4-2B | BC | Pf | Y | - | Y | Y | Y |
| **SET105** | D19a | BC 4-2B | BC | Pf | Y | Y | - | Y | Y |
| **SET106** | D20a | BC 4-2B | BC | Pf | Y | Y | Y | Y | Y |
| **SET107** | D21a | BC 4-2B | BC | Pf | -DD | Y | Y | Y | Y |
| **SET108** | D17b | BC 4-2B | BC | Pf | Y | - | - | Y | - |
| **SET109** | D18b | BC 4-2B | BC | Pf | Y | - | Y | Y | - |
| **SET110** | D19b | BC 4-2B | BC | Pf | Y | Y | - | Y | - |
| **SET111** | D20b | BC 4-2B | BC | Pf | Y | Y | Y | Y | - |
| **SET112** | D21b | BC 4-2B | BC | Pf | -Dig | Y | Y | Y | - |
| **SET113** | D17c | BC 4-2B | BC | Pf | Y | - | - | Y | Y |
| **SET114** | D18c | BC 4-2B | BC | Pf | Y | - | Y | Y | - |
| **SET115** | D19c | BC 4-2B | BC | Pf | Y | Y | - | Y | - |
| **SET116** | D20c | BC 4-2B | BC | Pf | Y | Y | Y | Y | Y |
| **SETBK3-S** | D23 | Extraction blank | | | -DD | - | - | Y | - |
| **SETBK4-S** | D24 | Extraction blank | | | Y | Y | Y | Y | - |
| **SETBK5-S** | D25 | Extraction blank | | | Y | Y | Y | Y | Y |

Core and previous ID as per core slice designation in Sadoway (2014). Pf = permafrost; Y = treatment was used on sample; -DD = 1M EDTA demineralization overnight followed by proteinase K digestion buffer; -Dig = Same as DD without EDTA phase. SET samples on the left half of the divide were not sonicated. For samples on the right half that were, 25 µL of extract was added to 25 µL of EBT (see Table S14 for sonication run parameters). For samples that were sonicated, a subset was purified/concentrated with QiaQuick PCR purification kit back to 25 µL.

**Table S14** Sonication run parameters.

| | |
|---|---|
| Target bp (Peak) | 50–150 |
| Peak Incident Power (W) | 175 |
| Duty Factor | 10% |
| Cycles per burst | 200 |
| Treatment Time (s) | 480 |
| Temp (C) (+/-2) | 7 |

Minimum input volume is 50 μL, so 25 μL extracts were diluted with 25 μL EBT to bring up to volume.

**Table S15** SET-C sample list.

| SET ID | Core | Previous ID | 4°C Spin | 4°C Timing (hours) |
|---|---|---|---|---|
| SET118 | LLII 12-84-3 | LL3 | - | - |
| SET119 | LLII 12-84-3 | LL3 | - | - |
| SET120 | LLII 12-84-3 | LL3 | - | - |
| SET121 | LLII 12-84-3 | LL3 | Y | 1 |
| SET122 | LLII 12-84-3 | LL3 | Y | 1 |
| SET123 | LLII 12-84-3 | LL3 | Y | 1 |
| SET124 | LLII 12-84-3 | LL3 | Y | 6 |
| SET125 | LLII 12-84-3 | LL3 | Y | 6 |
| SET126 | LLII 12-84-3 | LL3 | Y | 6 |
| SET127 | LLII 12-84-3 | LL3 | Y | 19 |
| SET128 | LLII 12-84-3 | LL3 | Y | 19 |
| SET129 | LLII 12-84-3 | LL3 | Y | 19 |
| SET130 | LLII 12-217-8 | LL1 | - | - |
| SET131 | LLII 12-217-8 | LL1 | - | - |
| SET132 | LLII 12-217-8 | LL1 | - | - |
| SET133 | LLII 12-217-8 | LL1 | Y | 1 |
| SET134 | LLII 12-217-8 | LL1 | Y | 1 |
| SET135 | LLII 12-217-8 | LL1 | Y | 1 |
| SET136 | LLII 12-217-8 | LL1 | Y | 6 |
| SET137 | LLII 12-217-8 | LL1 | Y | 6 |
| SET138 | LLII 12-217-8 | LL1 | Y | 6 |
| SET139 | LLII 12-217-8 | LL1 | Y | 19 |
| SET140 | LLII 12-217-8 | LL1 | Y | 19 |
| SET141 | LLII 12-217-8 | LL1 | Y | 19 |
| SET142 | BC 4-2B | BC | - | - |
| SET143 | BC 4-2B | BC | - | - |
| SET144 | BC 4-2B | BC | - | - |
| SET145 | BC 4-2B | BC | Y | 1 |
| SET146 | BC 4-2B | BC | Y | 1 |
| SET147 | BC 4-2B | BC | Y | 1 |
| SET148 | BC 4-2B | BC | Y | 6 |
| SET149 | BC 4-2B | BC | Y | 6 |
| SET150 | BC 4-2B | BC | Y | 6 |
| SET151 | BC 4-2B | BC | Y | 19 |
| SET152 | BC 4-2B | BC | Y | 19 |
| SET153 | BC 4-2B | BC | Y | 19 |
| SETBK9 | Extraction Blank | | Y | 1 |
| SETBK10 | Extraction Blank | | Y | 6 |
| SETBK11 | Extraction Blank | | Y | 19 |
| SETLBK12 | Library Blank | | | |

All samples were physically disrupted with PowerBeads.

**Table S16** SET-D sample list.

| Sample Information | | | | Extraction Variants | | | |
|---|---|---|---|---|---|---|---|
| SET ID | Core | Previous ID | Sample Type | Lysing Method | Detergent | Lysing Timing (hours) | 4°C Timing (hours) |
| SET160 | LLII 12-217-8 | LL1 | Pf | PowerBead+Digest | Sarkosyl | 9 | 24 |
| SET161 | LLII 12-217-8 | LL1 | Pf | PowerBead+Digest | Sarkosyl | 9 | 24 |
| SET162 | LLII 12-217-8 | LL1 | Pf | PowerBead+Digest | Sarkosyl | 9 | 24 |
| SET163 | LLII 12-217-8 | LL1 | Pf | PowerBead+Digest | Sarkosyl | 19 | 24 |
| SET164 | LLII 12-217-8 | LL1 | Pf | PowerBead+Digest | Sarkosyl | 19 | 24 |
| SET165 | LLII 12-217-8 | LL1 | Pf | PowerBead+Digest | Sarkosyl | 19 | 24 |
| SET166 | LLII 12-217-8 | LL1 | Pf | PowerBead+Digest | Sarkosyl | 9 | 48 |
| SET167 | LLII 12-217-8 | LL1 | Pf | PowerBead+Digest | Sarkosyl | 9 | 48 |
| SET168 | LLII 12-217-8 | LL1 | Pf | PowerBead+Digest | Sarkosyl | 9 | 48 |
| SET169 | LLII 12-217-8 | LL1 | Pf | PowerBead+Digest | Sarkosyl | 19 | 48 |
| SET170 | LLII 12-217-8 | LL1 | Pf | PowerBead+Digest | Sarkosyl | 19 | 48 |
| SET171 | LLII 12-217-8 | LL1 | Pf | PowerBead+Digest | Sarkosyl | 19 | 48 |
| SET172 | LLII 12-217-8 | LL1 | Pf | Digest | Sarkosyl | 9 | 24 |
| SET173 | LLII 12-217-8 | LL1 | Pf | Digest | Sarkosyl | 9 | 24 |
| SET174 | LLII 12-217-8 | LL1 | Pf | Digest | Sarkosyl | 9 | 24 |
| SET175 | LLII 12-217-8 | LL1 | Pf | Digest | Sarkosyl | 19 | 24 |
| SET176 | LLII 12-217-8 | LL1 | Pf | Digest | Sarkosyl | 19 | 24 |
| SET177 | LLII 12-217-8 | LL1 | Pf | Digest | Sarkosyl | 19 | 24 |
| SET178 | LLII 12-217-8 | LL1 | Pf | Digest | Sarkosyl | 9 | 48 |
| SET179 | LLII 12-217-8 | LL1 | Pf | Digest | Sarkosyl | 9 | 48 |
| SET180 | LLII 12-217-8 | LL1 | Pf | Digest | Sarkosyl | 9 | 48 |
| SET181 | LLII 12-217-8 | LL1 | Pf | Digest | Sarkosyl | 19 | 48 |
| SET182 | LLII 12-217-8 | LL1 | Pf | Digest | Sarkosyl | 19 | 48 |
| SET183 | LLII 12-217-8 | LL1 | Pf | Digest | Sarkosyl | 19 | 48 |
| SET184 | LLII 12-217-8 | LL1 | Pf | EDTA-Digest | Sarkosyl | 9 | 24 |
| SET185 | LLII 12-217-8 | LL1 | Pf | EDTA-Digest | Sarkosyl | 9 | 24 |
| SET186 | LLII 12-217-8 | LL1 | Pf | EDTA-Digest | Sarkosyl | 9 | 24 |
| SET187 | LLII 12-217-8 | LL1 | Pf | EDTA-Digest | Sarkosyl | 19 | 24 |
| SET188 | LLII 12-217-8 | LL1 | Pf | EDTA-Digest | Sarkosyl | 19 | 24 |
| SET189 | LLII 12-217-8 | LL1 | Pf | EDTA-Digest | Sarkosyl | 19 | 24 |
| SET190 | LLII 12-217-8 | LL1 | Pf | EDTA-Digest | Sarkosyl | 9 | 48 |
| SET191 | LLII 12-217-8 | LL1 | Pf | EDTA-Digest | Sarkosyl | 9 | 48 |
| SET192 | LLII 12-217-8 | LL1 | Pf | EDTA-Digest | Sarkosyl | 9 | 48 |
| SET193 | LLII 12-217-8 | LL1 | Pf | EDTA-Digest | Sarkosyl | 19 | 48 |
| SET194 | LLII 12-217-8 | LL1 | Pf | EDTA-Digest | Sarkosyl | 19 | 48 |
| SET195 | LLII 12-217-8 | LL1 | Pf | EDTA-Digest | Sarkosyl | 19 | 48 |
| SET196 | BC 4-2B | BC | Pf | PowerBead+Digest | Sarkosyl | 9 | 24 |
| SET197 | BC 4-2B | BC | Pf | PowerBead+Digest | Sarkosyl | 9 | 24 |
| SET198 | BC 4-2B | BC | Pf | PowerBead+Digest | Sarkosyl | 9 | 24 |
| SET199 | BC 4-2B | BC | Pf | PowerBead+Digest | Sarkosyl | 19 | 24 |
| SET200 | BC 4-2B | BC | Pf | PowerBead+Digest | Sarkosyl | 19 | 24 |
| SET201 | BC 4-2B | BC | Pf | PowerBead+Digest | Sarkosyl | 19 | 24 |
| SET202 | BC 4-2B | BC | Pf | PowerBead+Digest | Sarkosyl | 9 | 48 |
| SET203 | BC 4-2B | BC | Pf | PowerBead+Digest | Sarkosyl | 9 | 48 |
| SET204 | BC 4-2B | BC | Pf | PowerBead+Digest | Sarkosyl | 9 | 48 |
| SET205 | BC 4-2B | BC | Pf | PowerBead+Digest | Sarkosyl | 19 | 48 |
| SET206 | BC 4-2B | BC | Pf | PowerBead+Digest | Sarkosyl | 19 | 48 |
| SET207 | BC 4-2B | BC | Pf | PowerBead+Digest | Sarkosyl | 19 | 48 |
| SET208 | BC 4-2B | BC | Pf | Digest | Sarkosyl | 9 | 24 |
| SET209 | BC 4-2B | BC | Pf | Digest | Sarkosyl | 9 | 24 |
| SET210 | BC 4-2B | BC | Pf | Digest | Sarkosyl | 9 | 24 |
| SET211 | BC 4-2B | BC | Pf | Digest | Sarkosyl | 19 | 24 |
| SET212 | BC 4-2B | BC | Pf | Digest | Sarkosyl | 19 | 24 |
| SET213 | BC 4-2B | BC | Pf | Digest | Sarkosyl | 19 | 24 |
| SET214 | BC 4-2B | BC | Pf | Digest | Sarkosyl | 9 | 48 |
| SET215 | BC 4-2B | BC | Pf | Digest | Sarkosyl | 9 | 48 |
| SET216 | BC 4-2B | BC | Pf | Digest | Sarkosyl | 9 | 48 |
| SET217 | BC 4-2B | BC | Pf | Digest | Sarkosyl | 19 | 48 |
| SET218 | BC 4-2B | BC | Pf | Digest | Sarkosyl | 19 | 48 |
| SET219 | BC 4-2B | BC | Pf | Digest | Sarkosyl | 19 | 48 |
| SET220 | BC 4-2B | BC | Pf | EDTA-Digest | Sarkosyl | 9 | 24 |
| SET221 | BC 4-2B | BC | Pf | EDTA-Digest | Sarkosyl | 9 | 24 |
| SET222 | BC 4-2B | BC | Pf | EDTA-Digest | Sarkosyl | 9 | 24 |
| SET223 | BC 4-2B | BC | Pf | EDTA-Digest | Sarkosyl | 19 | 24 |
| SET224 | BC 4-2B | BC | Pf | EDTA-Digest | Sarkosyl | 19 | 24 |
| SET225 | BC 4-2B | BC | Pf | EDTA-Digest | Sarkosyl | 19 | 24 |
| SET226 | BC 4-2B | BC | Pf | EDTA-Digest | Sarkosyl | 9 | 48 |
| SET227 | BC 4-2B | BC | Pf | EDTA-Digest | Sarkosyl | 9 | 48 |
| SET228 | BC 4-2B | BC | Pf | EDTA-Digest | Sarkosyl | 9 | 48 |
| SET229 | BC 4-2B | BC | Pf | EDTA-Digest | Sarkosyl | 19 | 48 |
| SET230 | BC 4-2B | BC | Pf | EDTA-Digest | Sarkosyl | 19 | 48 |
| SET231 | BC 4-2B | BC | Pf | EDTA-Digest | Sarkosyl | 19 | 48 |
| BK15 | | | E-B | PowerBead+Digest | Sarkosyl | 9 | |
| BK16 | | | E-B | EDTA-Digest | Sarkosyl | 9 | |
| BK17 | | | E-B | PowerBead+Digest | Sarkosyl | 19 | |
| BK18 | | | E-B | EDTA-Digest | Sarkosyl | 19 | |
| SET244 | BC 4-2B | BC | Pf | PowerBead+Digest | SDS | 19 | 24 |
| SET245 | BC 4-2B | BC | Pf | PowerBead+Digest | SDS | 19 | 24 |
| SET246 | BC 4-2B | BC | Pf | PowerBead+Digest | SDS | 19 | 24 |
| SET247 | BC 4-2B | BC | Pf | PowerBead+Digest | SDS | 19 | 48 |
| SET248 | BC 4-2B | BC | Pf | PowerBead+Digest | SDS | 19 | 48 |
| SET249 | BC 4-2B | BC | Pf | PowerBead+Digest | SDS | 19 | 48 |
| SET250 | BC 4-2B | BC | Pf | EDTA-Digest | SDS | 19 | 24 |
| SET251 | BC 4-2B | BC | Pf | EDTA-Digest | SDS | 19 | 24 |
| SET252 | BC 4-2B | BC | Pf | EDTA-Digest | SDS | 19 | 24 |
| SET253 | BC 4-2B | BC | Pf | EDTA-Digest | SDS | 19 | 48 |
| SET254 | BC 4-2B | BC | Pf | EDTA-Digest | SDS | 19 | 48 |
| SET255 | BC 4-2B | BC | Pf | EDTA-Digest | SDS | 19 | 48 |
| BK19 | | | E-B | PowerBead+Digest | SDS | 19 | 24 |
| BK20 | | | E-B | EDTA-Digest | SDS | 19 | 48 |

Pf = permafrost; E-B = extraction blank. SET-$D_1$ in grey (left and upper right sections), SET-$D_2$ in blue (bottom right section).

**Table S17** Map-filtered reads *MEGAN* LCA-assignment summary, SET-E.

| SedaDNA Extraction | Targeting Strategy | Total reads | Map-filtered summation | | | | Map-filtered select major clade summations (reads mapped-to-baits, string de-duplicated, ≥ 24bp) | | | | | | | | | | | |
|---|---|---|---|---|---|---|---|---|---|---|---|---|---|---|---|---|---|---|
| | | | Mapped-to-baits* | | *BLASTn* aligned & *MEGAN* assigned | | Bacteria and Archaea | | Fungi | | Metazoa | | Virdiplantae | | No *BLASTn* hits | | Not LCA-assigned | |
| PowerSoil | Enrichment | 14,292,697 | 41,592 | **0.3%** | 36,693 | ***0.3%*** | 15 | **0.0%** | 0 | **0.0%** | 143 | **0.3%** | 36,507 | **87.8%** | 2,149 | **5.2%** | 2,750 | **6.6%** |
| Modified Dabney | Enrichment | 15,516,557 | 961,734 | **6.4%** | 835,364 | ***5.6%*** | 282 | **0.0%** | 56 | **0.0%** | 8,152 | **0.8%** | 826,203 | **85.9%** | 68,228 | **7.1%** | 58,132 | **6.0%** |
| PowerSoil | Shotgun | 6,071,164 | 161 | **0.0%** | 50 | ***0.0%*** | 0 | **0.0%** | 0 | **0.0%** | 0 | **0.0%** | 50 | **31.1%** | 103 | **64.0%** | 8 | **5.0%** |
| Modified Dabney | Shotgun | 14,911,050 | 2,216 | **0.0%** | 470 | ***0.0%*** | 4 | **0.2%** | 0 | **0.0%** | 8 | **0.4%** | 449 | **20.3%** | 1,642 | **74.1%** | 104 | **4.7%** |
| D'Costa et al. | Metabarcoding | 1,097,644 | 3176 | **0.3%** | 3078 | ***0.3%*** | 0 | **0.0%** | 0 | **0.0%** | 463 | **14.6%** | 2,608 | **82.1%** | 57 | **1.8%** | 41 | **1.3%** |

Map-filtered summation: percent of total reads.
Map-filtered major clade summations: percent of mapped-to-baits*.

**Table S18** Non-map-filtered reads *MEGAN* LCA-assignment summary, SET-E.

| SedaDNA Extraction | Targeting Strategy | Total reads | Non-map-filtered select major clade summations (string de-duplicated and ≥ 24bp) | | | | | | | | | | | |
|---|---|---|---|---|---|---|---|---|---|---|---|---|---|---|
| | | | Bacteria & Archea | | Fungi | | Metazoa | | Virdiplantae | | No *BLASTn* hits | | Not LCA-assigned | |
| PowerSoil | Enrichment | 14,292,697 | 243,941 | **1.7%** | 3,234 | **0.0%** | 40,345 | **0.3%** | 462,592 | **3.2%** | 9,178,308 | **64.2%** | 877,851 | **6.1%** |
| Modified Dabney | Enrichment | 15,516,557 | 127,382 | **0.8%** | 11,354 | **0.1%** | 93,089 | **0.6%** | 1,685,455 | **10.9%** | 9,368,011 | **60.4%** | 526,480 | **3.4%** |
| PowerSoil | Shotgun | 6,071,164 | 113,216 | **1.9%** | 728 | **0.0%** | 24,177 | **0.4%** | 44,852 | **0.7%** | 4,997,565 | **82.3%** | 488,337 | **8.0%** |
| Modified Dabney | Shotgun | 14,911,050 | 150,986 | **1.0%** | 3,141 | **0.0%** | 87,745 | **0.6%** | 233,197 | **1.6%** | 11,863,278 | **79.6%** | 499,037 | **3.3%** |
| D'Costa et al. | Metabarcoding | 1,097,644 | 43 | **0.0%** | 0 | **0.0%** | 20,245 | **1.8%** | 76,953 | **7.0%** | 14,097 | **1.3%** | 17,690 | **1.6%** |

Non-map-filtered major clade summations: percent of total reads.

### Figures

**Figure S1** Bioanalyzer, high sensitive DNA assay, SET-A.

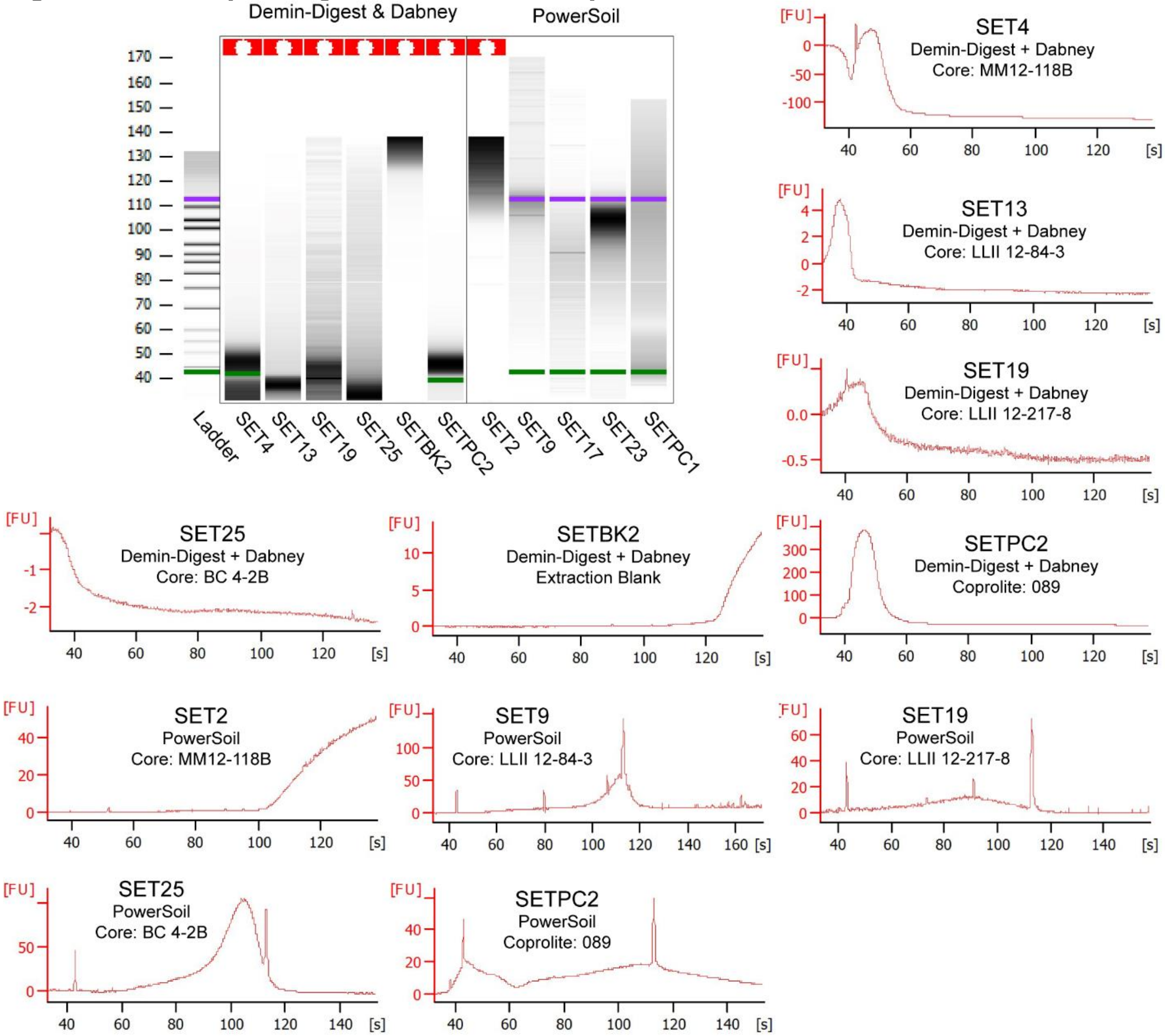

Run on an Agilent 2100 Bioanalyzer. Note that lanes 2-8 failed (too darkly coloured to detect baseline florescence), likely due to a high inhibition (humic) load.
PowerSoil: DNeasy PowerSoil extraction kit.
Demin-Digest + Dabney: demineralization (0.5 M EDTA) and digestion (proteinase K buffer, see Table 2) (each overnight separately) followed by purification with a high-volume binding buffer and silica column following Dabney et al. (2013).
089: *N. shastensis* palaeofeces from (Poinar et al., 1998).
Core ID as per Sadoway (2014).

**Figure S2** Bioanalyzer, high sensitive DNA assay, SET-A with a 1/10 dilution.

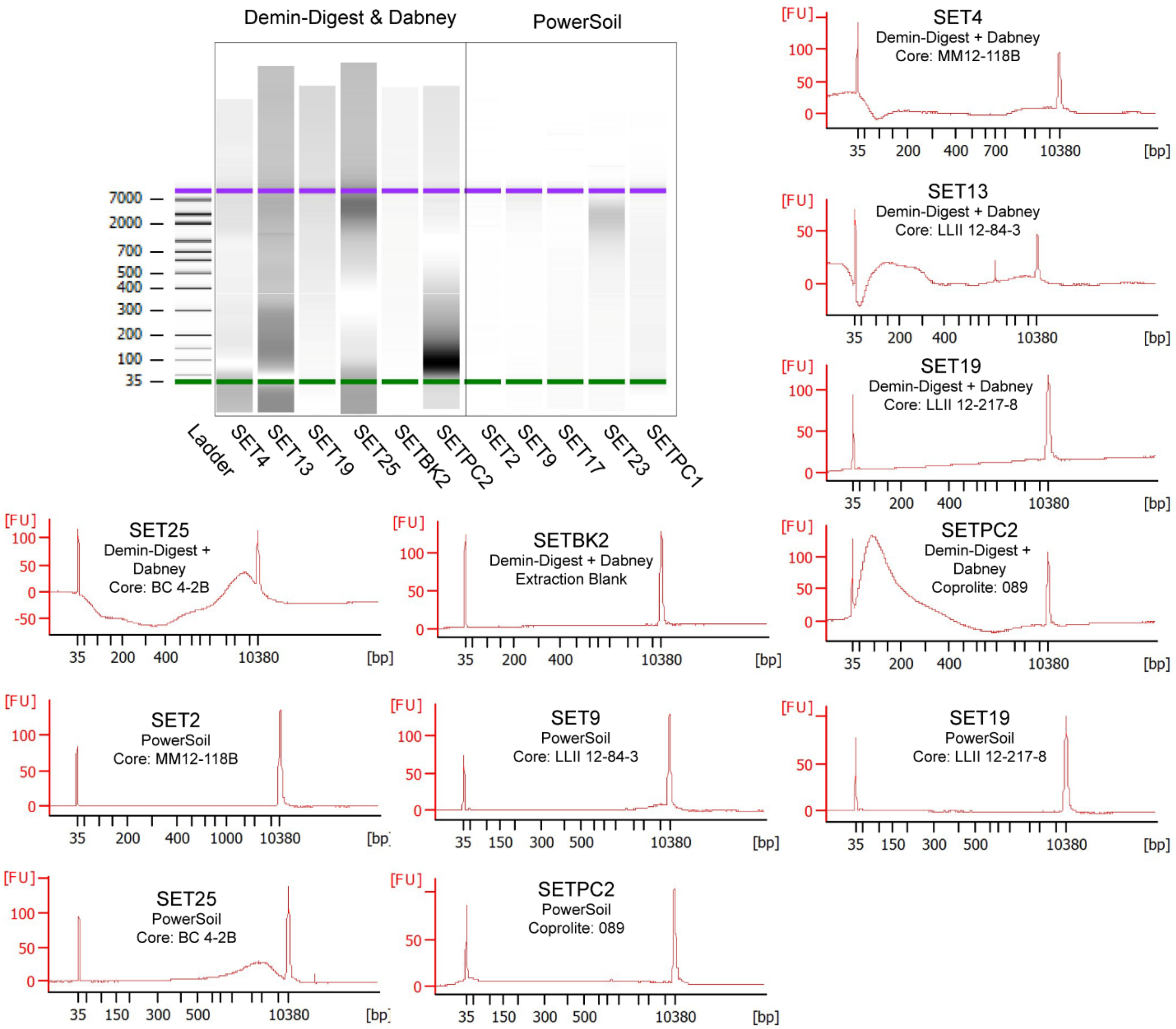

Run on an Agilent 2100 Bioanalyzer.
PowerSoil: DNeasy PowerSoil extraction kit.
Demin-Digest + Dabney: demineralization (0.5 M EDTA) and digestion (proteinase K buffer, see Table 2) (each overnight separately) followed by purification with a high-volume binding buffer and silica column following Dabney et al. (2013).
089: *N. shastensis* palaeofeces from (Poinar et al., 1998).
Core ID as per Sadoway (2014).

**Figure S3** SET-A, indexing RFU barchart for qPCR cycles 1 and 12.

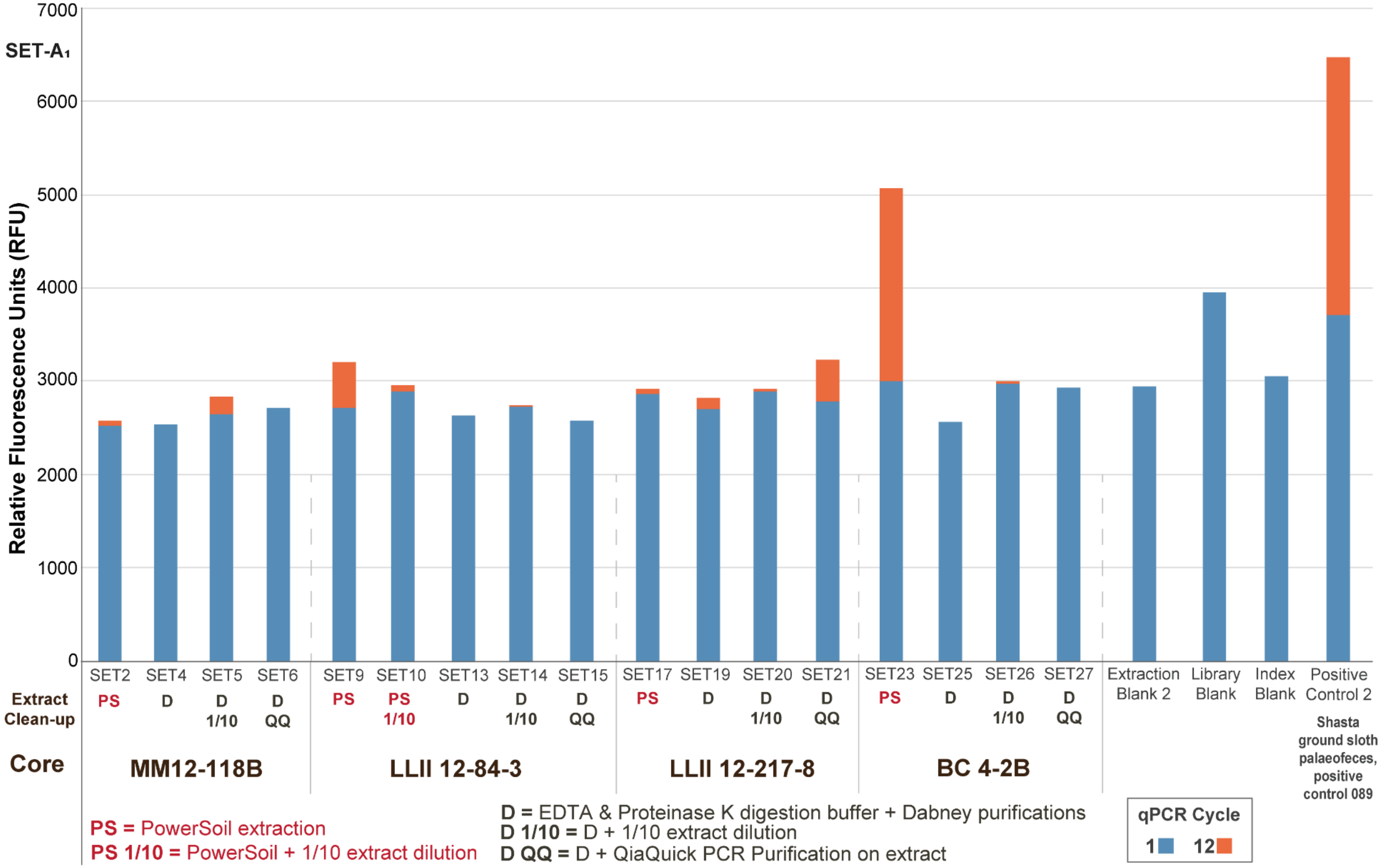

Extract clean-up indicates how the purified elutes were ‘cleaned’ prior to double stranded library preparation.

**Figure S4** SET-A, Inhibition indices of inhibitor clean-up strategies.

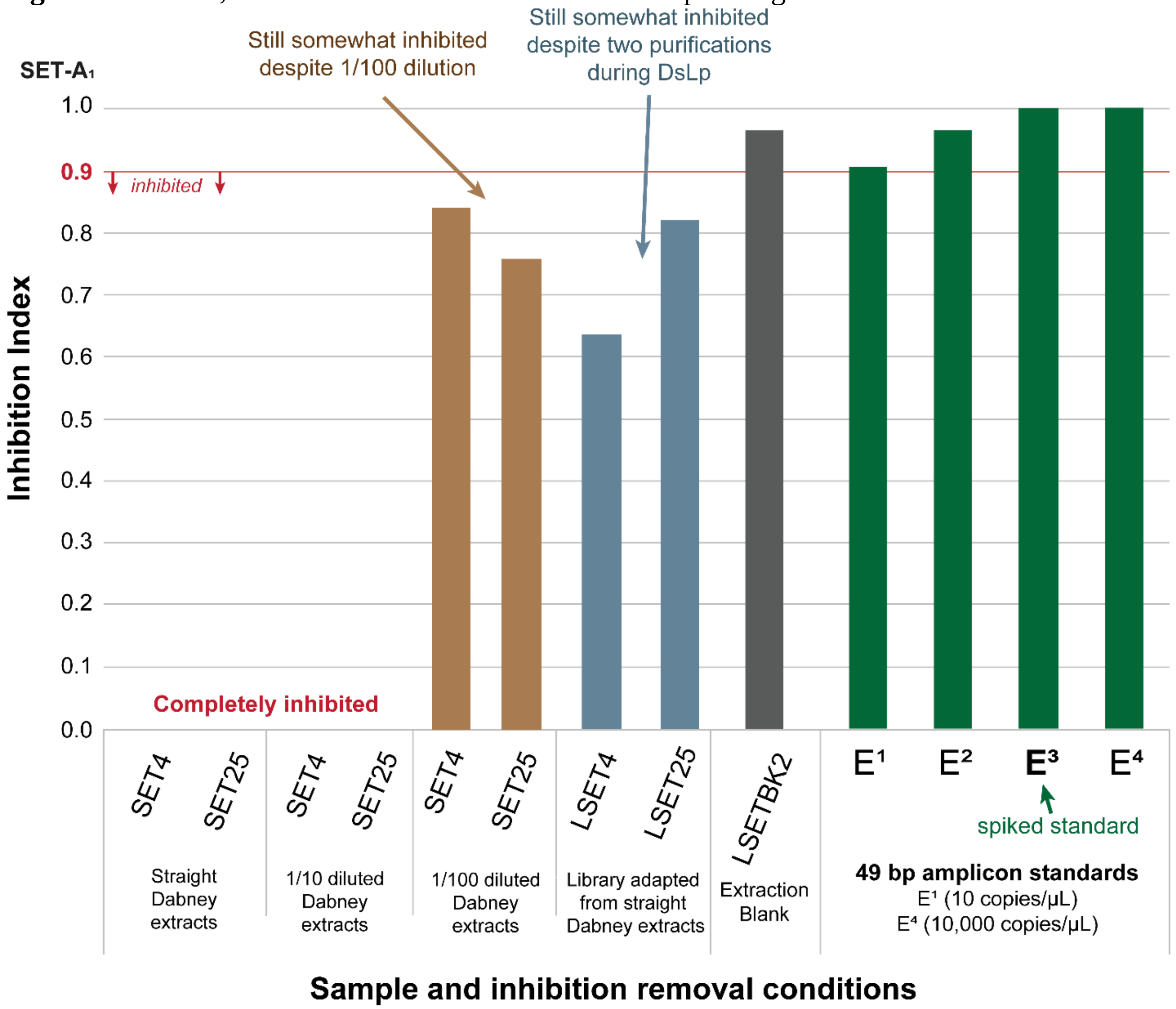

First three column sets are extracts that were spiked for the assay. SET samples with a preceding ‘L’ denote libraries that were assayed for inhibition.

**Figure S5** SET-A, Indexing qPCR reaction of positive control spiked samples prior to library preparation.

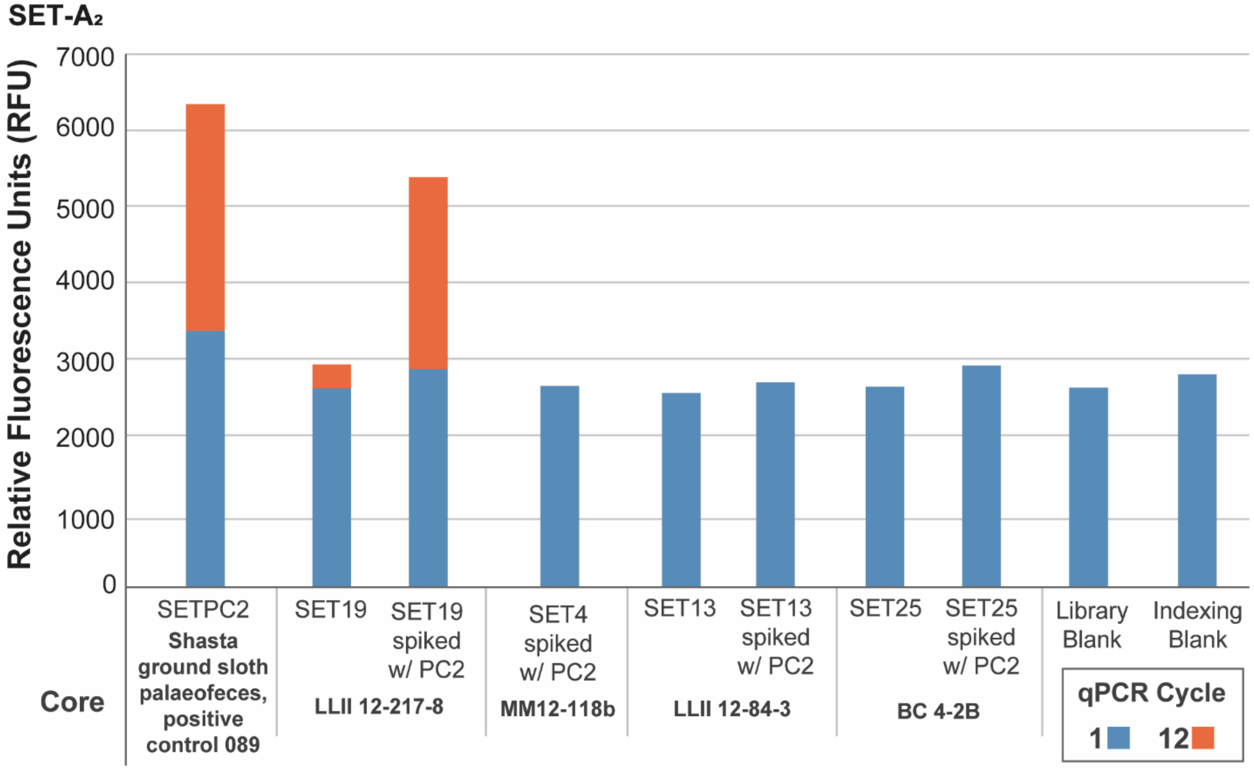

The SET4 extract was exhausted; only SET4 with a positive control spike was tested in this experiment.

**Figure S6** SET-B, comparing treatments for enzymatic inhibitor removal by their DNA retention.

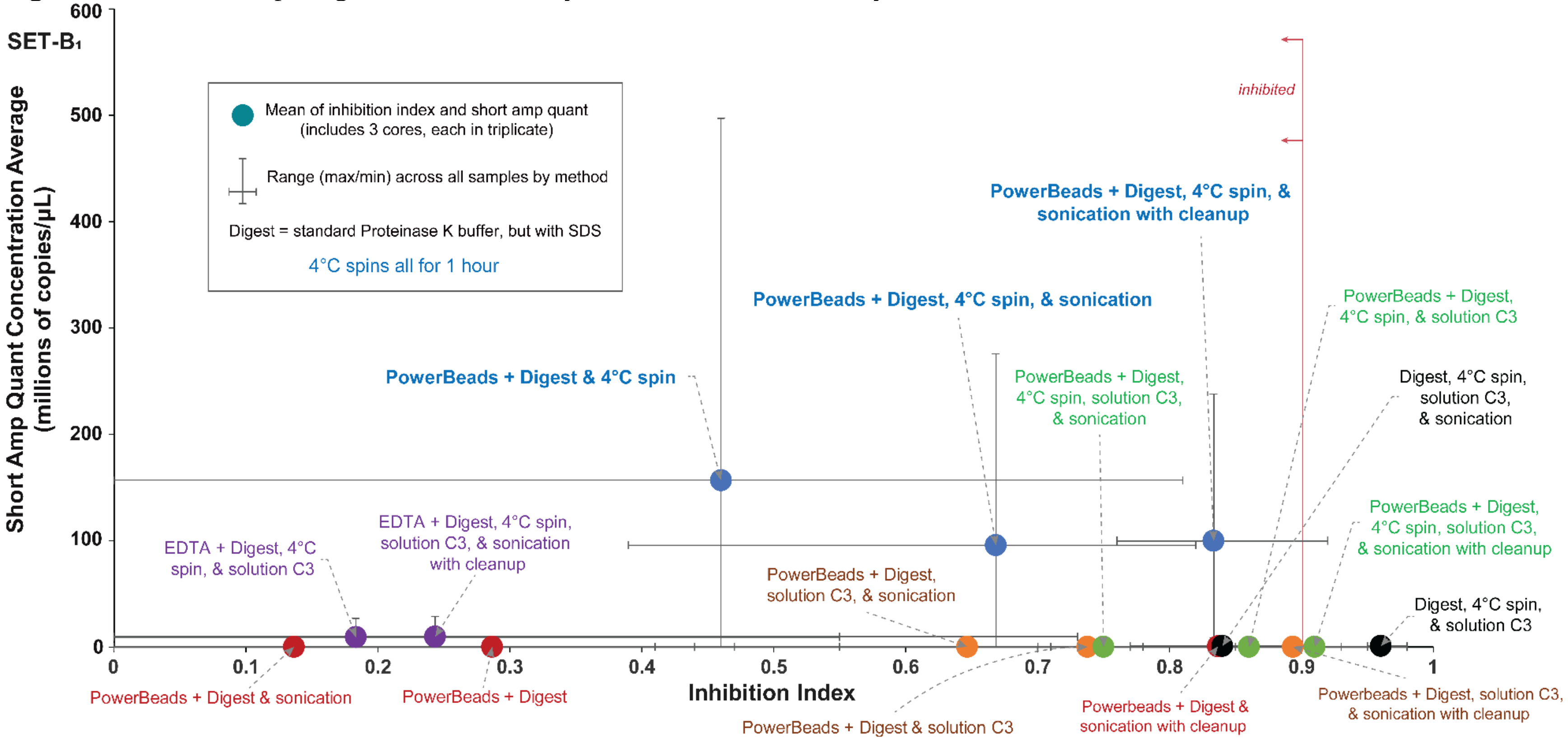

Details for the short amp DNA quantification can be found in Table S8. See Table S7 and Figure E14 (main text) for details on the inhibition index. See Table S13 for SET-B sample list. Short amp qPCR standard curves for plates 1 and 2 respectively: E = 103.8% and 92.4%, $R^2$= 0.999 and 0.995, slope = −3.234 and −3.519.

**Figure S7** Conceptual balance of overcoming sedaDNA inhibition co-elution

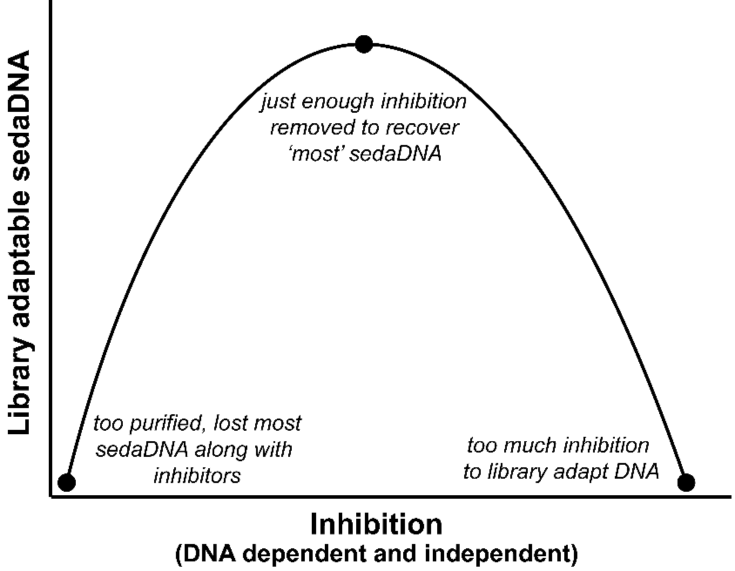

**Figure S8** SET-B, qPCR indexing reaction to confirm correlation with short amp quantification.

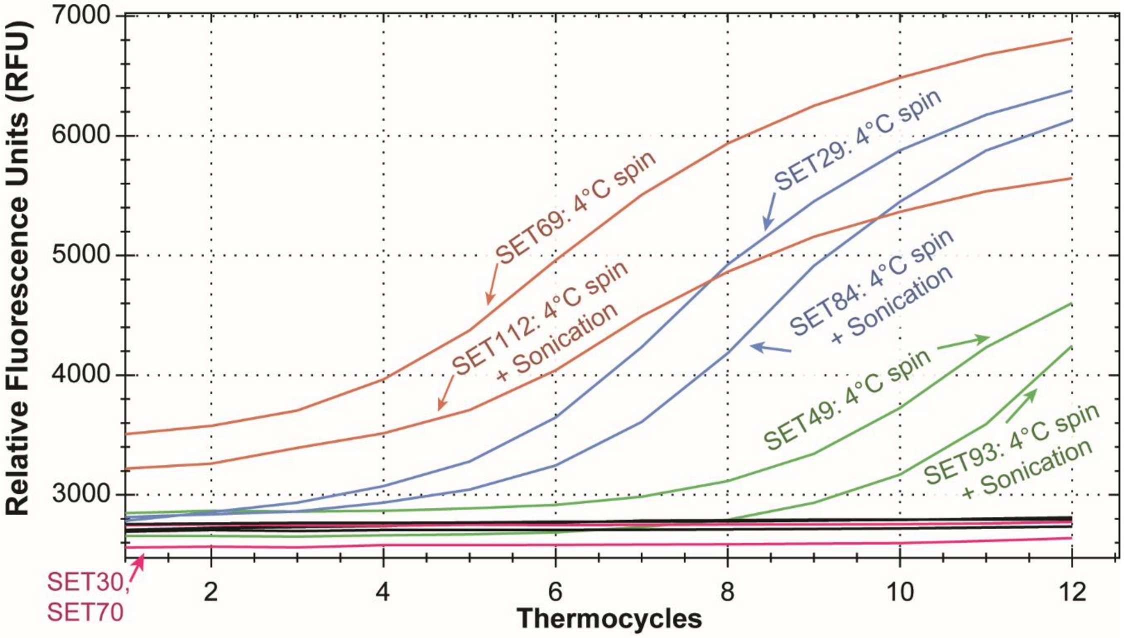

SET-B₁

| SET ID | Core | PowerBeads | PowerSoil Solution | 1 hour 4°C Spin | Sonication | Post-Sonication Purification |
|---|---|---|---|---|---|---|
| SET69 | BC 4-2B | Y | - | Y | - | - |
| SET112 | BC 4-2B | Y | - | Y | Y | - |
| SET29 | LLII 12-84-3 | Y | - | Y | - | - |
| SET84 | LLII 12-84-3 | Y | - | Y | Y | - |
| SET49 | LLII 12-217-8 | Y | - | Y | - | - |
| SET93 | LLII 12-217-8 | Y | - | Y | Y | - |
| SET30 | LLII 12-84-3 | Y | Y | - | - | - |
| SET70 | BC 4-2B | Y | Y | - | - | - |

See Table S6 for indexing qPCR specifications. See Table S13 for SET-B sample list.

**Figure S9** Variable duration 4°C centrifuge on the carryover of enzymatic inhibitors and library adapted DNA, SET-C.

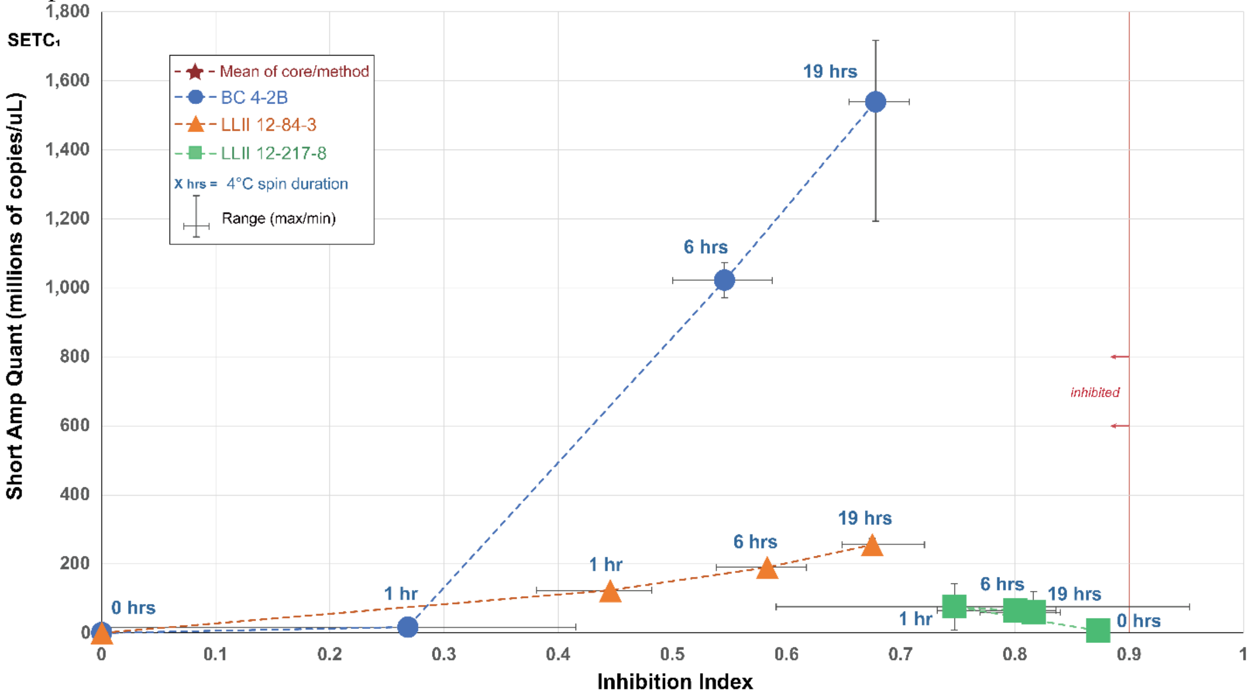

Short amp qPCR standard curve: E = 100.7%, $R^2$= 0.998, slope = −3.306. See Table S15 for sample list.

**Figure S10** Total double stranded DNA with variable extract input and blunt-end repair (BER) enzymatic concentrations, SET-C.

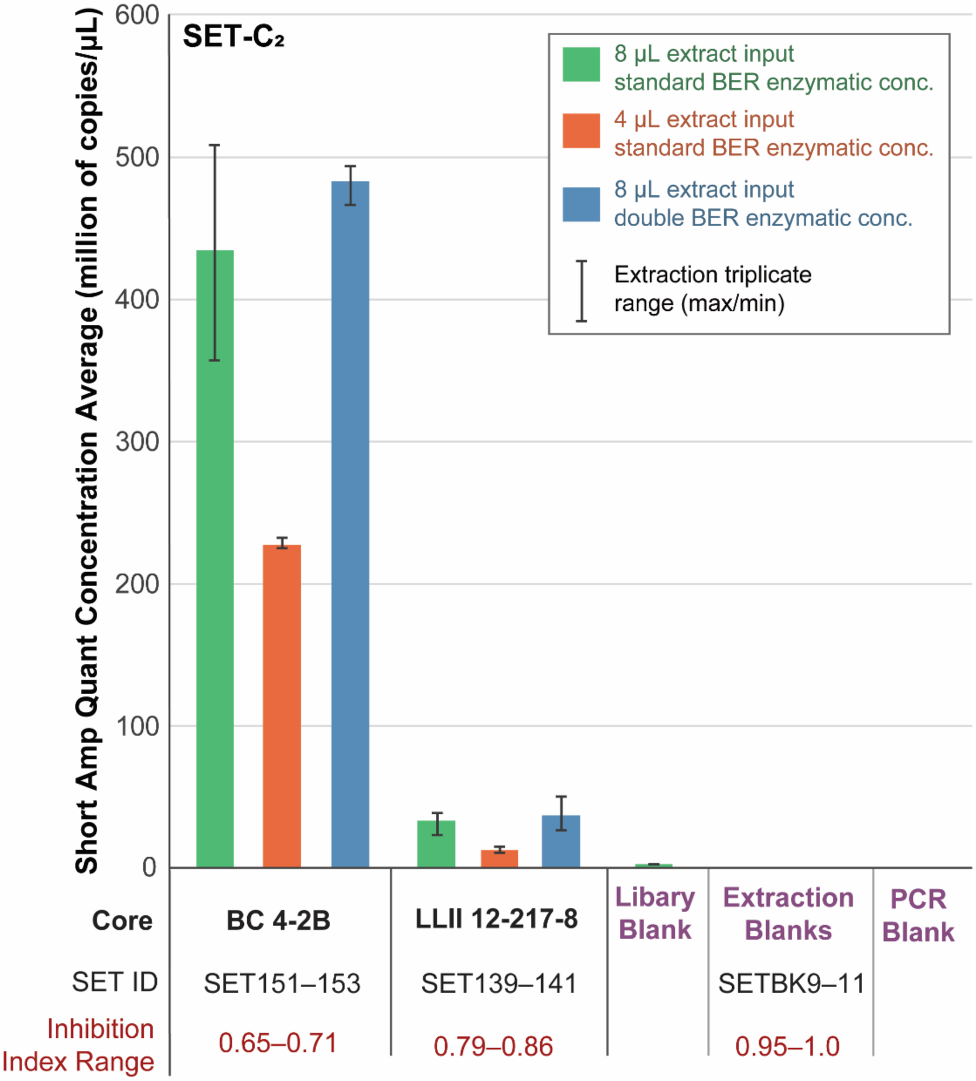

BER enzymatic conc. = blunt-end repair enzymatic concentrations. PCR triplicate used to determine DNA concentration average per SET sample. Extraction triplicate used to determine mean and range of DNA concentration average by method and core. Short amp qPCR standard curve: E = 100.3%, $R^2$= 0.992, slope = −3.314. An inhibition index < 0.9 is inhibited.

**Figure S11** SET-D, core BC 4-2B, variation in co-eluate inhibitor retention by lysing method and inhibitor removal procedure.

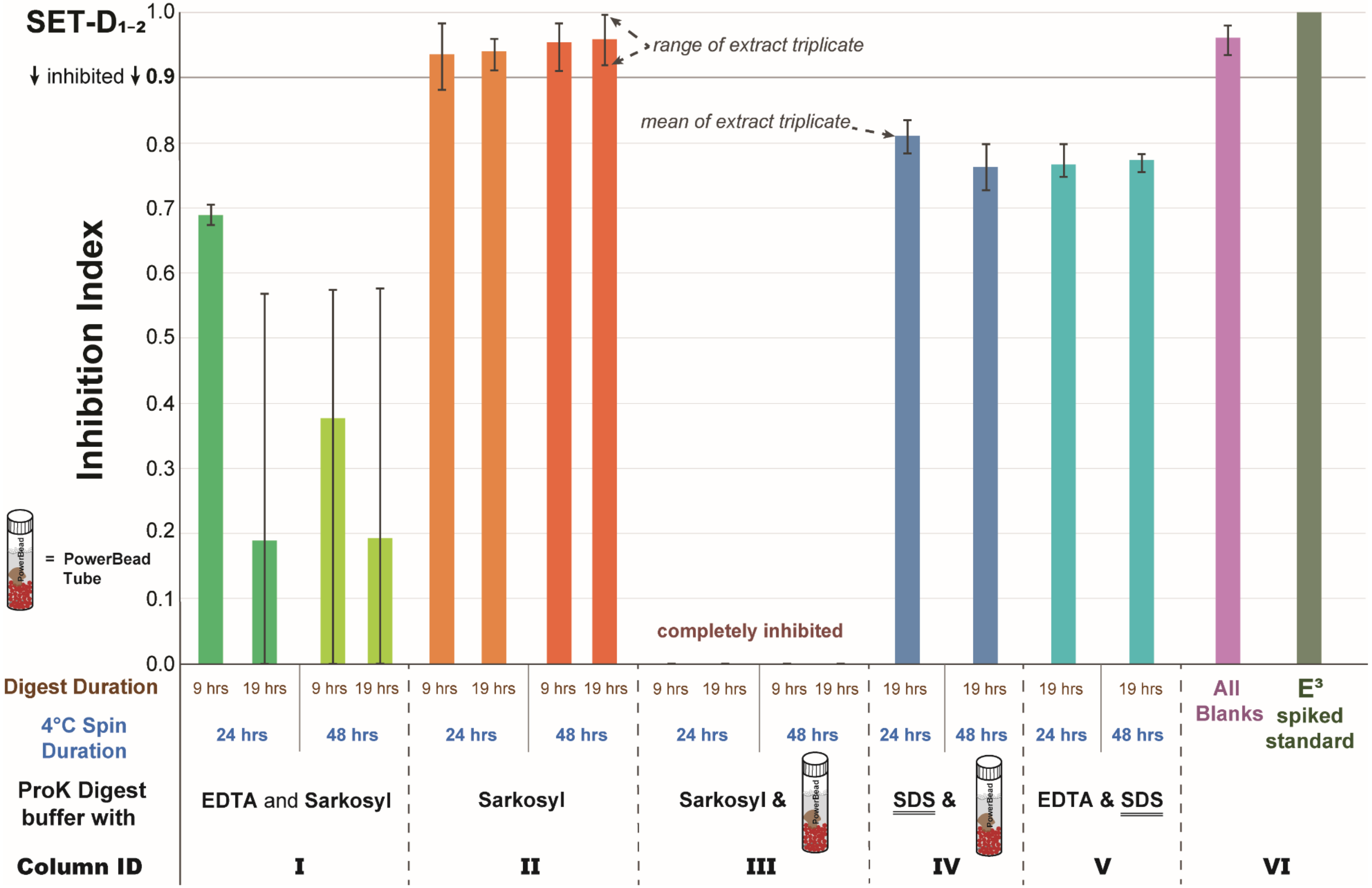

PCR triplicate used to determine inhibition index per SET sample. Extraction triplicate used to determine mean and range of inhibition indices by method and core. See Table S16 for sample list.

**Figure S12** SET-D Variable lysis disruption and cold spin duration for core BC 4-2B.

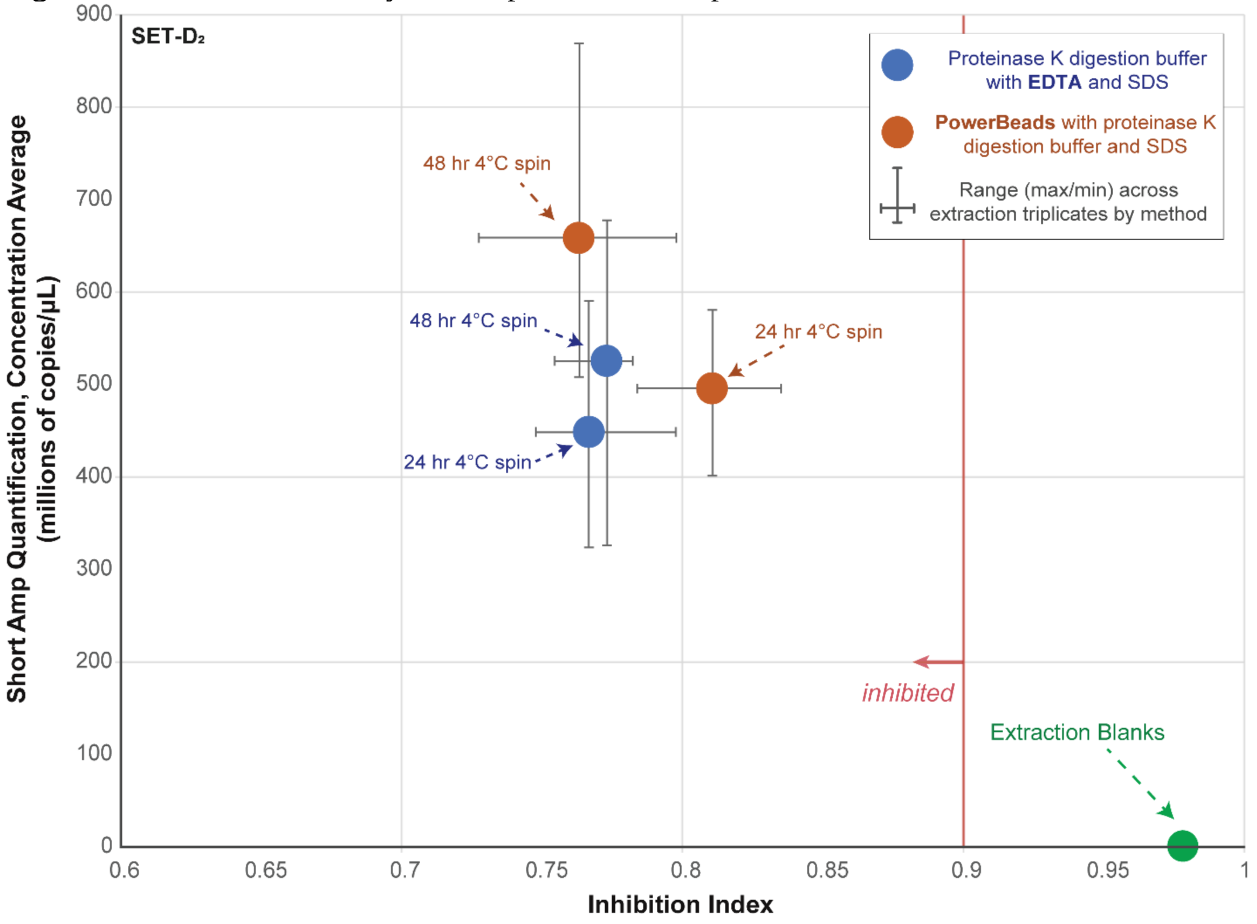

Details for the short amp DNA quantification can be found in Table S8. See Table S7 and Figure 1 (main text) for details on the inhibition index. See Table S16 for SET-D sample list. PCR triplicates used to determine average copies per µL per SET sample. Extraction triplicates used to determine mean and range of inhibition indices and short amp quantifications by method. Short amp qPCR standard curve: E = 100.5%, $R^2$= 0.998, slope = −3.310.

**Figure S13** Variable extract amplification of core LLII 12-217-8, *trn*L 'endogenous' qPCR

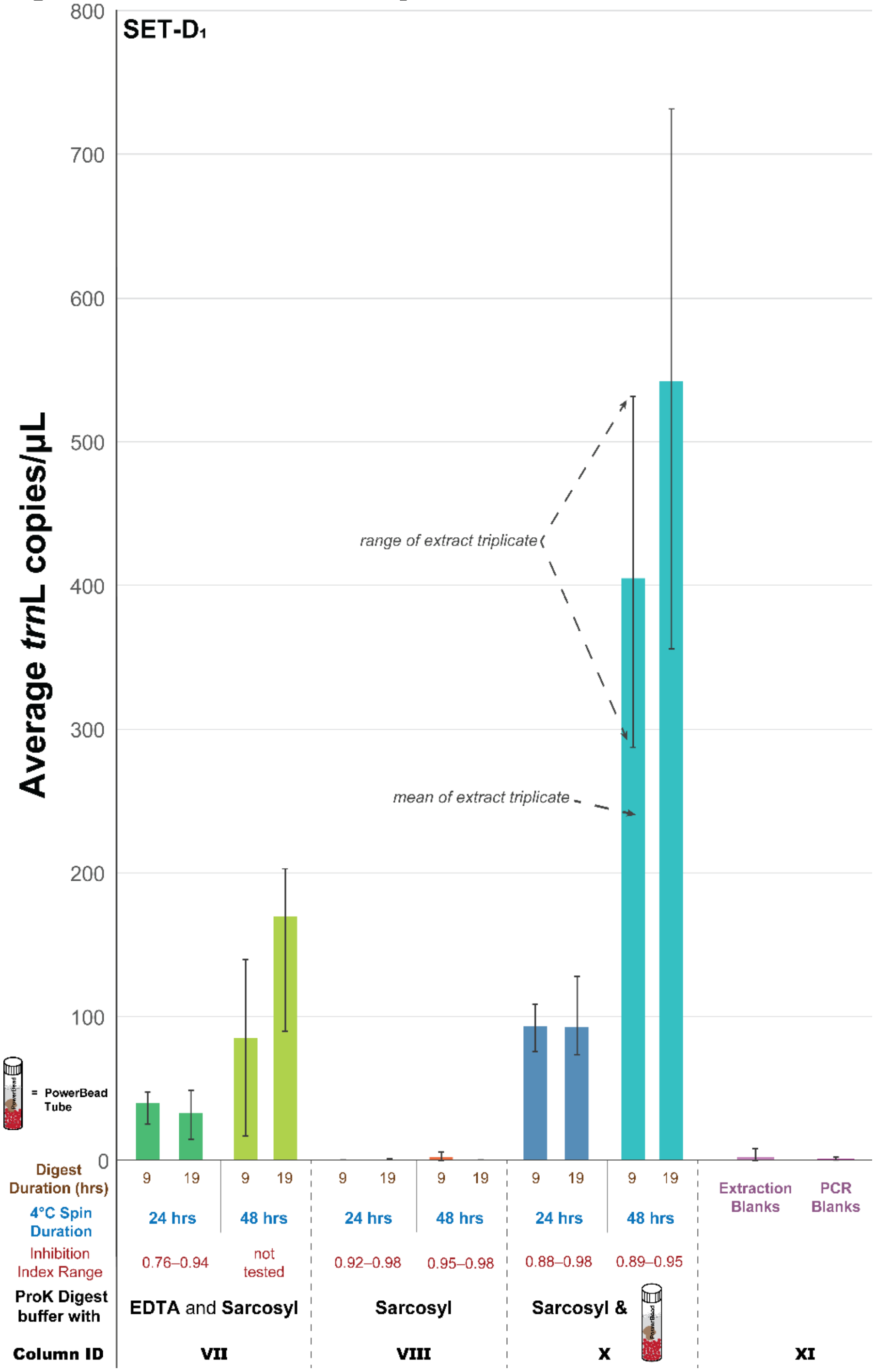

This core sample had low inhibition in all experiments, regardless of the inhibition removal technique used. It was the only "uninhibited" sample in the SET-A positive control spike test (Figure S5). The wide variation in DNA concentrations is due to non-standard PCR amplification curves. As such, *these values are unreliable indicators of actual DNA concentration in the extracts*. This data was included despite being unreliable because it shows that column VIII has no amplifiable DNA and no inhibition, ruling it out as a potential lysis option (as related to column II in Figure S11). PCR triplicates used to determine average *trn*L copies per µL per SET sample. Extraction triplicate used to determine mean and range of inhibition indices by method. See Table S16 for sample list. QPCR standard curves for plates 1 and 2 respectively: E = 97.5% and 101.1%, $R^2$= 0.983 and 0.985, slope = −3.373 and −3.296. Column IDs correlated with Figure S11 and Figure S14.

**Figure S14** Variable AmpliTaq Gold concentrations on *trn*L 'endogenous' sedaDNA qPCR amplifications

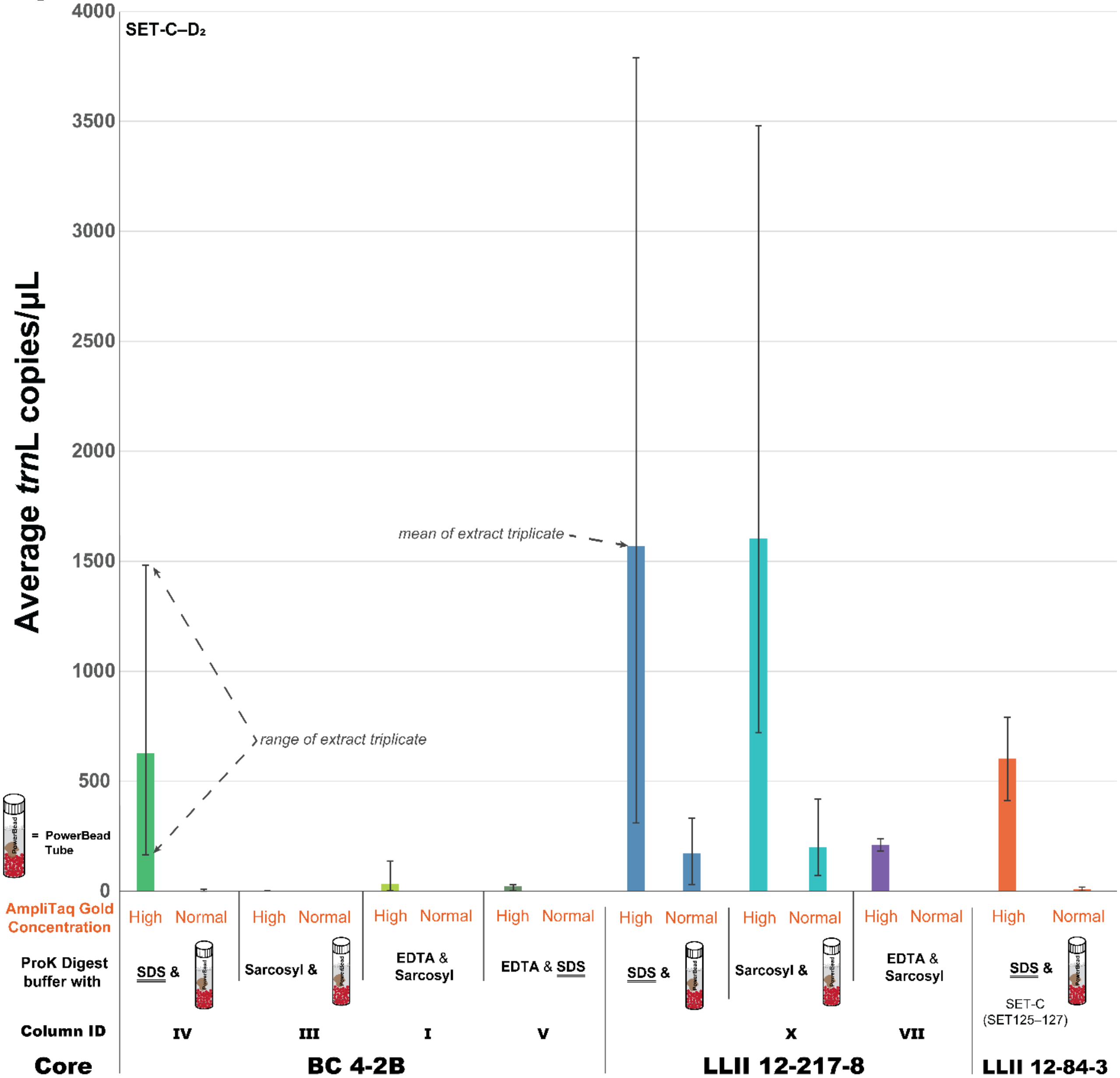

The wide variation in DNA concentration is due to non-standard PCR amplification curves. As such, *these values are unreliable indicators of actual DNA concentration in the extracts.* This data was included despite being unreliable because it shows that there is some sort of inhibition affecting these extracts, even for the core with low levels of inhibition (LLII 12-217-8, e.g. Figure S5). PCR triplicates used to determine average *trn*L copies per µL per SET sample. Extraction triplicate used to determine mean and range of inhibition indices by method. QPCR standard curve: E = 103.8%, $R^2$= 0.983 and 0.992, slope = −3.234. Column IDs correlated with Figure S11 and Figure S13.

542 **Figure S15** Metagenomic comparison of Upper Goldbottom permafrost core MM12-118b, reads
543 mapped to baits, normalized counts.

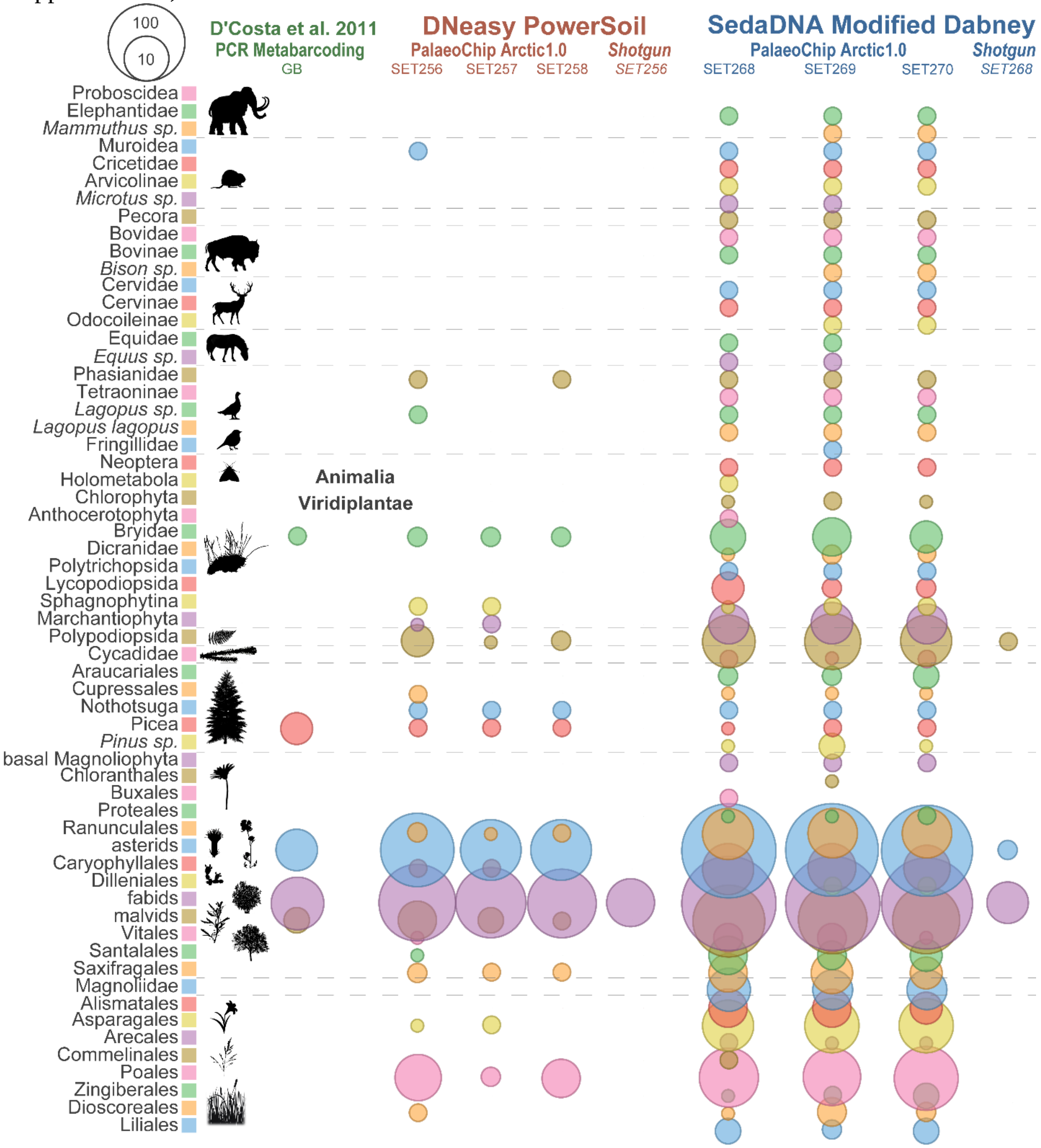

544
545 Core slice dated to 9,685 cal-BP (Mahony, 2015; Sadoway, 2014). Values indicate total reads assigned to
546 that taxon node for Animalia, and a clade summation of reads for Viridiplantae.
547

**Figure S16** Metagenomic comparison of Lucky Lady II permafrost core LLII-12-84-3, reads mapped to baits, normalized counts.

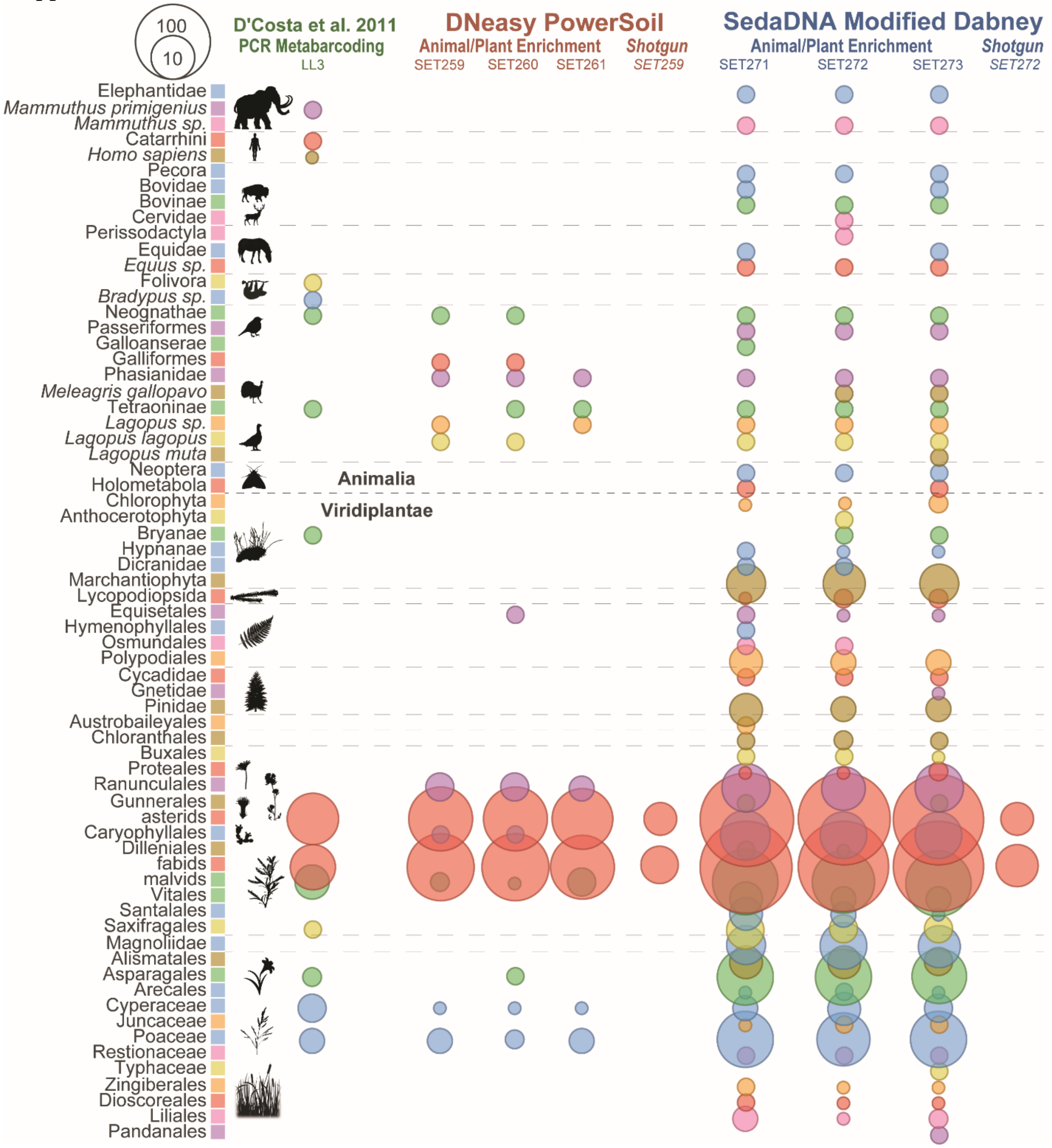

Core slice dated to 13,205 cal-BP (Sadoway, 2014). Values indicate total reads assigned to that taxon node for Animalia, and a clade summation of reads for Viridiplantae.

**Figure S17** Metagenomic comparison of Lucky Lady II permafrost core LLII-12-217-8, reads mapped to baits, normalized counts.

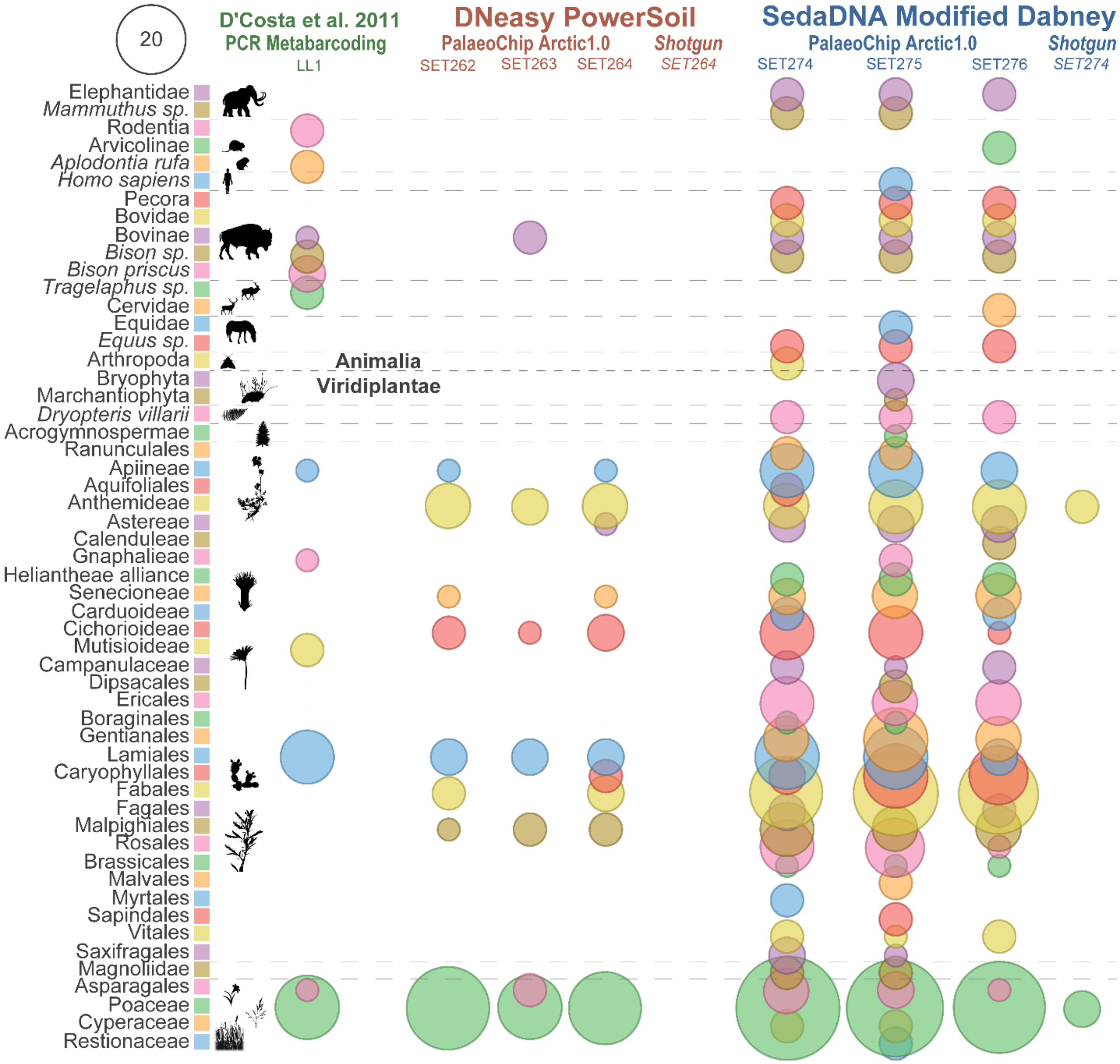

Core slice dated to 15,865 cal-BP (Sadoway, 2014). Values indicate total reads assigned to that taxon node for Animalia, and a clade summation of reads for Viridiplantae.

**Figure S18** Metagenomic comparison of Bear Creek permafrost core BC 4-2B, reads mapped to baits, normalized counts.

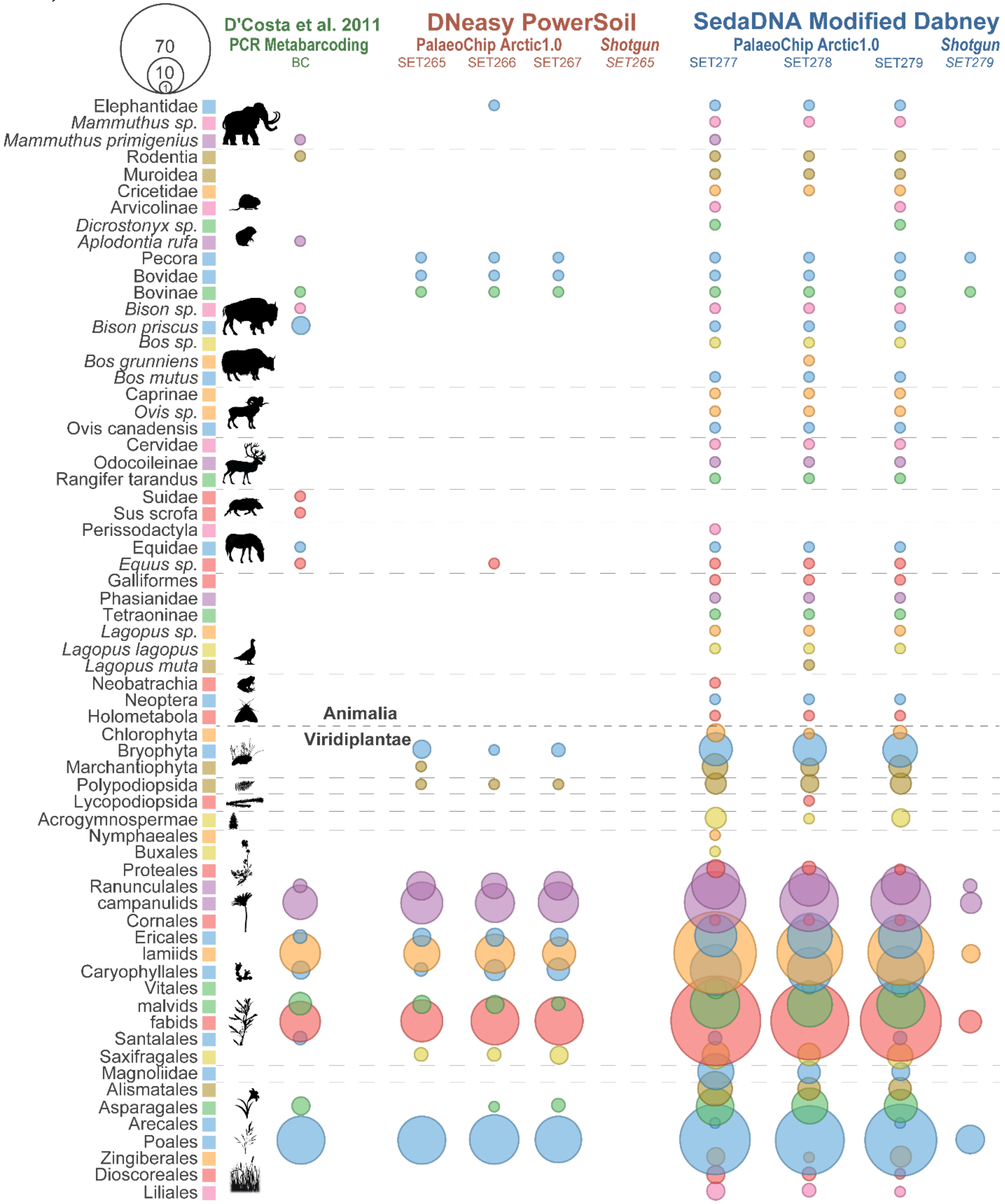

Core slice dated to ~30,000 cal-BP (D'Costa et al., 2011; Mahony, 2015; Sadoway, 2014). Values indicate total reads assigned to that taxon node for Animalia, and a clade summation of reads for Viridiplantae.

**Figure S19** Metagenomic comparison of Upper Goldbottom permafrost core MM12-118b, all reads (not map-filtered), absolute counts (non-normalized).

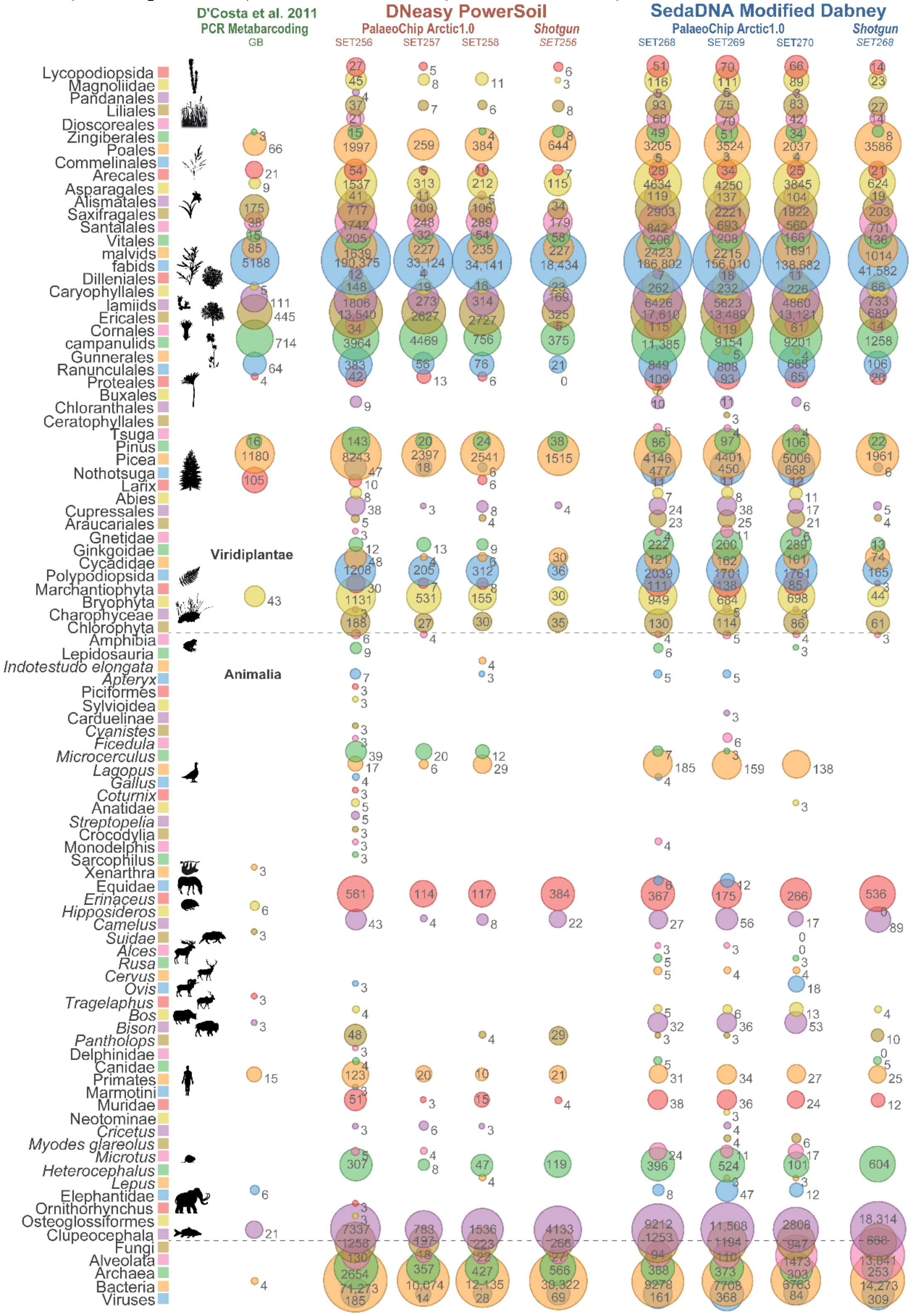

Core slice dated to 9,685 cal-BP (Mahony, 2015; Sadoway, 2014). Values indicate total reads assigned to that taxon node for Animalia, and a clade summation of reads for Viridiplantae.

**Figure S20** Metagenomic comparison of Lucky Lady II permafrost core LLII-12-84-3, all reads (not map-filtered), absolute counts (non-normalized).

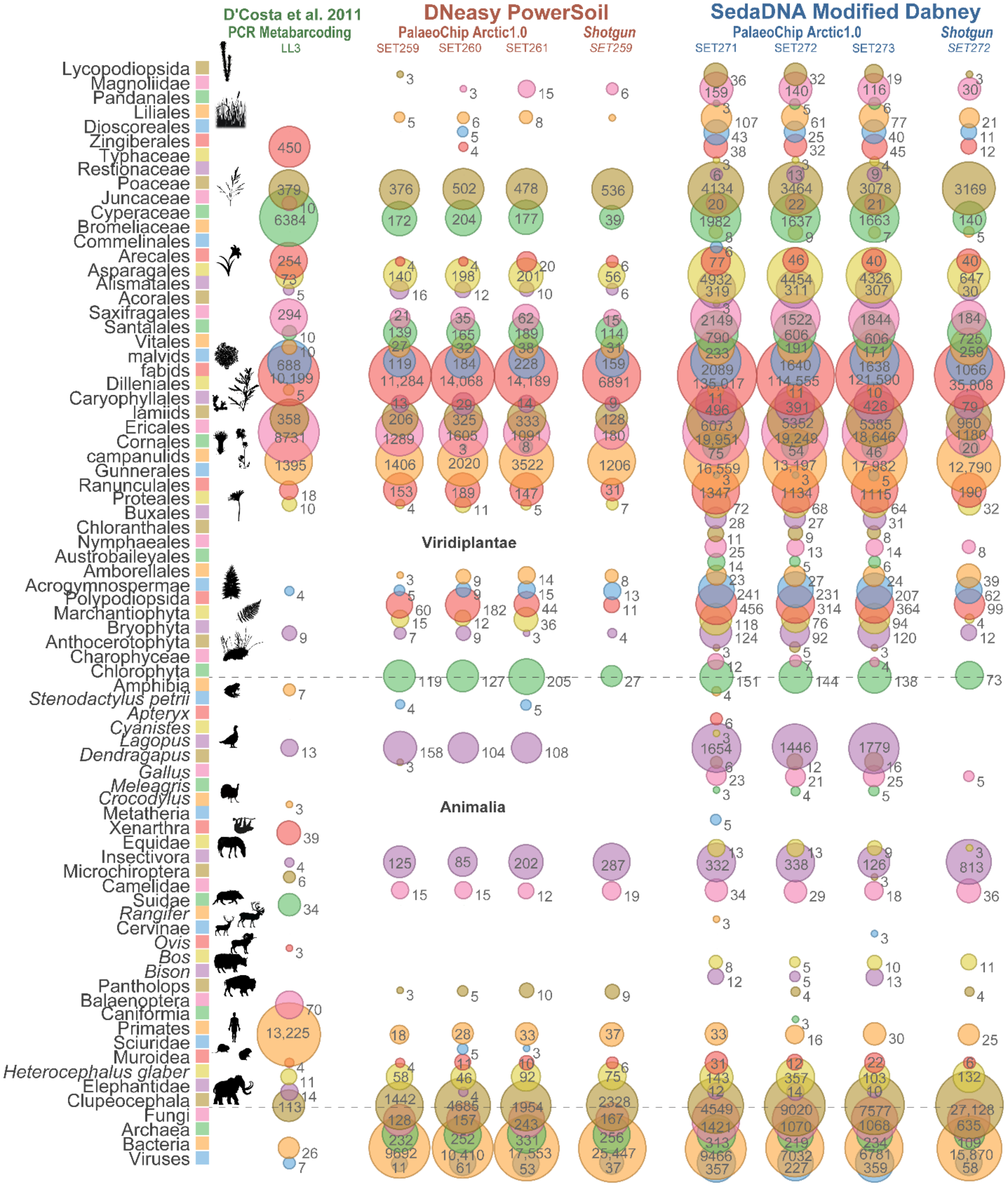

Core slice dated to 13,205 cal-BP (Sadoway, 2014). Values indicate total reads assigned to that taxon node for Animalia, and a clade summation of reads for Viridiplantae.

**Figure S21** Metagenomic comparison of Lucky Lady II permafrost core LLII-12-217-8, all reads (not map-filtered), absolute counts (non-normalized).

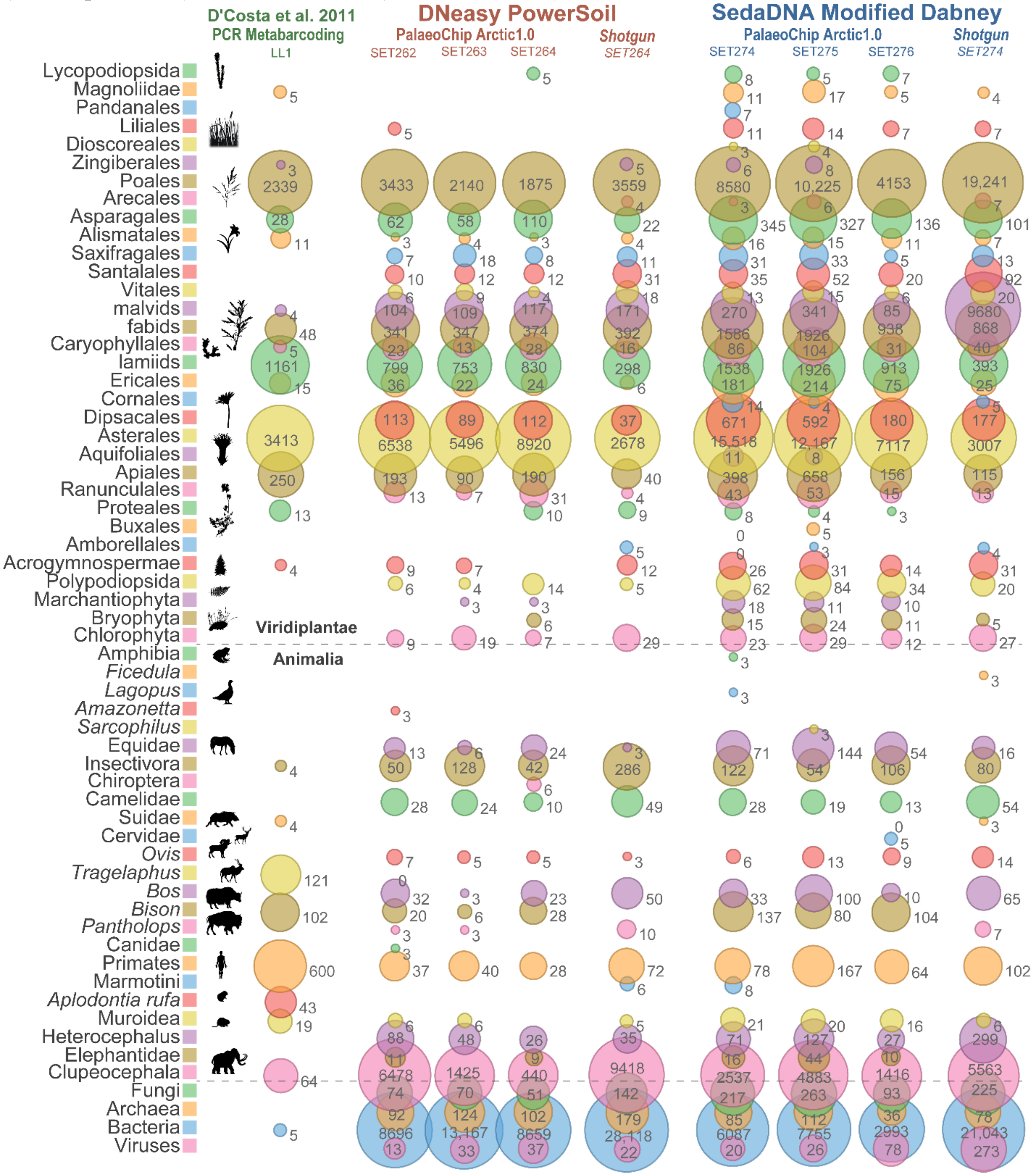

Core slice dated to 15,865 cal-BP (Sadoway, 2014). Values indicate total reads assigned to that taxon node for Animalia, and a clade summation of reads for Viridiplantae.

**Figure S22** Metagenomic comparison of Bear Creek permafrost core BC 4-2B, all reads (not map-filtered), absolute counts (non-normalized).

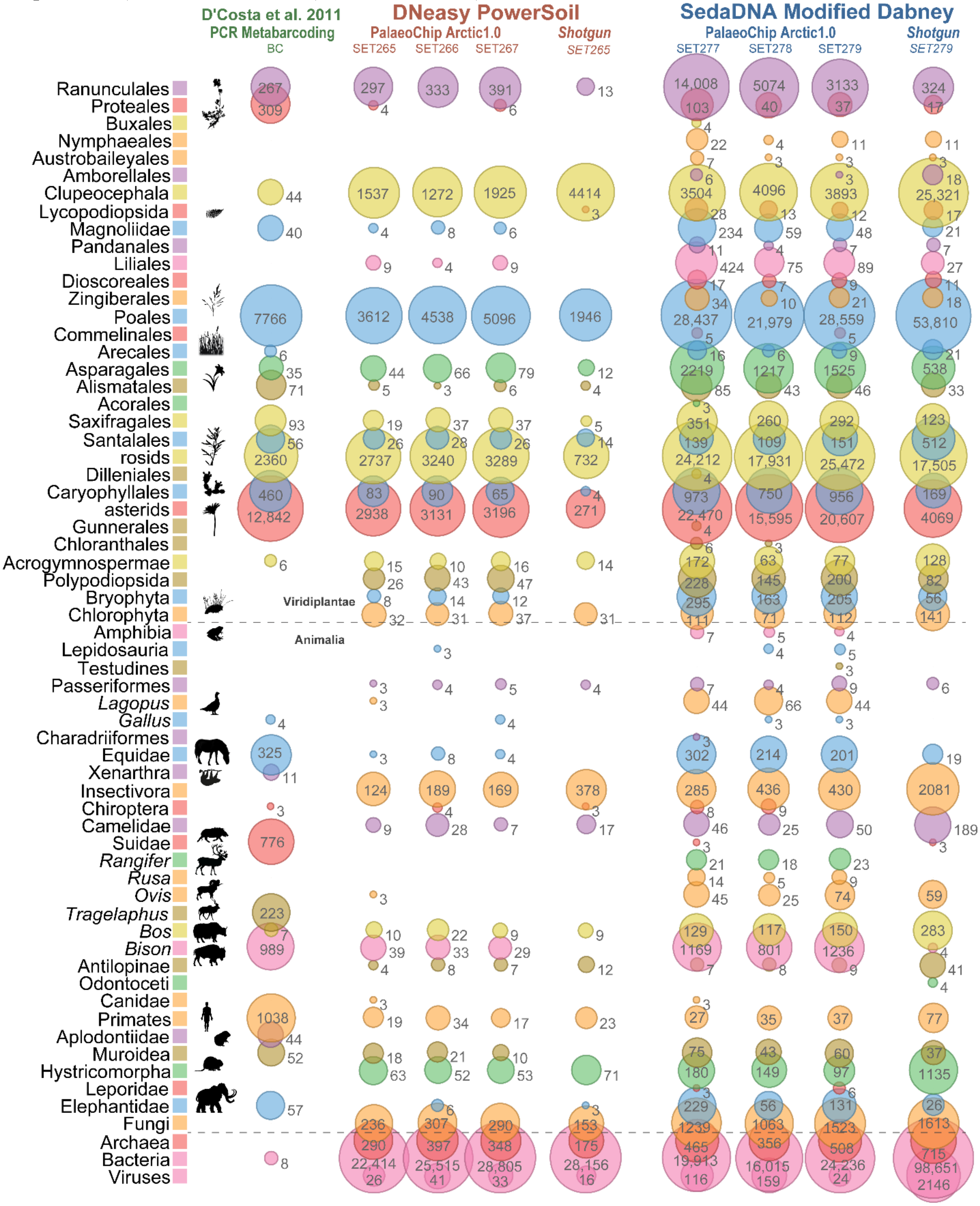

Core slice dated to ~30,000 cal-BP (D'Costa et al., 2011; Mahony, 2015; Sadoway, 2014). Values indicate total reads assigned to that taxon node for Animalia, and a clade summation of reads for Viridiplantae.

**Figure S23** Metagenomic comparison of extraction and library blanks, all reads (not map-filtered), absolute counts (non-normalized).

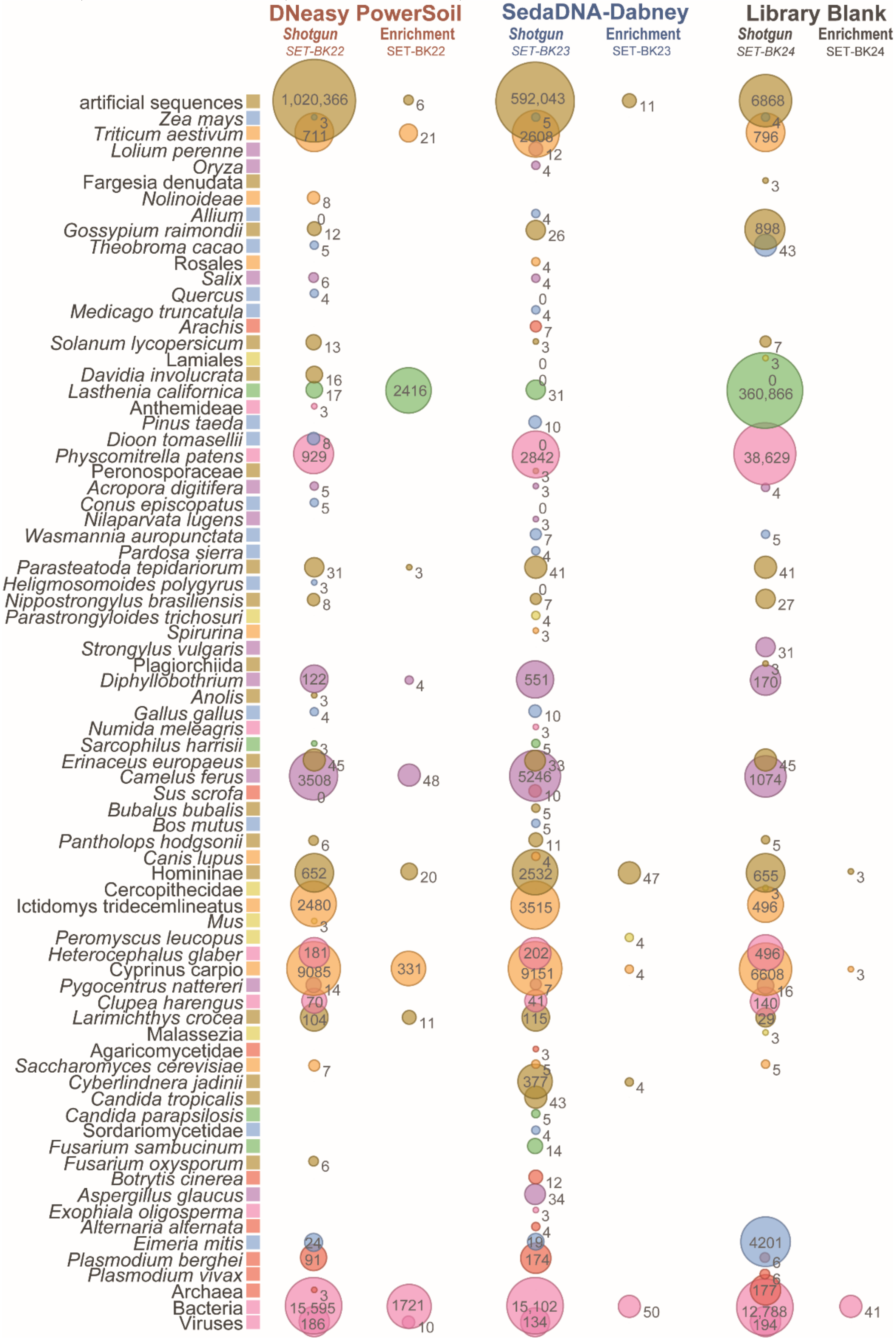

Values indicate total reads assigned to that taxon node.

**Figure S24** Metagenomic comparison of Upper Goldbottom permafrost core MM12-118b, all reads (not map-filtered), normalized counts.

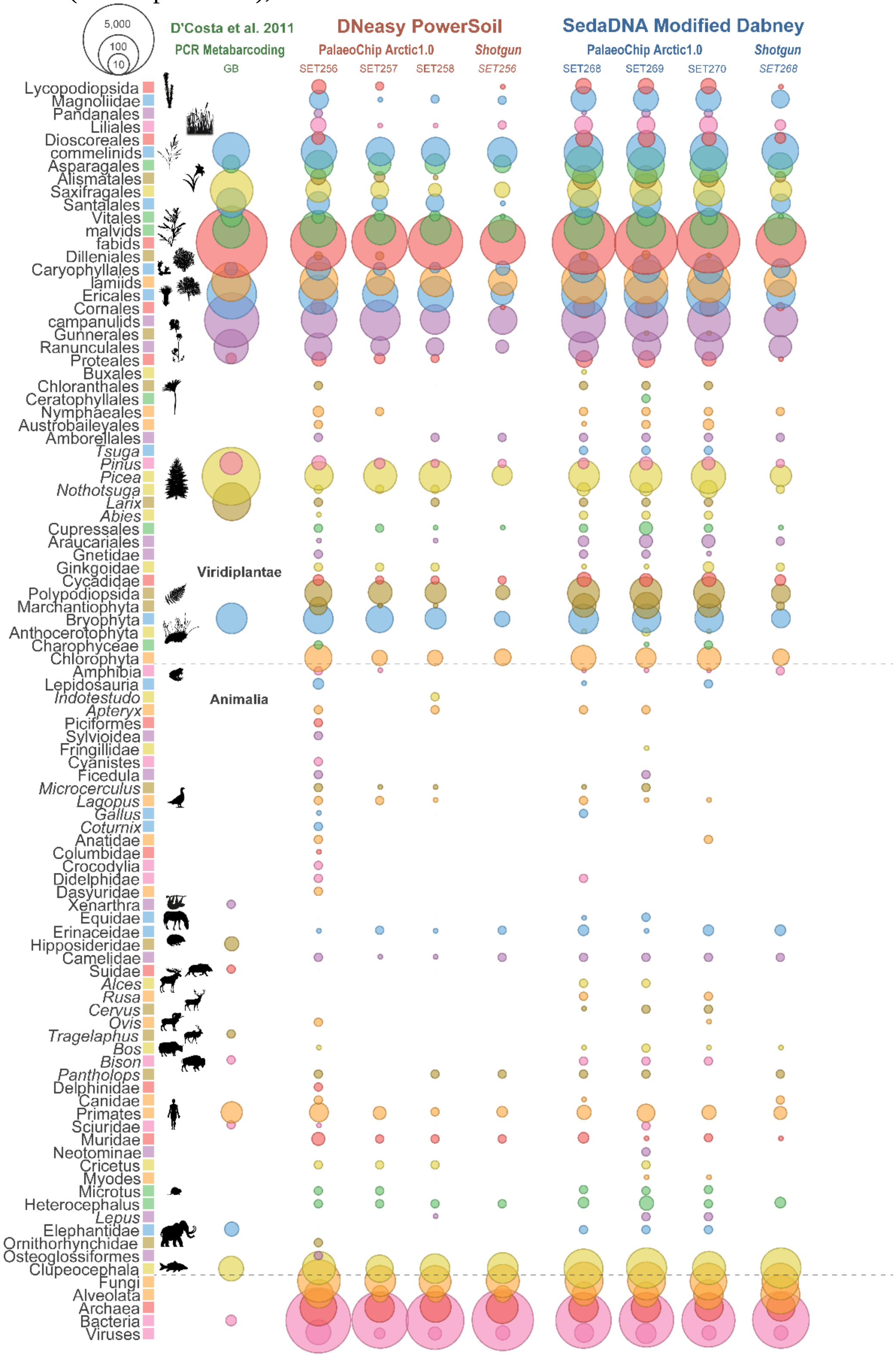

Core slice dated to 9,685 cal-BP (Mahony, 2015; Sadoway, 2014). Values indicate total reads assigned to that taxon node for Animalia, and a clade summation of reads for Viridiplantae.

602 **Figure S25** Metagenomic comparison of Lucky Lady II permafrost core LLII-12-84-3, all reads
603 (not map-filtered), normalized counts.

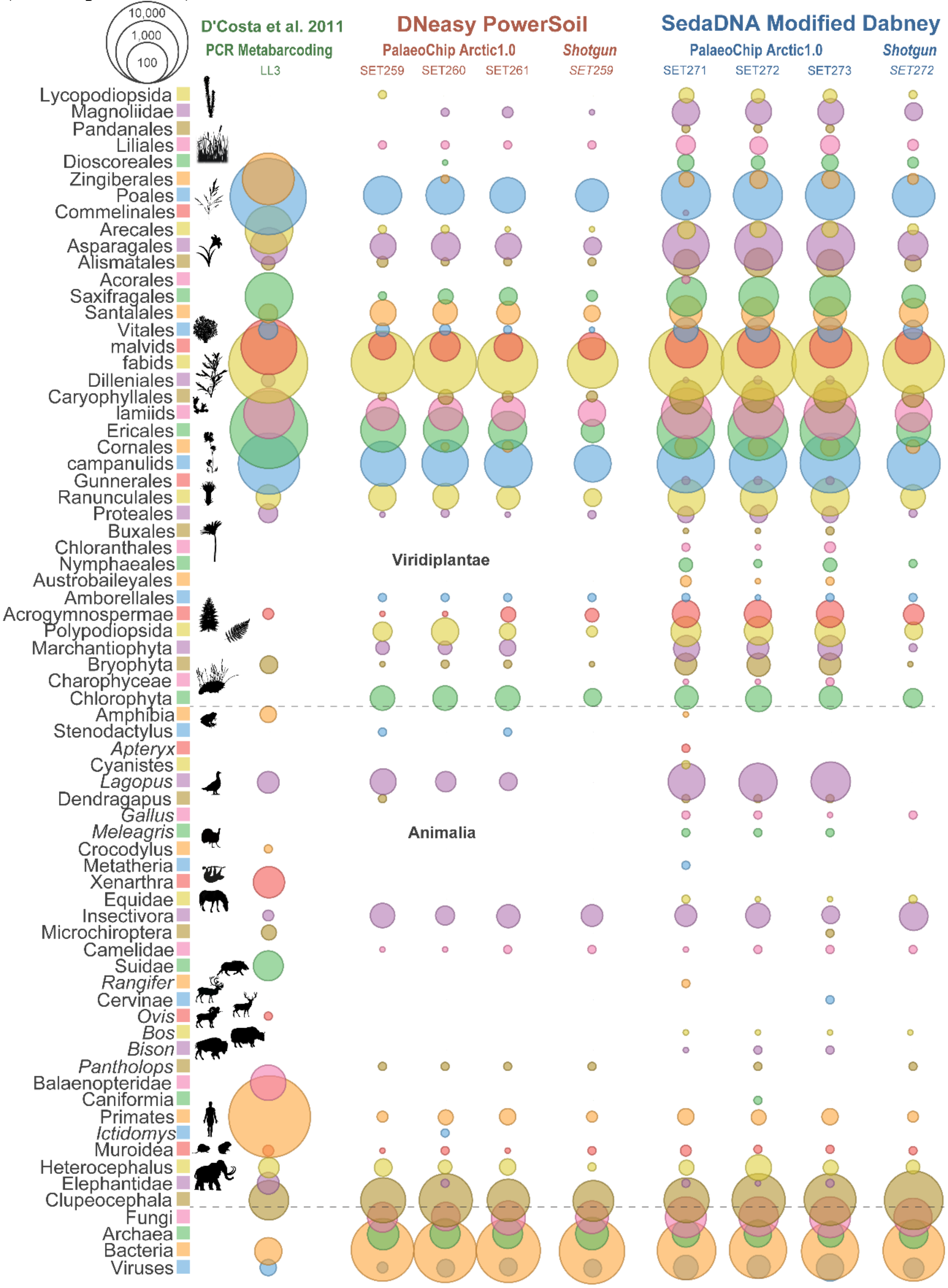

604
605 Core slice dated to 13,205 cal-BP (Sadoway, 2014). Values indicate total reads assigned to that
606 taxon node for Animalia, and a clade summation of reads for Viridiplantae.
607

**Figure S26** Metagenomic comparison of Lucky Lady II permafrost core LLII-12-217-8, all reads (not map-filtered), normalized counts.

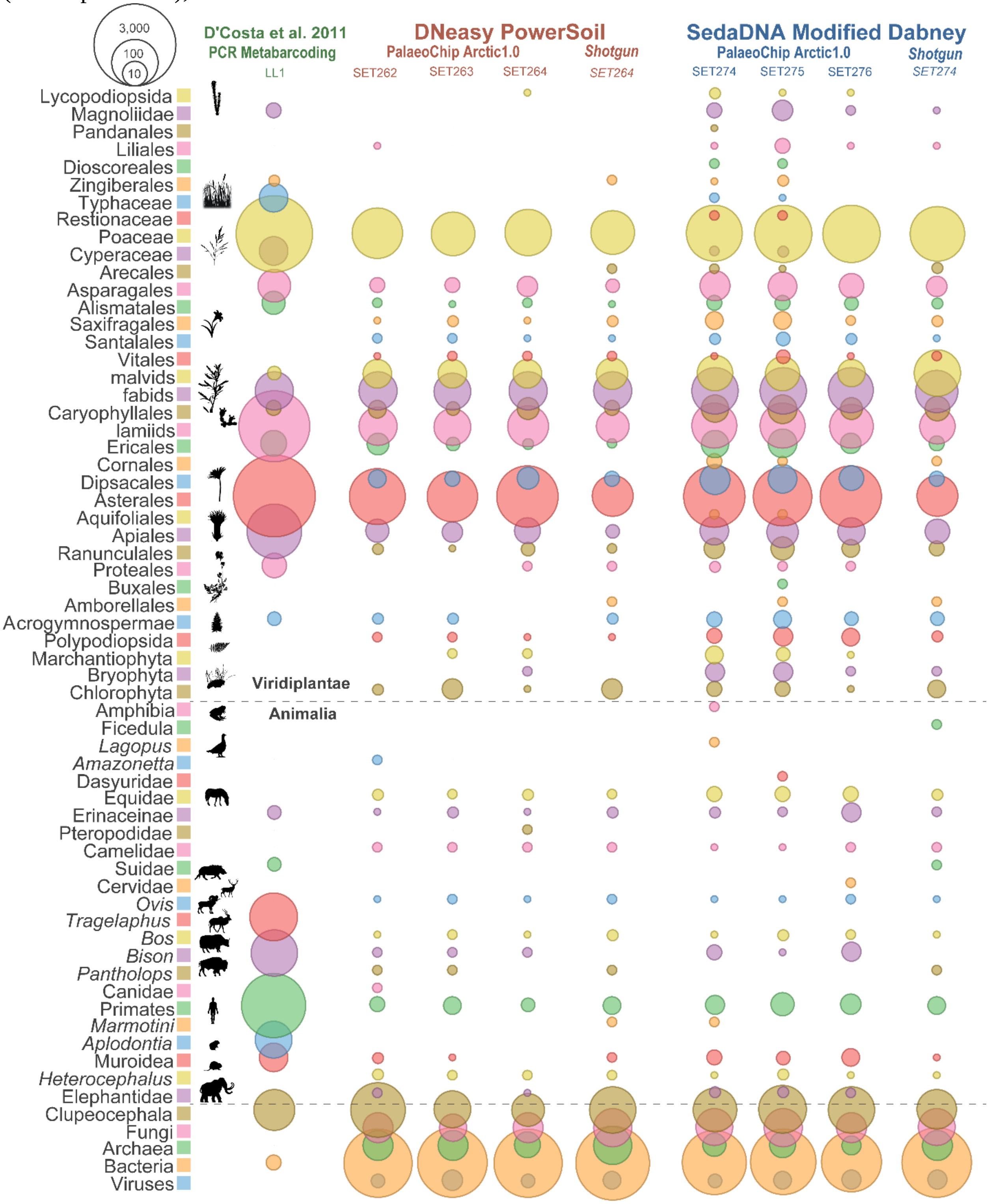

Core slice dated to 15,865 cal-BP (Sadoway, 2014). Values indicate total reads assigned to that taxon node for Animalia, and a clade summation of reads for Viridiplantae.

**Figure S27** Metagenomic comparison of Bear Creek permafrost core BC 4-2B, all reads (not map-filtered), normalized counts.

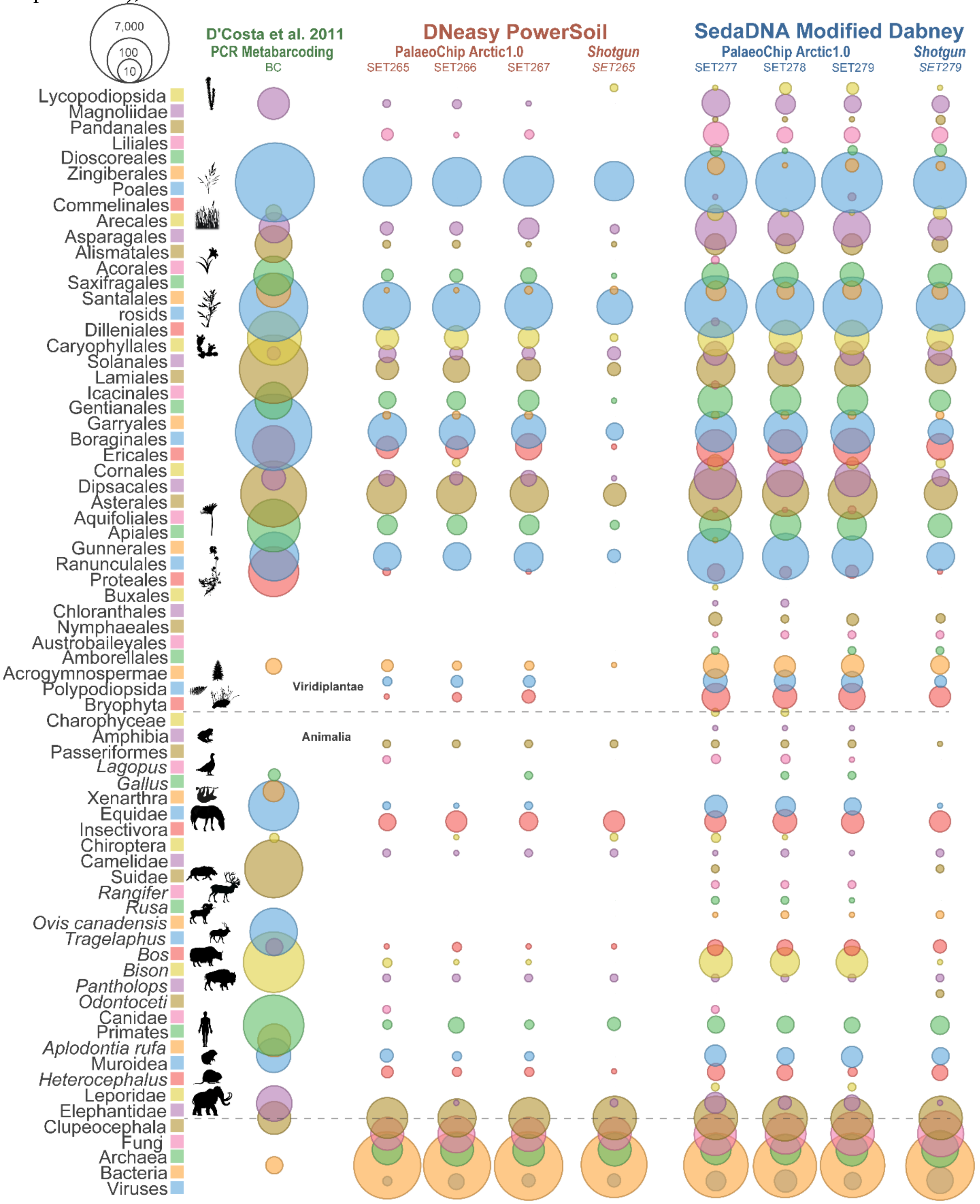

Core slice dated to ~30,000 cal-BP (D'Costa et al., 2011; Mahony, 2015; Sadoway, 2014). Values indicate total reads assigned to that taxon node for Animalia, and a clade summation of reads for Viridiplantae.

### Appendix A References

Alaeddini, R., 2012. Forensic implications of PCR inhibition - A review. Forensic Sci. Int. Genet. 6, 297–305. doi:10.1016/j.fsigen.2011.08.006

Arnold, L.J., Roberts, R.G., Macphee, R.D.E., Haile, J.S., Brock, F., M??ller, P., Froese, D.G., Tikhonov, A.N., Chivas, A.R., Gilbert, M.T.P., Willerslev, E., 2011. Paper II - Dirt, dates and DNA: OSL and radiocarbon chronologies of perennially frozen sediments in Siberia, and their implications for sedimentary ancient DNA studies. Boreas 40, 417–445. doi:10.1111/j.1502-3885.2010.00181.x

Bezanilla, M., Manne, S., Laney, D.E., Lyubchenko, Y.L., Hansma, H.G., 1995. Adsorption of DNA to Mica, Silylated Mica, and Minerals: Characterization by Atomic Force Microscopy. Langmuir 11, 655–659. doi:10.1021/la00002a050

Blum, S.A.E., Lorenz, M.G., Wackernagel, W., 1997. Mechanism of retarded DNA degradation and prokaryotic origin of DNases in nonsterile soils. Syst. Appl. Microbiol. 20, 513–521. doi:10.1016/S0723-2020(97)80021-5

Cleaves, H.J., Crapster-Pregont, E., Jonsson, C.M., Jonsson, C.L., Sverjensky, D.A., Hazen, R.A., 2011. The adsorption of short single-stranded DNA oligomers to mineral surfaces. Chemosphere 83, 1560–1567. doi:10.1016/j.chemosphere.2011.01.023

Crecchio, C., Stotzky, G., 1998. Binding of DNA on humic acids: Effect of transformation of Bacillus subtilis and resistance to DNase. Soil Biol. Biochem. 30, 1061–1067.

D’Costa, V.M., King, C.E., Kalan, L., Morar, M., Sung, W.W.L., Schwarz, C., Froese, D., Zazula, G., Calmels, F., Debruyne, R., Golding, G.B., Poinar, H.N., Wright, G.D., 2011. Antibiotic resistance is ancient. Nature 477, 457–461. doi:10.1038/nature10388

Dabney, J., Knapp, M., Glocke, I., Gansauge, M.-T., Weihmann, A., Nickel, B., Valdiosera, C., Garcia, N., Paabo, S., Arsuaga, J.-L., Meyer, M., 2013. Complete mitochondrial genome sequence of a Middle Pleistocene cave bear reconstructed from ultrashort DNA fragments. Proc. Natl. Acad. Sci. 110, 15758–15763. doi:10.1073/pnas.1314445110

Enk, J., Devault, A., Widga, C., Saunders, J., Szpak, P., Southon, J., Rouillard, J.-M., Shapiro, B., Golding, G.B., Zazula, G., Froese, D., Fisher, D.C., Macphee, R.D.E., Poinar, H., 2016. Mammuthus population dynamics in Late Pleistocene North America: Divergence, Phylogeogrpaphy and Introgression. Front. Ecol. Evol. 4, 1–13. doi:10.3389/fevo.2016.00042

Greaves, M.P., Wilson, M.J., 1970. The degradation of nucleic acids and montmorillonite-nucleic-acid complexes by soil microorganisms. Soil Biol. Biochem. 2, 257–268. doi:10.1016/0038-0717(70)90032-5

Greaves, M.P., Wilson, M.J., 1969. The adsorption of nucleic acids by montmorillonite. Soil Biol. Biochem. 1, 317–323. doi:10.1016/0038-0717(69)90014-5

Haile, J., 2008. Ancient DNA from Sediments and Associated Remains. University of Oxford.

Karpinski, E., Mead, J.I., Poinar, H.N., 2016. Molecular identification of paleofeces from Bechan Cave, southeastern Utah, USA. Quat. Int. 443, 140–146. doi:10.1016/j.quaint.2017.03.068

Khanna, M., Yoder, M., Calamai, L., Stotzky, G., 2005. X-ray diffractometry and electron microscopy of DNA from Bacillus subtilis bound on clay minerals. Sci. Soils 3, 1–10. doi:10.1007/s10112-998-0001-3

Kircher, M., Sawyer, S., Meyer, M., 2012. Double indexing overcomes inaccuracies in multiplex

sequencing on the Illumina platform. Nucleic Acids Res. 40, 1–8. doi:10.1093/nar/gkr771
Koopal, L.K., Goloub, T.P., Davis, T.A., 2004. Binding of ionic surfactants to purified humic acid. J. Colloid Interface Sci. 275, 360–367. doi:10.1016/j.jcis.2004.02.061
Lorenz, M.G., Wackernagel, W., 1987. Adsorption of DNA to sand and variable degradation rates of adsorbed DNA. Appl. Environ. Microbiol. 53, 2948–2952.
Mahony, M.E., 2015. 50,000 years of paleoenvironmental change recorded in meteoric waters and coeval paleoecological and cryostratigraphic indicators from the Klondike goldfields, Yukon, Canada. University of Alberta. doi:10.1145/3132847.3132886
Meyer, M., Kircher, M., 2010. Illumina sequencing library preparation for highly multiplexed target capture and sequencing. Cold Spring Harb. Protoc. 5. doi:10.1101/pdb.prot5448
Ogram, A., Sayler, G., Gustin, D., Lewis, R., 1988. DNA adsorption to soils and sediments. Environ. Sci. Technol. 22, 982–984.
Otto, W.H., Britten, D.J., Larive, C.K., 2003. NMR diffusion analysis of surfactant-humic substance interactions. J. Colloid Interface Sci. 261, 508–513. doi:10.1016/S0021-9797(03)00062-6
Poinar, H.N., Hofreiter, M., Spaulding, W.G., Martin, P.S., Stankiewicz, B.A., Bland, H., Evershed, R.P., Possnert, G., P????bo, S., 1998. Molecular coproscopy: Dung and diet of the extinct ground sloth Nothrotheriops shastensis. Science (80-. ). 281, 402–406. doi:10.1126/science.281.5375.402
Sadoway, T.R., 2014. A Metagenomic Analysis of Ancient Sedimentary DNA Across the Pleistocene-Holocene Transition. McMaster University.
Schlager, B., Straessle, A., Hafen, E., 2012. Use of anionic denaturing detergents to purify insoluble proteins after overexpression. BMC Biotechnol. 12. doi:10.1186/1472-6750-12-95
Shaban, I.S., Mikulaj, V., 1998. Impact of an Anionic Surfactant Addition on Solubility of Humic Acid in Acid-Alkaline Solutions. Chem. Pap. 52, 753–755.
Sidstedt, M., Jansson, L., Nilsson, E., Noppa, L., Forsman, M., Rådström, P., Hedman, J., 2015. Humic substances cause fluorescence inhibition in real-time polymerase chain reaction. Anal. Biochem. 487, 30–37. doi:10.1016/j.ab.2015.07.002
Taberlet, P., Coissac, E., Pompanon, F., Gielly, L., Miquel, C., Valentini, A., Vermat, T., Corthier, G., Brochmann, C., Willerslev, E., 2007. Power and limitations of the chloroplast trnL (UAA) intron for plant DNA barcoding. Nucleic Acids Res. 35, e14. doi:10.1093/nar/gkl938
Tanford, C., 1980. The hydrophobic effect: formation of micelles and biological membranes, 2nd editio. ed. Wiley Interscience, New York.
Taylor, B.R., Parkinson, D., 1988. Patterns of water absorption and leaching in pine and aspen leaf litter. Soil Biol. Biochem. 20, 257–258. doi:10.1016/0038-0717(88)90047-8
